# Molecular basis for short-chain thioester hydrolysis by acyl hydrolase domains in *trans*-acyltransferase polyketide synthases

**DOI:** 10.1101/2023.08.11.552765

**Authors:** Christopher D. Fage, Munro Passmore, Ben P. Tatman, Helen G. Smith, Xinyun Jian, Upeksha C. Dissanayake, G. Andrés Cisneros, Gregory L. Challis, Józef R. Lewandowski, Matthew Jenner

## Abstract

Polyketide synthases (PKSs) are multi-domain enzymatic assembly lines that biosynthesise a wide selection of bioactive natural products from simple building blocks. In contrast to their *cis*-acyltransferase (AT) counterparts, *trans*-AT PKSs rely on stand-alone AT domains to load extender units onto acyl carrier protein (ACP) domains embedded in the core PKS machinery. *Trans*-AT PKS gene clusters also encode acyl hydrolase (AH) domains, which are predicted to share the overall fold of AT domains, but hydrolyse aberrant acyl chains from ACP domains, thus ensuring efficient polyketide biosynthesis. How such domains specifically target short acyl chains, in particular acetyl groups, tethered as thioesters to the substrate-shuttling ACP domains, with hydrolytic rather than acyl transfer activity, has remained unclear. To answer these questions, we solved the first structure of an AH domain and performed structure-guided activity assays on active site variants. Our results offer key insights into chain length control and selection against coenzyme A-tethered substrates, and clarify how the interaction interface between AH and ACP domains contributes to recognition of cognate and non-cognate ACP domains. Combining our experimental findings with molecular dynamics simulations allowed for the production of a data-driven model of an AH:ACP domain complex. Our results advance the currently incomplete understanding of polyketide biosynthesis by *trans*-AT PKSs, and provide foundations for future bioengineering efforts.

## INTRODUCTION

Type I modular polyketide synthases (PKSs) are impressive molecular machines responsible for the construction of complex, bioactive natural products that find numerous applications in both medicine and agriculture.^1,2^ Often likened to ‘molecular assembly lines’, these multi-domain enzymes conform to a paradigm of covalent substrate attachment for exceptional processivity. Attachment occurs at the free thiol of a co-enzyme A (CoA)-derived phosphopantetheine (Ppant) moiety that is post-translationally appended to *apo*-acyl carrier protein (ACP) domains by a phosphopantetheinyl transferase (PPTase) enzyme. Whilst tethered to this thiol, acyl groups are efficiently shuttled between active sites of catalytic domains, making ACP domains central to modular PKSs. Using enzymology akin to that of fatty acid synthases, a minimal PKS chain extension module consists of an acyltransferase (AT) domain that loads an (alkyl)malonyl-derived extender unit onto the ACP domain, and a ketosynthase (KS) domain that catalyses a Claisen-like decarboxylative condensation between the extender unit and the upstream polyketide chain to yield a β-keto-thioester intermediate. The optional presence of α/β-carbon-modifying domains within modules, such as ketoreductase (KR), dehydratase (DH), enoylreductase (ER) and methyltransferase (MT) domains, alter the resulting β-keto-thioester to further enhance chemical diversity.^3–5^

Evolution has given rise to two phylogenetically distinct classes of modular PKSs: *cis*-AT and *trans*-AT. In stark contrast to the ‘textbook’ *cis*-AT PKSs, AT domains are absent from the chain extension modules of *trans*-AT assembly lines. Instead, the loading of extender units onto ACP domains is catalysed by stand-alone AT domains encoded by distinct genes elsewhere in the biosynthetic gene cluster (BGC).^6^ Interestingly, many *trans*-AT PKS BGCs also encode a *trans*-acting AT-like domain possessing hydrolytic activity towards ACP-bound thioesters. These enzymes, subsequently classified as acyl hydrolase (AH) domains, are distinct from AT domains at the sequence level and form a separate phylogenetic clade.^7^ Although AH domains were originally hypothesised to ‘proofread’ PKSs by hydrolysing any stalled intermediates from ACP domains, subsequent analysis of their substrate specificity showed a clear preference for short, linear acyl chains attached to ACP domains.^8^ This suggested a more precise housekeeping role for AH domains: namely, the removal of unwanted acetyl groups from ACP domains installed by erroneous PPTase-catalysed transfer from acetyl-CoA, or by spontaneous decarboxylation of malonyl-ACP domain thioesters (**Fig. 1A**). Such a role is analogous to that played by type II thioesterase (TE_II_) domains in *cis*-AT PKSs, which have been shown to similarly prefer short acyl chains attached to ACP domains.^9,10^ Nevertheless, TE_II_ domains are fundamentally divergent from AT domains and possess a single subdomain (instead of two) with a catalytic triad (instead of a dyad). These features highlight the disparate mechanisms that PKSs have evolved to solve key biochemical problems.

**Figure 1.**
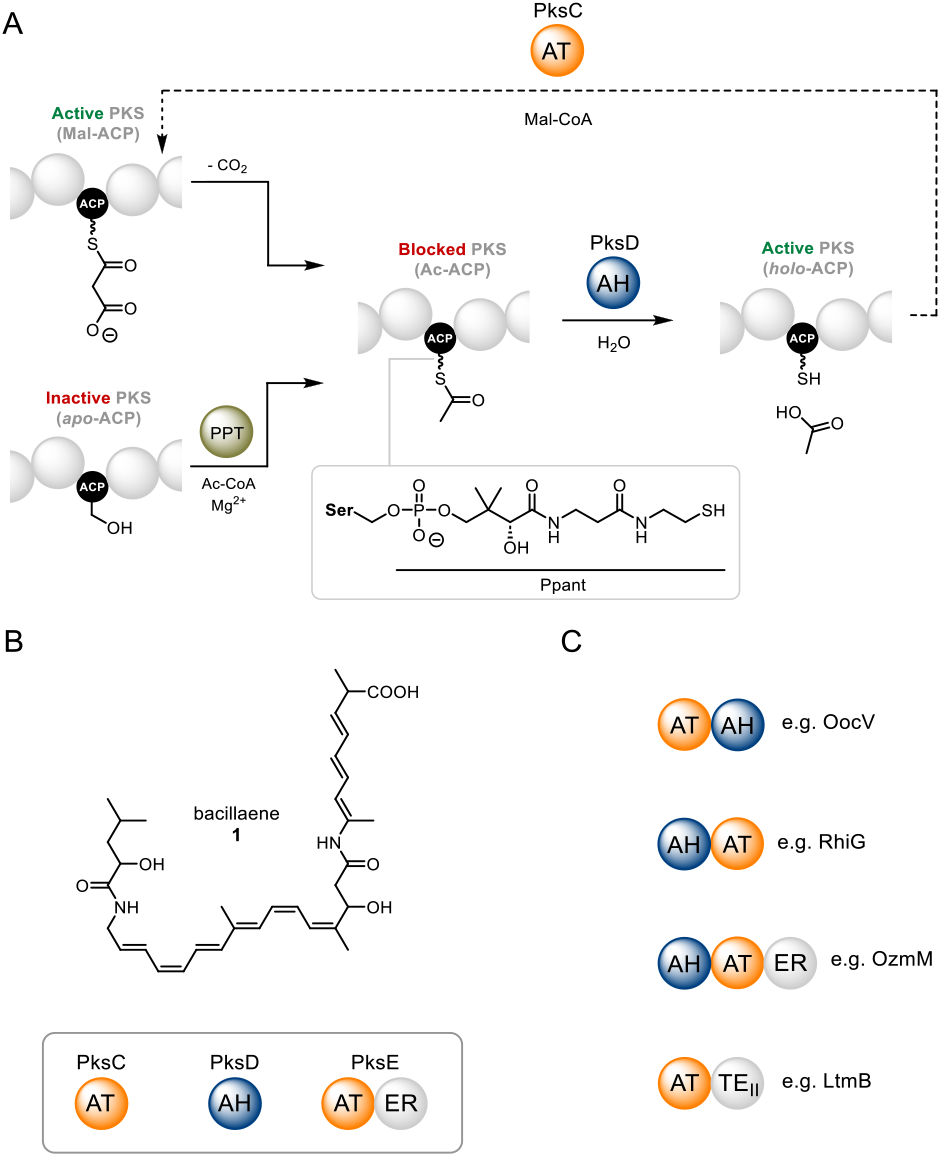
Catalytic roles of AH and AT domains and their occurrence in *trans*-AT PKSs. **A**). Proofreading and extender unit selection in the bacillaene *trans*-AT PKS. Spontaneous decarboxylation of malonyl extender units or erroneous loading of acetyl-CoA by a PPTase (PPT; left) results in a stalled acetyl-ACP species, which blocks the PKS pathway (center). PksD catalyses hydrolysis of the acetyl group to give a *holo*-ACP species (right), which can then be malonylated through PksC activity (top). **B**). Chemical structure of the bacillaene product of the PksX pathway, with employed AT and AH domains shown. **C**). Examples of known AT and AH domain fusions in proteins from *trans*-AT PKS pathways.

Although *trans*-acting AT and AH domains can be found as stand-alone proteins, for example PksC (AT) and PksD (AH) from the bacillaene PKS (**Fig. 1B**), they can also occur as di- or tri-domain fusions with AT-AH (e.g. OocV), AH-AT (e.g. RhiG), AT-ER (e.g. PksE) or AH-AT-ER (e.g. OzmM) architectures (**Fig. 1C**).^6^ Interestingly, AT domains, but not AH domains, can pair with a C-terminal ER domain, possibly as a result of favourable gene fusion events. It is worth noting that the closely related lactimidomycin and migrastatin BGCs possess a *trans*-acting AT-TE_II_ di-domain (LtmB/MgsB)^11^ – the only known example of a hybrid protein containing elements of both *cis*-AT and *trans*-AT PKSs (**Fig. 1C**). Recent studies have afforded valuable insights into the structure and mechanism of *trans*-acting AT domains. For example, the disorazol AT domain, DisD, exhibits an α/β-hydrolase fold with two subdomains,^12^ similar to its *cis*-acting relatives.^13–16^ It is worth noting that, AT domains from *cis*-AT PKSs are often crystallised as covalently tethered KS-AT di-domains,^17,18^ whilst only a remnant of the KS-AT linker region (or ‘flanking subdomain’) can be found appended to the KS domains from *trans*-AT PKSs.^19,20^ Whereas the majority of *trans*-acting AT domains use malonyl-CoA extender units, examples of alkylmalonyl-CoA extender unit utilisation have been observed.^21–23^ Specificity towards malonyl extender units is achieved through conserved GH**<u>S</u>**xGE and xF**<u>H</u>**S motifs (catalytic dyad residues highlighted), where the xF**<u>H</u>**S motif’s Phe residue is positioned to preclude α-substituted extender units.^24^ In addition, a recently reported crystal structure in which DisD was crosslinked to a cognate ACP domain yielded residue-level insight into the protein-protein interactions governing malonyl transfer, in close agreement with previously published alanine scanning mutagenesis results.^12,25^

Bioinformatics-based prediction of the AH domain structure suggests an α/β-hydrolase fold as observed for AT domains; however, the mechanism by which the same overall fold catalyses chain length-controlled hydrolysis of ACP-bound acyl groups has remained unclear. Herein, we report the first structure of an AH domain from a *trans*-AT PKS, which, when compared to AT domain counterparts, provides insights into mechanism and substrate specificity. Roles for key residues are further supported by intact protein-MS assays of AH domain variants. In addition, the molecular basis for interactions between AH and ACP domains is illuminated through alanine scanning mutagenesis, mass spectrometry (MS)-based carbene footprinting and molecular dynamics (MD) simulations. Finally, AH activity is demonstrated on a panel of carrier protein domains from other biosynthetic systems, highlighting the utility of AH domains for dissecting machineries that depend on carrier protein domains to shuttle intermediates. Our study adds to the emerging understanding of the unique catalytic strategies employed by *trans*-AT PKSs to assemble diverse bioactive compounds.

## RESULTS AND DISCUSSION

### Structural analysis of the PksD domain

To gain insight into the differential catalytic roles of AH and AT domains in spite of a putatively shared fold, we pursued a crystal structure of a representative AH domain. PksD, the AH domain from the bacillaene pathway (**Fig. 1B**), was a reasonable starting point as PksC, the corresponding AT domain from the same pathway, had been previously crystallised and characterised,^26,27^ providing an excellent point of comparison. The *pksD* gene from *Bacillus subtilis* str. 168 was cloned and the corresponding protein was overproduced in *Escherichia coli* as an N-terminal His_8_-tagged fusion protein, which was purified to homogeneity using immobilized metal-ion affinity chromatography and size-exclusion chromatography (**Table S1**). The identity of the purified protein was confirmed by ESI-Q-TOF-MS analysis (**Fig. S1**). Crystallographic screens and subsequent optimisations of both native and selenomethionine (SeMet)-derived PksD, including streak seeding for the latter,^28^ yielded diffraction-quality crystals (**Fig. S2A, Tables S2** and **S3**). A single-wavelength anomalous dispersion dataset, diffracting to ∼1.96 Å from a SeMet-derived crystal, revealed four chains per asymmetric unit in a *C*2 unit cell (**Table S4**). One of these chains was employed as a molecular replacement search model to phase reflections from a native crystal, which diffracted to ∼2.20 Å. Eight native PksD chains were identified per asymmetric unit in a *C2* unit cell with significantly different parameters than the SeMet derivative (**Table S4**). Notwithstanding discrepancies in crystal packing, SeMet-derived and native PksD chains superpose very well (**Fig. S2B**), and by virtue of the higher resolution of the anomalous dataset, all structural analyses were performed on the SeMet-derived model.

Structural evaluation of DisD (AT) (PDB: 5ZK4),^25^ PksC (AT) (PDB: 5DZ6) and PksD (AH) (PDB: 8AVZ) reveals very similar topologies, albeit with significant deviations in the relative positions of secondary structure elements between the two *trans*-acting domain types (compare **Fig. 2A** and **2B**). Conspicuously, a latch structure consisting of residues Ile287-Arg324 (maroon in **Fig. 2A**) is appended to helix α15 of PksD that is absent from the AT domains. This feature supplies an additional 3_10_-helix, β-strand, α-helix and intervening loops that account for ∼12% of the mass of PksD (**Fig. S3**). In *cis*-acting AT domains, such as that from the third module of the 6-deoxyerythronolide B PKS (PDB: 2QO3),^29^ the latch furnishes an interface between the flanking subdomain and the AT domain, with a C-terminal loop tracing the surface of the KS domain (**Fig. 2C** and **2D**). Interestingly, the latch region is present in many structures of AT domains from both primary and secondary metabolism, with variations in length (20-36 residues) arising from truncations of the unstructured loops (**Fig. S4**).

**Figure 2.**
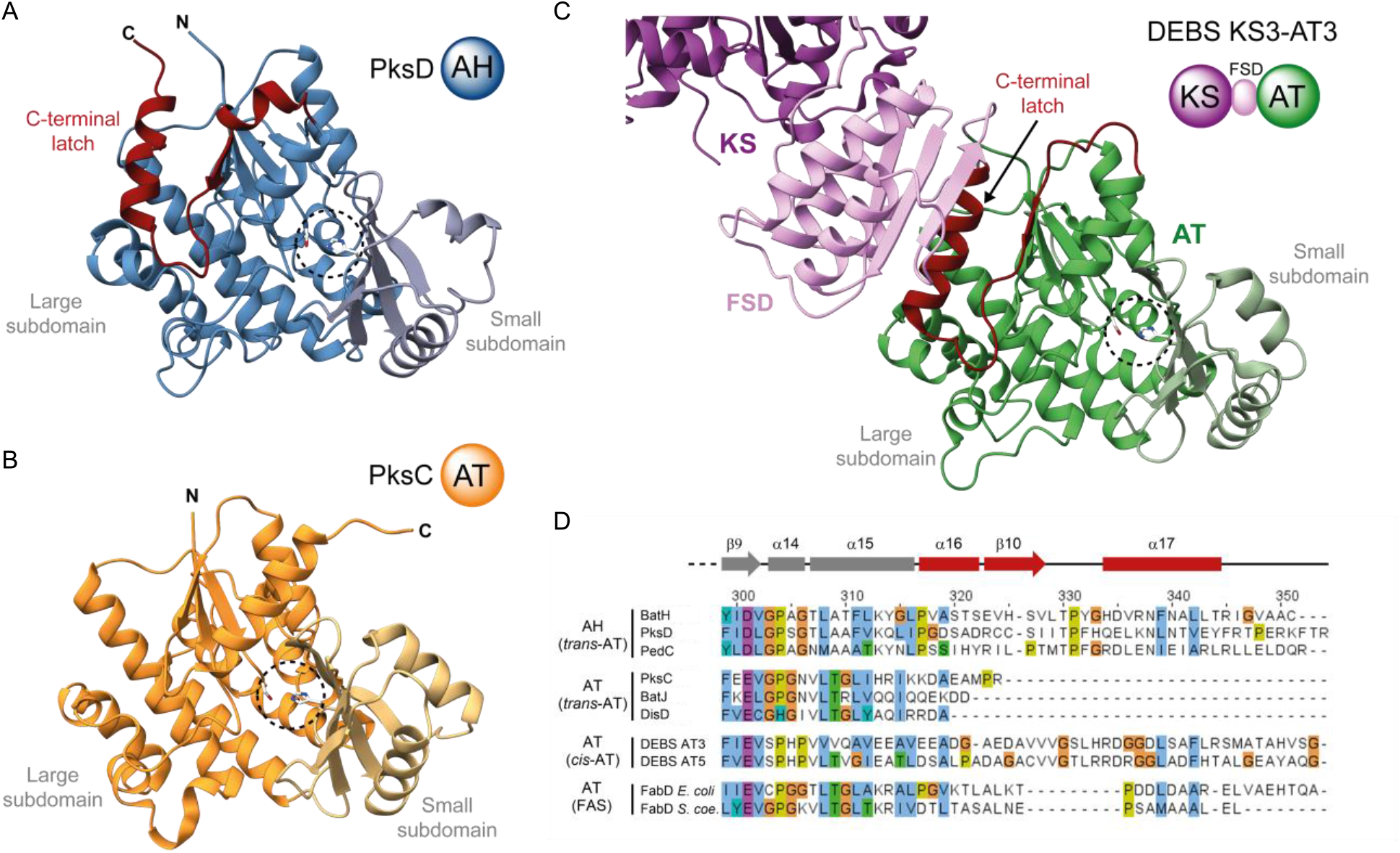
Structure of PksD and comparison to AT domains involved in polyketide biosynthesis. Catalytic residues are highlighted as sticks, and the small ferredoxin-like subdomain is coloured in a lighter shade than the large subdomain. Structural comparison of **A**). PksD (AH) (PDB: 8AVZ) and **B**). PksC (AT) (PDB: 5DZ6) from the bacillaene *trans*-AT PKS. The overall topology of both domains is highly similar, apart from the C-terminal latch appended to PksD (highlighted in red). **C**). Structure of the KS-AT di-domain fragment from module 3 of 6-deoxyerythronolide B synthase (DEBS; PDB: 2QO3). In *cis*-AT PKSs like DEBS, the latch is sandwiched between the flanking subdomain (FSD) and the large subdomain of the AT domain; a C-terminal loop wraps around the FSD and packs closely to the KS domain surface. **D**). Multiple sequence alignment of AH and AT domains from *trans*-AT PKSs, *cis*-AT PKSs and bacterial type II fatty acid synthases (FASs). C-terminal sequences are shown to highlight the presence/absence of the latch. The secondary structure elements of PksD (common among the selected domains, with differing boundaries) are displayed above. Abbreviations: *S. coe.* (*Streptomyces coelicolor*).

Despite the high purity of our PksD samples, dual elution peaks were evident in the size-exclusion chromatography (**Fig. S1A** and **S5A**). Such a profile is diagnostic of a monomer-dimer equilibrium, and we indeed identified a limited interface between pairs of PksD monomers in the crystal structure. At the centre of the interface, we observed significant but unmodeled electron density suggestive of a heavy atom. The occupying atom is symmetrically coordinated with tetrahedral geometry by Cys236 and His238 side chains from each of two monomers. X-ray fluorescence experiments were carried out to identify the coordinated atom, whereupon the presence of zinc was determined from an emission spectrum and an edge scan (**Fig. S5B** and **S5C**). Placement of a Zn^2+^ ion resulted in satisfactory refinement (**Fig. S5D**, **S5E** and **S5F**). Curiously, the observed chelation pattern resembles that of zinc fingers, a family of intramolecular motifs used primarily by DNA- and RNA-binding proteins to regulate gene expression.^30^ A similar role is deemed improbable for AH domains, especially given the lack of conservation of Cys236 and His238 (**Fig. S6**); however, Zn^2+^ ion binding is also unlikely to be an artefact of crystallisation as no zinc was included in the crystal screening and optimisation conditions. Whether PksD exists as a dimer, uniquely among its homologs, under physiological conditions remains to be determined. As discussed above, AH domains often form part of AH-AT-ER tri-domain proteins, and whilst the overall architecture of such multi-domain proteins remains unclear, *trans*-acting ER domains are known to be dimeric (e.g. the ER region of the DfnA AT-ER di-domain protein).^31^ In such instances, the AH and AT domains would be pseudo-dimeric as a consequence of the ER domain, but presumably functional despite this oligomeric state.

### Structural and functional analysis of PksD catalytic mechanism

Akin to its AT domain relatives, PksD appears to employ a Ser-His catalytic dyad (**Fig. 3A**, **3B** and **3C**). His201 is optimally positioned to deprotonate the hydroxyl group of Ser99 (< 3 Å between Nε and Oγ), thus activating it for nucleophilic attack on incoming ACP domain-bound substrates; His201 may likewise be poised to activate a water molecule for hydrolytic cleavage following acylation of Ser99 with the substrate (see our MD simulations below). *E. coli* FabD (PDB: 2G2Z), an AT domain involved in fatty acid biosynthesis, was previously crystallised in a non-covalent complex with CoA and with the nucleophilic Ser92 acylated with a malonyl group.^32^ The positively charged guanidino group of the conserved Arg117 residue forms a stabilising salt bridge with the negatively charged carboxylate group of the malonyl (**Fig. 3C**). In AH domains, however, no such stabilisation is possible as this position is occupied by Gln (Gln124 in PksD) or, less frequently, other residues (**Fig. 3A, 3D** and **Fig. S7**). Substituting the conserved Arg side chain for a shorter and uncharged one is presumably important in preventing correctly loaded extender units (e.g. malonyl and methylmalonyl groups) from being transferred to and/or hydrolysed by the AH domain. In line with these observations, previous work showed that mutating the malonyl-binding Arg to Gln in PedD, an AT from the pederin pathway, afforded acylation of the nucleophilic Ser residue with an acetyl group carried by an ACP domain, but not subsequent hydrolysis.^8^ In another distinctive feature, the putative catalytic Ser99 residue of PksD is often preceded by another Ser residue instead of the strictly conserved His residue found in AT domains (e.g. G(A/S/V)**<u>S</u>**L vs. GH**<u>S</u>**L motif; catalytic Ser residue bolded and underlined) (**Fig. 3D**). No role has been ascribed to the residue at this position, despite its strong conservation in AT domains.

**Figure 3.**
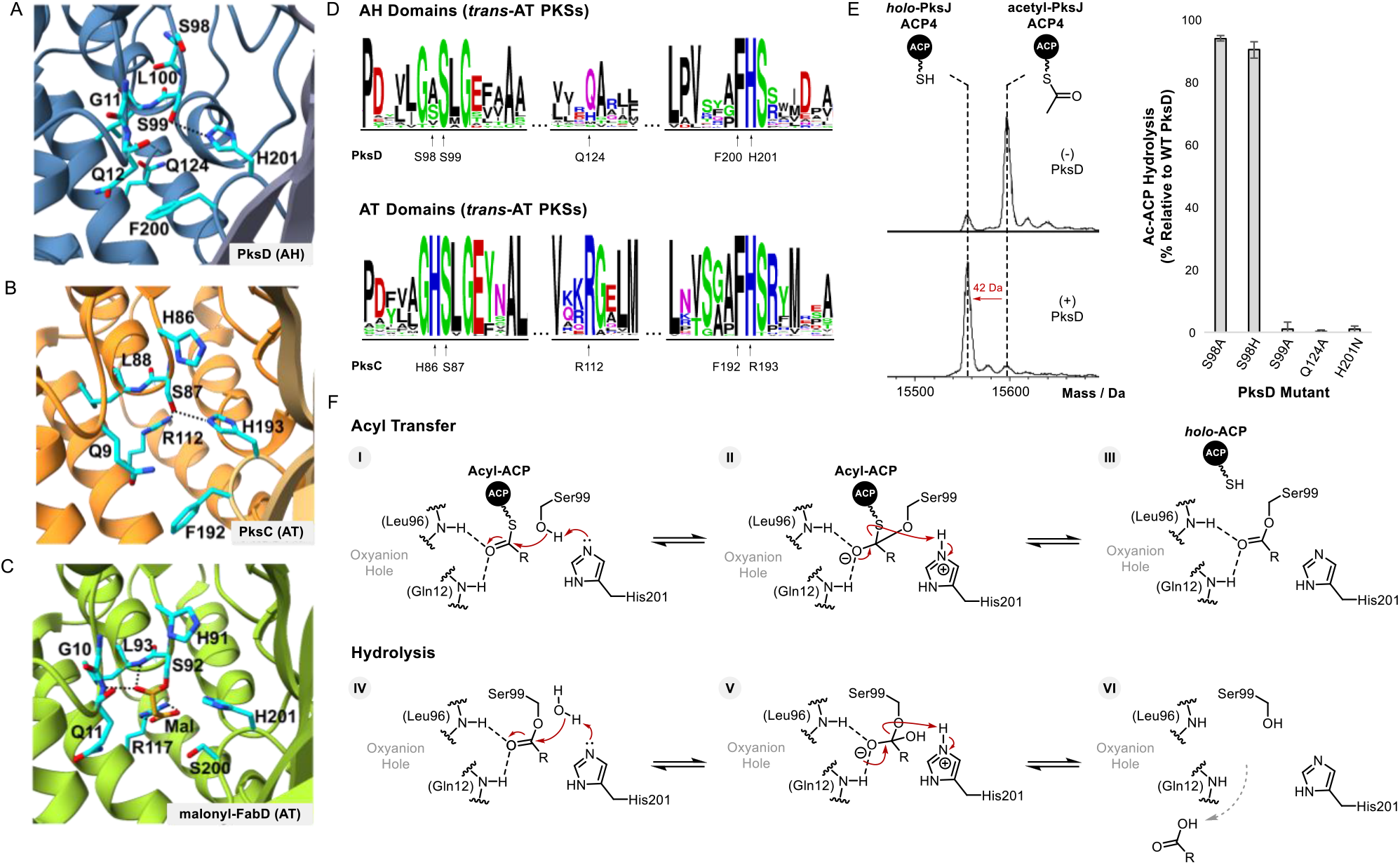
Comparison of AH and AT active sites provides insights into substrate specificity and catalytic mechanism. Direct comparison of active site residues for **A**). PksD (AH) (PDB: 8AVZ), **B**). PksC (AT) (PDB 5DZ6) and **C**). FabD (AT) (PDB: 2G2Z). The positively charged Arg residue in PksC and FabD is essential for acyl transfer and promotes binding of the malonyl group. Absence of an Arg residue at this position is diagnostic of an AH domain (usually replaced with Gln/His; see **Fig. S7**). **D**). Sequence logo comparison of active site residues in AH and AT domains from *trans*-AT PKS systems. Letter height indicates the relative degree of sequence conservation at each position. The residue type and position for PksD and PksC are shown beneath their respective sequence logos. **E**). Left: stacked, deconvoluted ESI-Q-TOF spectra of acetylated PksJ ACP4 before (*top*) and after (*bottom*) incubation with PksD. Right: effect of PksD active site mutations on hydrolytic activity, plotted relative to WT PksD. Error bars represent the standard deviation of three replicates. **F**). Proposed catalytic mechanism for PksD-catalysed hydrolysis of an ACP domain-tethered thioester. The R-group represents a short alkyl chain.

To evaluate the contributions of residues in the active site of PksD, we prepared several variants and compared their hydrolytic activities to that of the wild-type enzyme. PksJ ACP4, a cognate ACP domain of PksD from the bacillaene *trans*-AT PKS, was enzymatically loaded with an acetyl group, and then incubated with WT PksD or mutants thereof for 5 min, after which the reaction was acid-quenched and analysed by intact protein-MS. In this assay, WT PksD showed near-complete hydrolytic conversion of acetyl-PksJ ACP4 to *holo*-PksJ ACP4 after 5 min (**Fig. 3E** and **Fig. S8**). In contrast, mutation of either residue in the putative catalytic dyad (S99A and H201N) fully abolished hydrolytic cleavage of the acetyl group (**Fig. 3E**), suggesting critical roles not dissimilar from those of the equivalent pair in AT domains. Given the strict conservation of His in the nucleophilic G**<u>H</u>**SL motif of AT domains, we analysed the corresponding Ser residue in the G(A/S/V)**<u>S</u>**L motif of PksD. That S98A and S98H mutations had little effect on the enzyme’s ability to hydrolyse substrate (**Fig. 3E**) suggests a non-crucial role for the residue preceding the nucleophilic Ser in AH domains, as further supported by a lack of strict conservation at this position (**Fig. 3D**). Conversely, a Q124A mutation resulted in inactivity towards acetyl-PksJ ACP4 (**Fig. 3E**), indicating an important role for the Arg-substituting residue in either catalysis or structure (e.g. organising the geometry of the substrate binding pocket; see below).

Taken together, the AH reaction likely proceeds through an acyl-enzyme intermediate similar to that formed by AT domains during acyl transfer, but in the reverse direction (i.e. acyl transfer from the ACP domain to the AH domain; top of **Fig. 3F**). In further contrast to AT domains, which partition their acyl-enzyme intermediates down acyl transfer (productive) or hydrolysis (non-productive) pathways according to substrate viability,^33^ AH domains have evolved to productively couple acyl transfer with hydrolysis (bottom of **Fig. 3F**).

### Insights into PksD substrate specificity and chain length control

Previous work showed that PedC, the AH from the pederin pathway, preferentially hydrolyses short, unbranched acyl chains bound to ACP domains.^7,8^ To determine whether this substrate profile is conserved among AH domains, we measured PksD-catalysed hydrolysis of acetyl-, butyryl-, hexanoyl-, octanoyl-, β-hydroxybutyryl- and malonyl-PksJ ACP4 using our intact protein-MS assay. Indeed, our experiments confirmed a clear preference for short, unbranched acyl chains on PksJ ACP4, with only trace hydrolysis of the malonyl group, in line with our expectations of PksD as a housekeeping enzyme (**Fig 4A**). In addition, a computational energy decomposition analysis of acetyl- and malonyl-PksD relative to the unmodified form showed pronounced differences. Here, the total interaction energies of acetyl-PksD and the unmodified form were comparable and favourable (approx. -30 kcal/mol), whereas that of malonyl-PksD was unfavourable (approx. +94 kcal/mol), with numerous residues affected across the protein structure, indicating a suboptimal substrate (**Fig. 4B, Fig. S9**).

**Figure 4.**
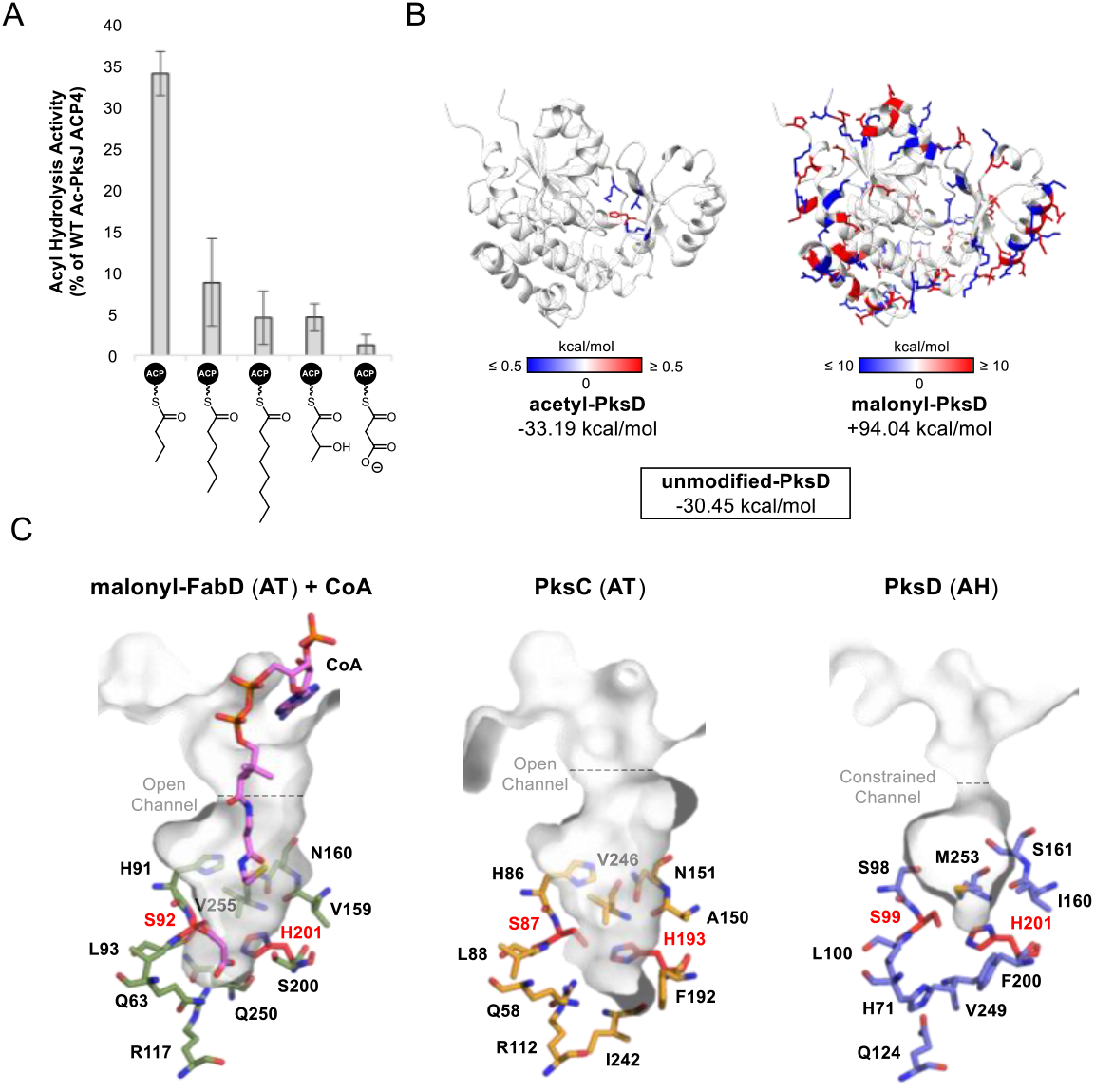
Substrate scope of PksD and analysis of the acyl chain binding pocket. **A**). Bar chart showing PksD-catalysed hydrolysis of acyl-PksJ ACP4 species (along x-axis) relative to acetyl-PksJ ACP4. PksD exhibits a clear preference for short, unbranched acyl chains. Only trace hydrolysis was detected for the malonyl-ACP species, in agreement with previous observations that AH domains are unable to process such substrates. Error bars represent the standard deviation of three replicates. **B**). Total interaction energy of Ser99-modified PksD with respect to the unmodified form. Differences in energy between acetyl-bound and unmodified (*left*) vs. malonyl-bound and unmodified (*right*) are shown. A negative interaction energy stabilises the system (blue), whereas a positive interaction energy destabilises the system (red). Major stabilising and destabilising residues are highlighted on the structure of PksD for the acetylated form (> +/-0.5 kcal/mol) and malonylated form (> +/-10 kcal/mol). **C**). Juxtaposition of the substrate binding pockets of FabD (PDB: 2G2Z), PksC (PDB: 5DZ6) and PksD (PDB: 8AVZ), suggesting a more constrained pocket and entrance channel in PksD. Pockets/channels are rendered as grey semi-transparent surfaces, with select residues shown as sticks coloured by atom.

Understanding how AH domains exert control over substrate specifcity could facilitate rational engineering efforts to broaden AH domain specificity, allowing diverse intermediates to be offloaded and thus *trans*-AT PKS pathways to be dissected. To study this aspect, we turned to our X-ray structure. As in AT domains from PKSs, Phe200 (strictly conserved) and the oxyanion hole-forming Gln12 (strictly conserved as Gln/His) of PksD are near the nucleophilic Ser99 residue (**Fig. 3A, 3B and 3D**). Juxtaposing a single chain of PksD (AH) with that of PksC (AT), PksE (**AT**-ER) (PDB: 5DZ7) and DisD (AT) reveals a less accessible substrate binding pocket in the former (**Fig. 4C, Fig. S10 – S12**). In PksD, the longer and generally less structured region between helices α3 and α4 (i.e. loop α3/α4), the Phe200-containing loop β6/α10 and the Gln12-containing loop β1/α1 associate through a series of van der Waals interactions involving Gln12, Gly13 and Gln15-Tyr17 (loop β1/α1), Lys58, Val60 and Pro63-Asp65 (loop α3/α4), Ile67 (helix α4), Pro195-Tyr198 and Phe200 (loop β6/α10), I128 (helix α6) and Val249 (helix α12), in addition to other residues. This intimate arrangement is stabilised by an extensive H-bonding network involving Phe64 (loop α3/α4), Gln15 (loop β1/α1), Gln12 (loop β1/α1), His71 (helix α4), Gln124 (helix α6) and Phe245 (helix α12); the oxyanion hole-forming Gln12 is also H-bonded Ser70 (helix α4) (**Fig. S10**). Ordered water molecules further confine these three loops, mediating H-bonds between Asp65 (loop α3/α4) and Ser197 (loop β6/α10), between Tyr17 (loop β1/α1) and Asp62 and Val60 (loop α3/α4) and between Gln15 (loop β1/α1) and Phe64. As well as restricting access to the active site of PksD, the aforesaid interactions constrain the positions of key side chains relative to the nucleophilic Ser99 to a greater extent than in AT domains (**Fig. S10 – S12**). For example, the compact side chain of Gln124 (by an Arg residue in ATs) participates in the aforementioned H-bonding network, drawing Val249 nearer to itself and to Phe200 and thereby providing hydrophobic/steric bulk that clashes with the charged malonyl group. The H-bonding network and van der Waals contacts also restrain the Gln12 side chain, which rests above the Phe200 phenyl ring and again clashes with the malonyl group. Further van der Waals interactions with Ile67, His71, Ile128, Met137, Val196, Ser197, Tyr198, His201 and Ile205 confine Phe200 in close proximity to Ser99. All of these factors likely limit the chain length and degree of branching of incoming substrates, with occasional thermal fluctuations allowing for exceptions.

That Gln124 is essential in shaping the substrate binding pocket of PksD is reflected in our finding of abolished activity for the Q124A variant. Although we sought to interrogate the role of Phe200, an F200A mutation failed to yield soluble protein, indicating that the phenyl ring – closely packed against the side chains of Met137, Ile160 and Val249 – is indispensable for proper folding of the AH domain. Taken together, PksD clearly adopts a very subtle mechanism for achieving substrate control, resulting in an active site architecture that is sensitive to point mutations. Producing a stable and active hydrolase with an expanded substrate profile, whilst desirable, may require a more focused engineering campaign (e.g. involving directed evolution) beyond the scope of this work.

### Analysing PksD-catalysed hydrolysis via molecular dynamics and quantum mechanics/molecular mechanics (QM/MM) simulations

To more closely examine the mechanism of PksD-catalysed acyl hydrolysis (i.e. steps IV-VI in **Fig. 3F**), we performed a series of classical MD (cMD; 50 ns) simulations followed by accelerated MD (aMD; 320 ns) on PksD with the Ser99 residue acetylated. In numerous aMD frames, water (WAT) molecules met the distance (His201[Nε2]→WAT[H] ≤ 3 Å, WAT[O]→Ac[C1] ≤ 3.2 Å) and angular (WAT[O]-Ac[C1]-Ac[O] = 100°-110°; i.e. approaching the Bürgi-Dunitz angle) requirements for a near-attack conformation.^34^ In 10 ns cMD simulations initiated from several such frames, the His201 side chain frequently recruited water molecules (**Fig. S13**) meeting the attack criteria. In a representative frame (**Fig. 5A** and **Video S1**), the carbonyl group of the substrate is activated toward nucleophilic attack by an oxyanion hole formed by Gln12 and Leu100. The methyl group of the substrate nestles against the side chains of His71, Phe200, Val249 and Gln12, which helps position the carbonyl group. Moreover, His71 is wedged between Phe200 and Val249 and stabilised by H-bonds to the side chains of Gln12 and Gln124 (as in **Fig. S10A** and **B**). In the representative snapshot (**Fig. 5A**), the methyl group of the substrate occupies a small binding pocket at the back of the active site, which cannot accommodate larger substituents. Larger substrates can only be accommodated by occupying the channel above this pocket. While the channel is mostly hydrophobic, a hydrophilic patch lining one side of it would disfavour binding of longer hydrocarbon chains. A cMD simulation with a hexanoyl group bound to the Ser99 residue identified a frame conforming to a near-attack conformation required for hydrolysis (**Fig. 5B** and **Video S2**). The reduced number of frames in the near-attack conformation for hexanoyl-PksD is reflected by hexanoyl-ACP thioesters hydrolysed with approximately 5% the efficiency of acetyl-ACP thioesters in our assays (**Fig. 4A**).

**Figure 5.**
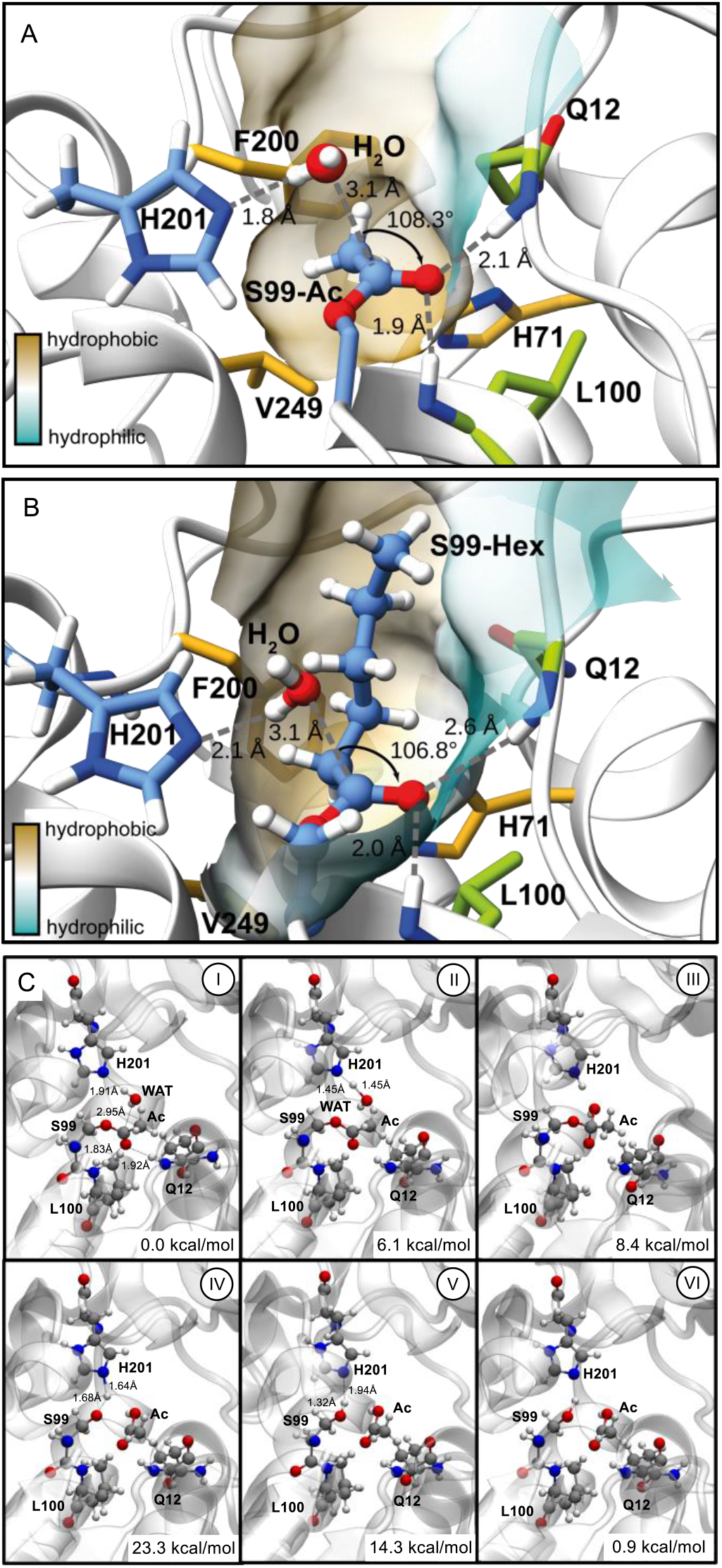
Snapshots of catalytic geometries for hydrolysis of acetyl- and hexanoyl-PksD. Example frame from cMD simulation of PksD with **A**). acetyl (Ac) and **B**). hexanoyl (Hex) groups bound to the catalytic Ser99 residue, in which distance and angular requirements for nucleophilic attack by a water (WAT) molecule are satisfied. In both cases, the backbone NH groups of Gln12 and Leu100 (green C atoms) form the oxyanion hole. Positioning of the acetyl group with respect to His201 and water is aided by nestling of the methyl group against the Phe200, Val249 and His71 side chains (gold C atoms). In contrast, the extended hexanoyl chain sites on the hydrophobic face of the end-capped pocket, explaining the severely limited hydrolysis of chains longer than C6. The semi-transparent surface rendering is coloured yellow (hydrophobic) or cyan (hydrophilic). **C**). Calculated structures along the catalytic reaction path for hydrolytic cleavage of an acetyl group by PksD. Bond lengths and energies associated with each step are shown.

In the case of malonyl-PksD, the geometry required for catalysis could only be met if the carboxylate group were to occupy a space similar to that of the methyl group in the acetyl substrate – an arrangement disfavoured by the hydrophobic nature of the pocket. However, in a 215 ns cMD simulation with malonyl-PksD, the carboxylate moves away from the hydrophobic part of the pocket, which leads to the carbonyl pointing towards H201 and no longer in a catalytically viable conformation (**Fig. S14** and **Video S3**). Taken together, these findings explain the inverse correlation we observed between the hydrolytic efficiency of PksD and the length of the alkyl chain, as well as the inactivity of PksD towards malonate substrates (**Fig. 4A**).

QM/MM calculations were performed to investigate the mechanism of the hydrolysis reaction. No catalytically competent structures were found for the malonyl-bound system (all reaction energies were highly endooergic). Representative structures from the MD trajectories for acetyl-PksD were optimised to obtain an initial reactant structure (**Fig. 5C**, panel I). An initial reaction path optimisation was attempted with the Quadratic String Method^35^; however, an optimised minimum energy path could not be obtained due to the large conformational changes that occur during the reaction. Based on this, we performed constrained optimisations of structures along the reaction path that are consistent with a possible reaction mechanism, namely: protonation of His201 by a water molecule (**Fig. 5C**, panel II), and the concomitant nucleophilic attack of the hydroxide anion to the carbonyl of the acetyl group, with an overall energy of 8.4 kcal mol^-1^ (**Fig. 5C**, panel III). The final process involves the neutralization of Ser99 by proton transfer from His201. The calculated structures associated with this process have energies of 23.3 and 14.3 kcal mol^-1^ (**Fig. 5C**, panel IV and panel V, respectively), leading to the final product, which results in a slightly endoergic overall reaction energy of 0.9 kcal mol^-1^ (**Video S4**).

The calculated energies associated with the structures along the reaction path are consistent with energies for enzymatic catalysis.^36,37^ The high energy associated with the final proton transfer step is due to suboptimal arrangement of the His201 and Ser99, which could be ameliorated by possible dynamics of the system, although no large conformational fluctuations for these residues were observed during the MD simulations. Moreover, alternative pathways could be possible such as product release prior to the proton transfer, which could allow for larger fluctuations of the His201 / Ser99 pair. However, these alternative processes could not be investigated due to computational limitations.

### Investigating the activity of PksD towards pantetheine and CoA thioesters

The efficient hydrolytic activity of AH domains towards short-chain acyl thioesters of ACP domains and N-acetylcysteamine^7,8^ raises questions about the fate of similar CoA thioesters in the native cellular milieu. Given their structural homology to AT domains, which accept CoA thioesters for acyl transfer, AH domains should process CoA thioesters unless a gatekeeping mechanism has evolved to prevent their hydrolysis. To test for such a mechanism, we began by probing the ability of PksD to process acetyl-CoA (**Fig. 6A**), using the same molar ratio as in assays with acetyl-PksJ ACP4. After 3 h, no PksD-mediated hydrolysis of acetyl-CoA was observed via LC-MS (**Fig. 6B**). We next performed this reaction in the presence of acetyl-pantetheine (acetyl-Pant; **Fig. 6A**), which lacks the 3ꞌ-phosphoadenosine moiety of CoA but closely emulates the acetylated Ppant moiety of ACP-bound substrates. In contrast to acetyl-CoA, acetyl-Pant was fully hydrolysed by PksD (compared to trace hydrolysis observed in an enzyme-free control; **Fig. 6C**).

**Figure 6.**
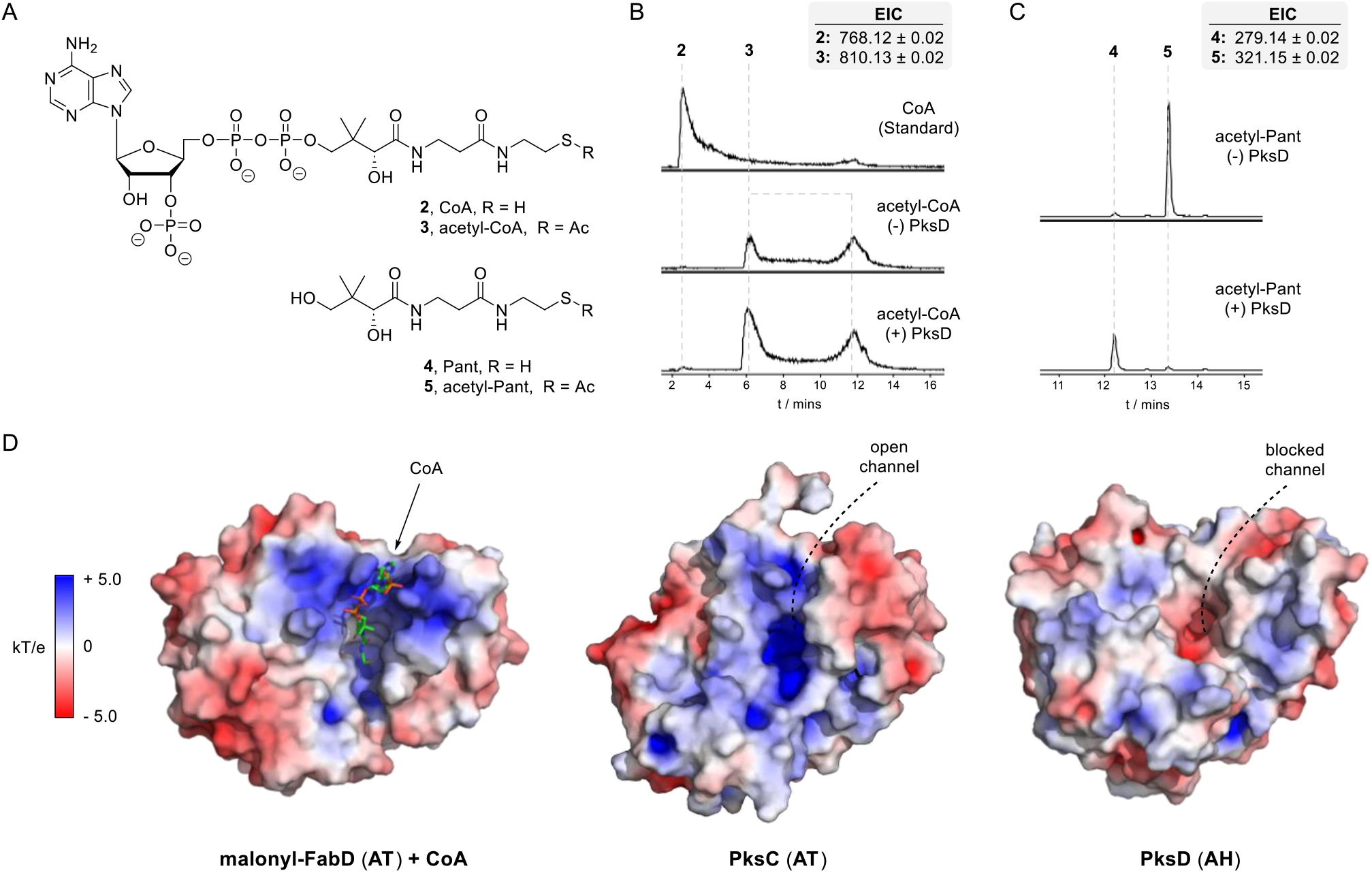
PksD selects against thioesters of coenzyme A. **A**). Chemical structures of CoA, acetyl-CoA, Pant and acetyl-Pant (**2** – **5**, respectively). **B – C**). Extracted ion chromatograms (EICs) from UHPLC-ESI-Q-TOF-MS analyses of PksD with acetyl-CoA or acetyl-Pant. **B**). No hydrolysis of acetyl-CoA was observed in the presence or absence of PksD (bottom and middle chromatograms, respectively), by comparison to a CoA standard (top chromatogram). Two peaks, each containing the correct mass ions, were observed for acetyl-CoA (likely due to different protonation configurations of the diphosphate) **C**). After 3 h, complete conversion of acetyl-Pant to Pant was observed in the presence of PksD (bottom chromatogram), whilst only trace Pant was observed in the negative control lacking enzyme (top chromatogram). **D**). Structures of FabD (PDB: 2G2Z), PksC (PDB: 5DZ6) and PksD (PDB: 8AVZ) displaying solution-phase surface electrostatic maps (calculated using APBS). The entrances to AT domains carry CoA-and malonyl-stabilising positive charges, whereas the same regions of AH domains carry no or negative charges.

To explain these findings, we inspected APBS-calculated electrostatic potentials along the surfaces of FabD (AT), PksC (AT) and PksD (AH).^38^ Whilst the channels leading to the active sites of the AT domains are laden with positive charges that stabilise the negatively charged CoA and malonyl moieties, the corresponding regions of PksD carry either negative or null charges that repel such groups (**Fig. 6D**). The exclusion of malonyl groups by AH domains is consistent with our and previously reported experiments.^7,8^ Surface renderings also highlight the lack of a defined 3′-phosphoadenosine-binding channel in PksD, when compared to FabD and PksC (**Fig. 6D**). Here, Tyr164 appears to occupy part of the channel in PksD, with the equivalent position an Ala or Ser in FabD and PksC, respectively (**Fig. S15**).

### Mapping the AH domain-binding epitope of the PksJ ACP4 domain using alanine scanning mutagenesis

Several structures of AT and ACP domains in complex and in isolation have provided insight into their domain-domain interaction interfaces, particularly the AT:ACP domain complex from the disorazol *trans*-AT PKS.^25^ Given the high degree of structural homology between AT and AH domains, we examined whether the two domain types exhibit similar interaction patterns towards their cognate ACP domains.

Inspired by previous successes,^12,39^ we employed an alanine scanning mutagenesis approach to define the AH:ACP domain interaction epitope on the PksJ ACP4 domain, using a previously reported library of X→Ala mutants.^40^ By testing these acetyl-loaded mutants for PksD-catalysed hydrolysis in our intact protein-MS assay, loss of productive interaction at the AH:ACP domain interface could be pinpointed. Six residues, when individually truncated to a methyl side chain, led to a significant decrease in PksD-catalysed hydrolysis: Asp32, Gln40, Asp41, Ile47, Asp65 and Tyr70 (**Fig. 7A** and **Fig. S16**). Notably, circular dichroism spectra of these variants closely resemble that of wild-type PksJ ACP4 domain, suggesting that no significant perturbation of secondary structure occurred upon mutation (**Fig. S17**). Mapping the six residues onto an AlphaFold model of the PksJ ACP4 domain allowed a binding epitope to be visualised,^41,42^ with most of the residues proximal to the point of Ppant attachment (i.e. Ser46) (**Fig. 7A**). The positioning of these key residues is in good agreement with the reported AT:ACP domain interfaces from *trans*-AT PKSs (e.g. DisD:DisA ACP1 and KirCII AT:KirAII ACP5),^39,43^ suggesting AH and AT domains bind the ‘top’ face of the ACP domain through a highly similar mode (**Fig. S18**).

**Figure 7.**
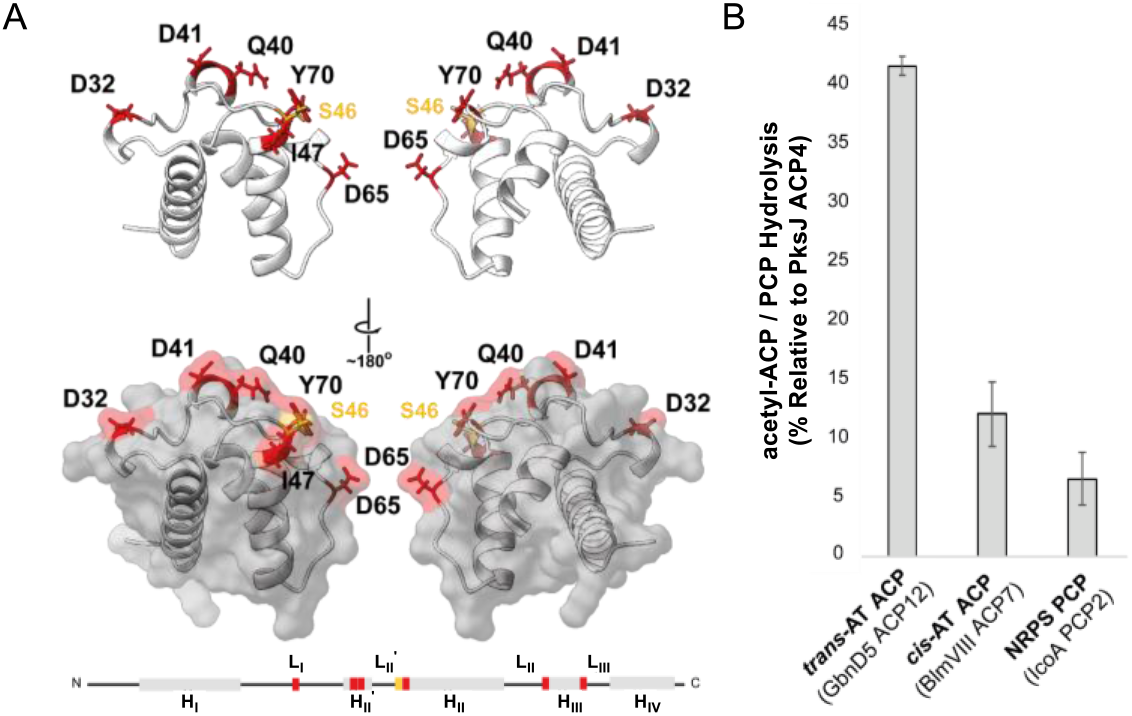
Identification of the PksD:PksJ ACP4 interaction epitope by scanning alanine mutagenesis. **A**). AlphaFold structure of PksJ ACP4 with significantly disruptive mutations (red carbons) and the nucleophilic Ser46 (orange carbons) shown as sticks. Semi-transparent surface renderings of the ACP domain reveal a solvent-accessible interaction epitope. A linear representation of the PksJ ACP4 structure is shown beneath and coloured according to the 3D representation. **B**). Bar chart showing PksD-catalysed hydrolysis of the acetyl group from select carrier protein domains relative to PksJ ACP4. PksD exhibits a clear preference for ACP domains from *trans*-AT PKSs, with <15% activity towards ACP/PCP domains from *cis*-AT PKS and NRPS pathways. Error bars represent the standard deviation of three replicates.

To expand this analysis, we examined the capacity of PksD to functionally interact with non-cognate carrier protein domains, including peptidyl carrier protein (PCP) domains from functionally related nonribosomal peptide synthetase (NRPS) assembly lines. We applied our intact protein-MS assay to a selection of acetyl-loaded carrier protein domains: GbnD5 ACP12 (from the gladiolin *trans*-AT PKS),^23,44^ BlmVIII ACP7 (from the bleomycin *cis*-AT PKS-NRPS)^45^ and IcoA PCP2 (from the icosalide NRPS)^46^ (**Fig. 7B**). Incubation with PksD resulted in 41 ± 8 %, 12 ± 1 %, and 7 ± 3 % acetyl group hydrolysis, respectively, relative to the acetyl-PksJ ACP4 domain. Whilst our findings demonstrate a clear preference for acetyl-loaded carrier protein domains from *trans*-AT PKSs, those from *cis*-AT PKSs and NRPSs could also be processed, albeit at much slower rates. Variation of key residues involved in binding PksD appears to be responsible for the decreased activity against non-cognate carrier protein domains, with several instances of charge reversal likely resulting in a suboptimal interaction interface (**Fig. S23**).

### Elucidating the ACP domain-binding interface of PksD by carbene footprinting

Subsequently, we sought to visualise the interaction epitope from the perspective of the AH domain. Deletions of helix α17 (Δ302-324) and helices α16-α17 (Δ290-324) in the C-terminal latch region yielded insoluble protein, precluding downstream analysis but indicating this region is important for correct folding. We next turned to carbene footprinting, a structural MS technique wherein masking or unmasking of residues due to decreased or increased solvent accessibility, respectively, is detected by LC-MS analysis after covalent modification with a carbene probe and proteolytic cleavage into peptides.^47^ This technique has been employed in mapping the interactions between cognate ACP and catalytic domains in related systems.^40,44,48^

When we applied this approach to a PksD:*holo*-PksJ ACP4 domain complex, we observed a localised masked region consisting of residues 270-294 (strand β9 -helix α16) of the AH domain. These residues are situated at the entrance to the substrate binding pocket and comprise the equivalent surface occupied by ACP domains in reported AT:ACP domain complexes (**Fig. 8** and **Fig. S20**).^25,49^ Most other detected peptides showed no change in the presence of the ACP domain; however, part of helix α2 (residues 26-31) and helix α9 (residues 180-182) were masked and unmasked, respectively, suggesting subtle conformational shifts in both subdomains. This is unsurprising given the previously reported conformational mobility of AT domains,^50,51^ and highlights the dynamic nature of these interactions.

**Figure 8.**
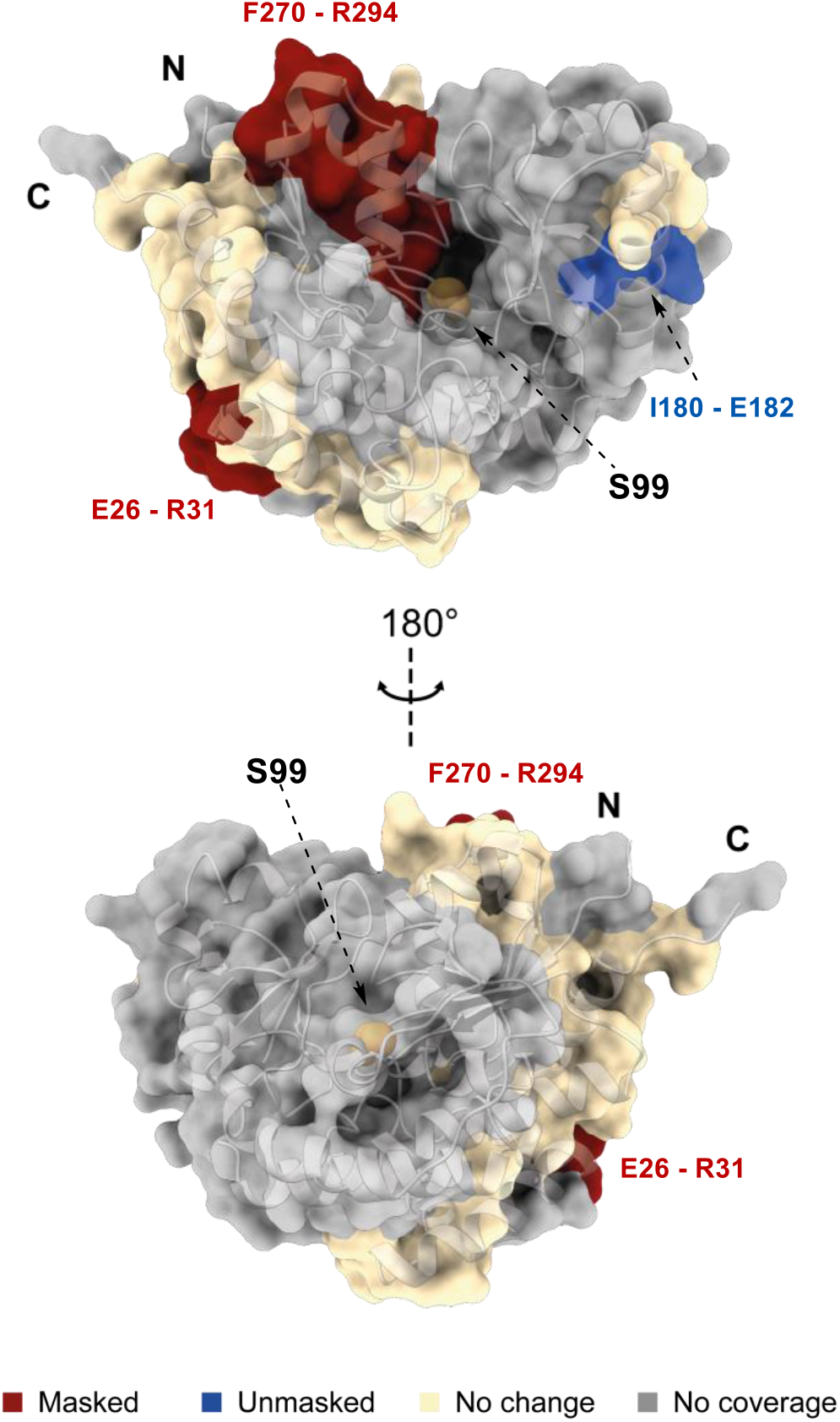
Carbene footprinting of the PksD:*holo*-PksJ ACP4 complex. Structure of PksD showing the locations of masked, unmasked and unaffected peptide regions resulting from trypsin and GluC digests in the presence of the *holo*-PksJ ACP4 domain. The N-and C-termini are labelled and the active site Ser99 residue is shown as orange spheres. An extended masked region is situated at the entrance to the active site cavity, which is the likely docking site of the ACP domain. The localised unmasking is attributed to conformational changes in PksD upon ACP binding, as observed during MD simulations (**Fig. S25**).

### Computational docking and analysis of the PksD:PksJ ACP4 complex

As discussed above, our alanine scanning mutagenesis and carbene footprinting experiments yielded near-residue information on the AH:ACP domain interaction interface. Distance restraints from these datasets were applied to docking and MD simulations starting from our crystal structure of PksD and an AlphaFold model of the PksJ ACP4 domain.^41,42^ The *apo*-PksJ ACP4 was manually docked onto PksD using virtual reality in ChimeraX;^52^ acetyl-Ppant was then attached to Ser46 residue of the PksJ ACP4 domain and fed into the active site of PksD. Notably, AlphaFold Multimer,^53^ which became available during the preparation of this manuscript, predicts a similar position for the ACP domain but rotated ∼180° around the axis of the substrate channel of PksD (**Fig. S21**).

The resulting complex was then energy-minimised and subjected to 50 ns of cMD followed by 576 ns of aMD to explore conformational changes. We found that the ACP domain rotates with respect to PksD and exhibits a degree of conformational plasticity (especially with respect to helices II and III; **Fig. S21**). We also monitored geometries relevant to acyl transfer (i.e. steps I - II in **Fig. 3F**), and from three aMD frames approaching attack we initiated further 20 ns cMD simulations. In all three simulations, multiple frames matched the criteria for acyl transfer (**Fig. 9**, **Fig. S22** and **Video S5**).

**Figure 9.**
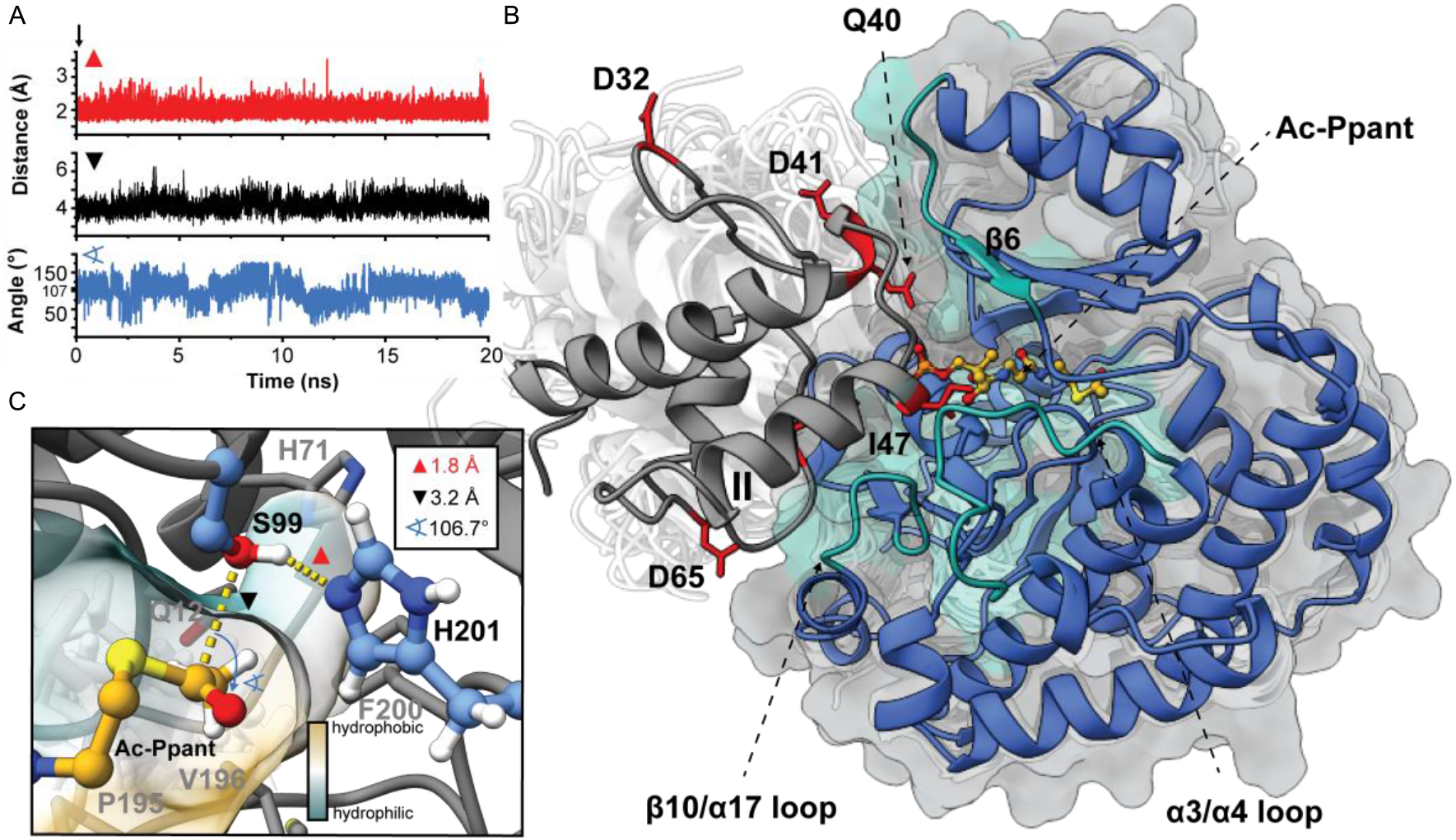
Representative cMD frame of the docked complex of PksD with the acetyl-PksJ ACP4 domain. **A**). Distance (Ser99[Oγ]→His201[Nε], red trace; Ser99[Oγ]→Ac[C1]-Ppant, black trace) and angle (Ser99[Oγ]-Ac[C1]-Ppant-Ac[O]-Ppant, blue trace) plots during a 20 ns cMD simulation of the PksD:acetyl-Ppant-PksJ ACP4 domain complex. **B**). Overview of the PksD:acetyl-Ppant-PksJ ACP4 domain complex. The position of the illustrated frame along the trajectory is indicated with an arrow in panel A. PksD is rendered as a blue cartoon and light grey semi-transparent surface and the PksJ ACP4 domain as a dark grey cartoon. PksD regions that interact with the PksJ ACP4 domain during the course of the simulation are highlighted in green and labelled. PksJ ACP4 domain surface residues identified by alanine scanning mutagenesis as important for productive binding to PksD are shown as red sticks; acetyl-Ppant C atoms are shown as yellow ball-and-sticks. White semi-transparent cartoons depict the range of conformations sampled by PksD and PksJ ACP4 in a subsequent 576 ns aMD simulation. **C**). Magnified view of the substrate binding pocket. Semi-transparent surfaces along the pocket are coloured according to hydrophobicity (yellow for hydrophobic, cyan for hydrophilic). Semi-transparent sticks illustrate the range of conformations explored by the acetyl-Ppant moiety during the 576 ns aMD simulation.

In a representative frame (**Fig. 9**), the ACP domain binds to PksD in an orientation nearly identical to that observed between the DisA ACP1 domain and the DisD AT (PDB: 5ZK4) (**Fig. S21B**, **S21D** and **S23**).^25^ However, the PksJ ACP4 domain is translated ‘forward’ relative to the DisA ACP1 domain such that helix II nestles between loop β10/α17 (on the C-terminal latch of PksD) and the lengthy loop α3/α4 (on the front face of PksD) – two features absent from DisD (**Fig. S23** and **S24**). This forward displacement also permits the compact helix I to associate with strand β6 and neighbouring loops of PksD. Finally, whereas the DisA ACP1 domain employs helix III to stack against the C-terminal helix α13 of DisD, such a feature is absent between helices II and III of the PksJ ACP4 domain and would otherwise clash with the C-terminal latch. Instead, an elongated loop interacts extensively with helix α15 and the C-terminal latch of PksD (**Fig. S19**). In a crosslinked structure of that AT VinK with the PCP VinL (PDB: 5CZD) from the vicenistatin *trans*-AT PKS, VinL exhibits a substantially different binding mode, despite the presence of a C-terminal latch on VinK (**Fig. S23**).^25,49^ As hinted at previously, other structural mismatches on VinK may be responsible for this discrepancy, including a truncated helix preceding the latch as well as features of the latch itself.^25^

Epitope-forming residues on the PksJ ACP4 domain identified by alanine scanning mutagenesis either sit directly at the AH domain interface (e.g. Gln40, Asp41, I47) or engage in stabilising interactions during the course of our MD simulations (e.g. Asp32, Asp65 and Tyr70). Moreover, helix α9 becomes more solvent-exposed due to lateral movement of 3_10_ helix α8 and the adjacent loop containing Asn159 and Ser160 (**Fig. S25**). In agreement with our carbene footprinting data, helix α9 (residues 180-182) shows unmasking upon ACP domain binding (**Fig. 8** and **Fig. S20**).

A representative frame (**Fig. 9C**) illustrates the spatial limitations of the substrate binding pocket in fitting long (> 6 carbons) or branched acyl chains, although the pocket does possess some dynamic qualities (**Fig. 9C** and **Fig. S26**). The channel features both hydrophobic and hydrophilic faces, which likely facilitates proper orientation of the acetyl-Ppant moiety for acyl transfer (**Fig. S26C** and **S26D**). We also carried out MD simulations on a docked complex between PksD and the malonyl-PksJ ACP4 domain to investigate how the AH domain behaves toward this non-preferred substrate. During the 280 ns aMD simulation, no single frame satisfies the required geometry (**Fig. S27**). The malonyl group appears unable to acylate Ser99 due to transient H-bonds between the carboxylate group and a range of polar residues near the active site, in addition to steric factors in the active site.

## Conclusion

In conclusion, we performed the first structure-function analysis of an AH domain, which rescues stalled *trans*-AT PKSs by hydrolysing short acyl chains aberrantly carried by ACP domains. Our crystal structure and biochemical assays allowed us to propose mechanisms for the selection and hydrolysis of short acyl (e.g. acetyl) chains carried by ACP domains but not by CoA. Moreover, our interrogation of the AH:ACP domain interface via scanning alanine mutagenesis coupled with LC-MS assays, carbene footprinting and MD simulations revealed a binding mode similar to that observed for an AT:ACP domain pair with small but significant deviations. Finally, we demonstrated AH domain activity against ACP and PCP domains from *cis*-AT PKSs and NRPSs, respectively. Our findings expand the knowledge of *trans*-AT PKS enzymology and may find use in bioengineering experiments or those designed to offload biosynthetic intermediates for subsequent analysis.

### Data Availability

The structure factor amplitudes and atomic coordinates of PksD have been deposited in the RCSB Protein Data Bank under PDB codes 8AVZ (SeMet derivative) and 8AW0 (native protein). The topology and trajectory files for the MD simulations discussed in the manuscript can be found on Zenodo DOI 10.5281/zenodo.7657414.

### Author Contributions

M.J. conceived and designed the study. C.D.F. crystallised PksD in native and SeMet forms, solved and refined their structures. M.J. and M.P. generated expression constructs, overproduced and purified proteins, constructed mutant plasmids and performed biochemical assays. X.J. produced the pHis_8_ expression vector used in this study and the IcoA PCP2 construct. H.G.S. synthesised the pantetheine and acetyl-pantetheine substrates. G.L.C. supervised the synthetic chemistry and assisted with interpretation of enzymatic activity and computational simulation data. M.J. performed and analysed carbene footprinting experiments. B.P.T. and J.R.L. conducted docking studies and molecular dynamics simulations on the substrate scope and protein interface. U.C.D. and G.A.C. conducted energy decomposition analyses and QM/MM calculations on the acetyl- and malonyl-bound forms of PksD. C.D.F. J.R.L. and M.J. wrote the manuscript with input from all authors.

## Acknowledgements

This work was supported by a Biotechnology and Biological Sciences Research Council (BBSRC) Discovery Fellowship to M.J. (BB/R012121/1). C.D.F, J.R.L and G.L.C acknowledge funding from the BBSRC (BB/R010218/1). M.P. is supported by a Midlands Integrative Bioscience Doctoral Training Partnership studentship (BB/M01116X/1). B.P.T. gratefully acknowledges Bruker and the University of Warwick for funding as part of the Warwick Centre for Doctoral Training in Analytical Science. H.G.S. was supported by an MRC IBR Doctoral Training Partnership (MR/N014294/1) Studentship and IAS Early Career Fellowship from the University of Warwick. X.J. was supported by fellowships from the University of Warwick and the China Scholarship Council. Some of the computing facilities were provided by the Scientific Computing Research Technology Platform of the University of Warwick. This study was partially funded by NIH Grant No. R01GM108583 to G.A.C. Computational time from CASCaM (NSF Grant Nos. CHE-1531468 and OAC-2117247), NSF Extreme Science and Engineering Discovery Environment, XSEDE (Project No. TG-CHE160044), and the University of Texas at Dallas’ CIRC are gratefully acknowledged. The Bruker MaXis II instrument used in this study was funded by the BBSRC (BB/M017982/1). The authors are grateful to Dr Nikola Chmel for assistance with circular dichroism measurements.

## TABLE OF CONTENTS

Acyl hydrolase (AH) domains catalyse the removal of short chain thioesters from carrier proteins in *trans*-acyltransferase polyketide synthases. The first structure of an AH domain is reported providing insights into catalytic mechanism, substrate tolerance and interface with its carrier protein-bound substrate.

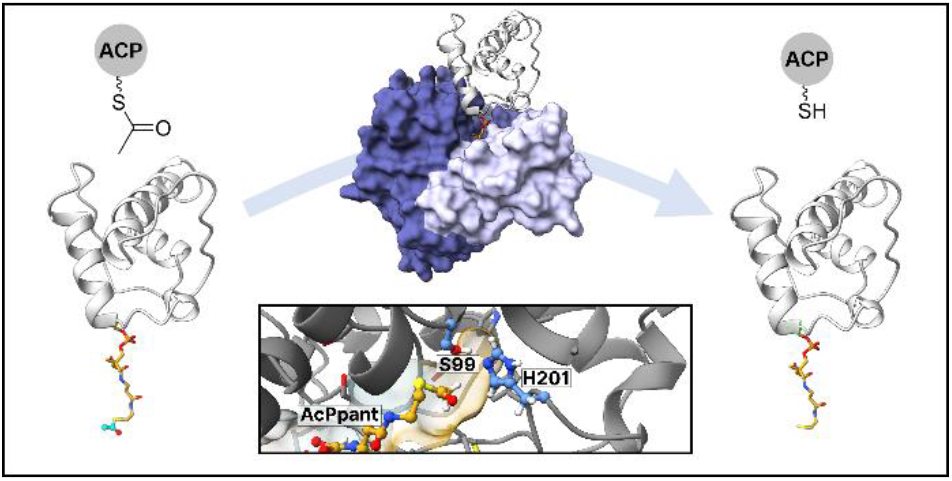

