## Supplementary Information for "Molecular basis for short-chain thioester hydrolysis by acyl hydrolase domains in *trans*-acyltransferase polyketide synthases"

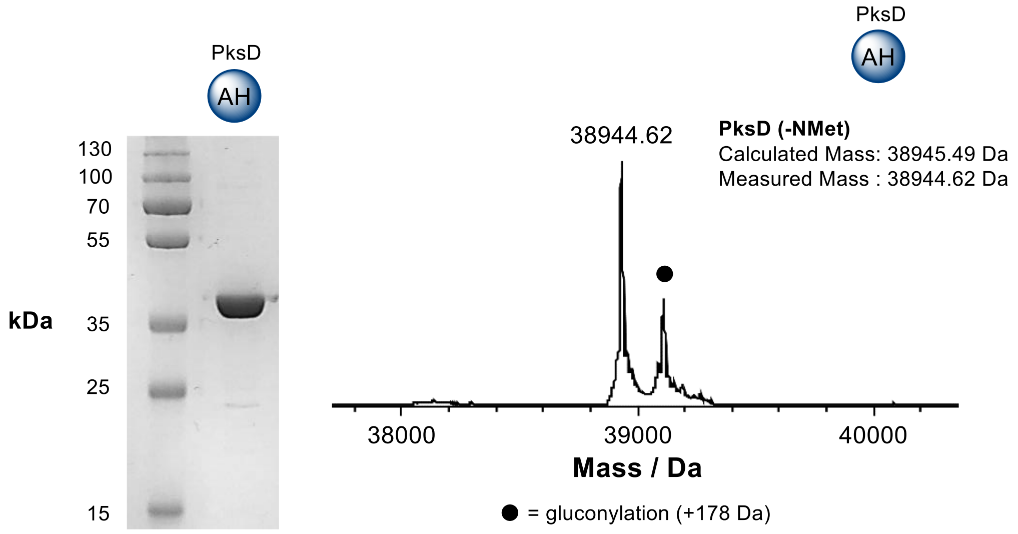

**Figure S1. SDS-PAGE and MS analysis of PksD.** Left: 10% SDS-PAGE gel of the ~39 kDa PksD domain following Ni-NTA purification. Right: Deconvoluted mass spectrum of purified PksD, with calculated and observed masses detailed. Note that gluconylation occurs as a posttranslational modification of the N-terminal His-tag during protein expression.

A

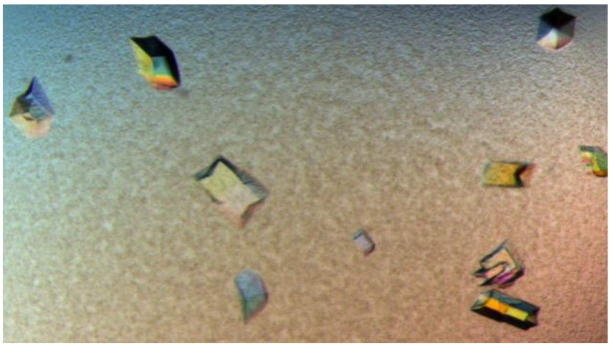

B

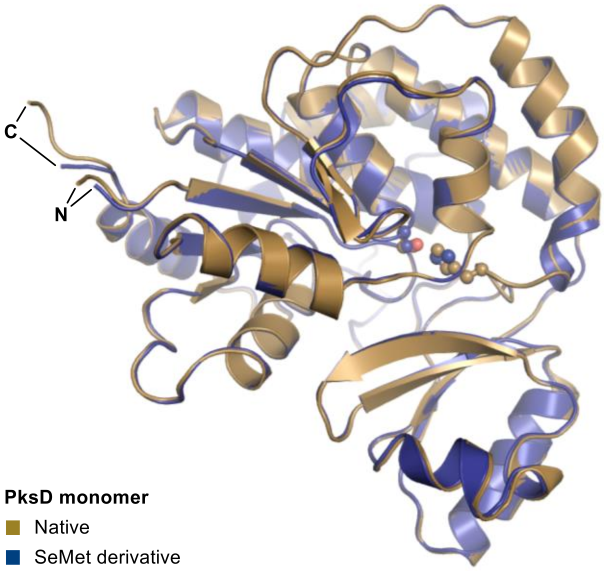

**Figure S2. Crystals of native PksD and structural comparison to SeMet-derived PksD.** A). Representative native PksD crystals, photographed under microscopic magnification. B). Superposition of chain A of SeMet-derived (blue) and native (sand) PksD shows minimal differences in backbone conformation with a C$\alpha$ r.m.s.d. of 0.186 Å for 285 atoms. Side chains of the catalytic dyad residues Ser99 and His201 are depicted as ball-and-sticks coloured by atom.

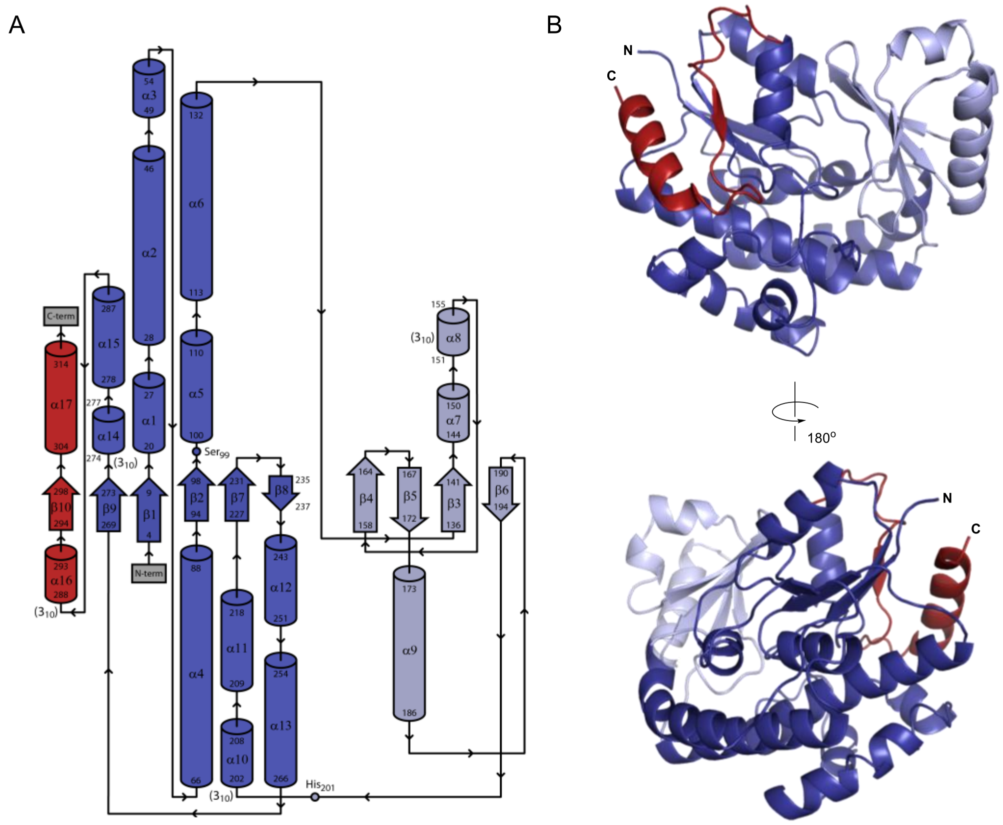

**Figure S3. Analysis of PksD structural elements.** A). Topology map of PksD. Helices and strands are shown as cylinders (α1-17) and block arrows (β1-10), respectively, and flexible loops as thin black arrows. All arrows point from N- to C-terminus. Positions of the catalytic dyad residues Ser99 and His201 are indicated. Colours: large subdomain (dark blue), small subdomain (light blue), C-terminal latch (red). Image derived from PDBsum server[1] output. B). Structure of PksD (chain A, SeMet derivative) from front and back views. Colouring of structural elements matches the topology map in panel A. Note that some secondary structures, such as helix α16, were not rendered by PyMOL due to inequivalent boundary definitions.

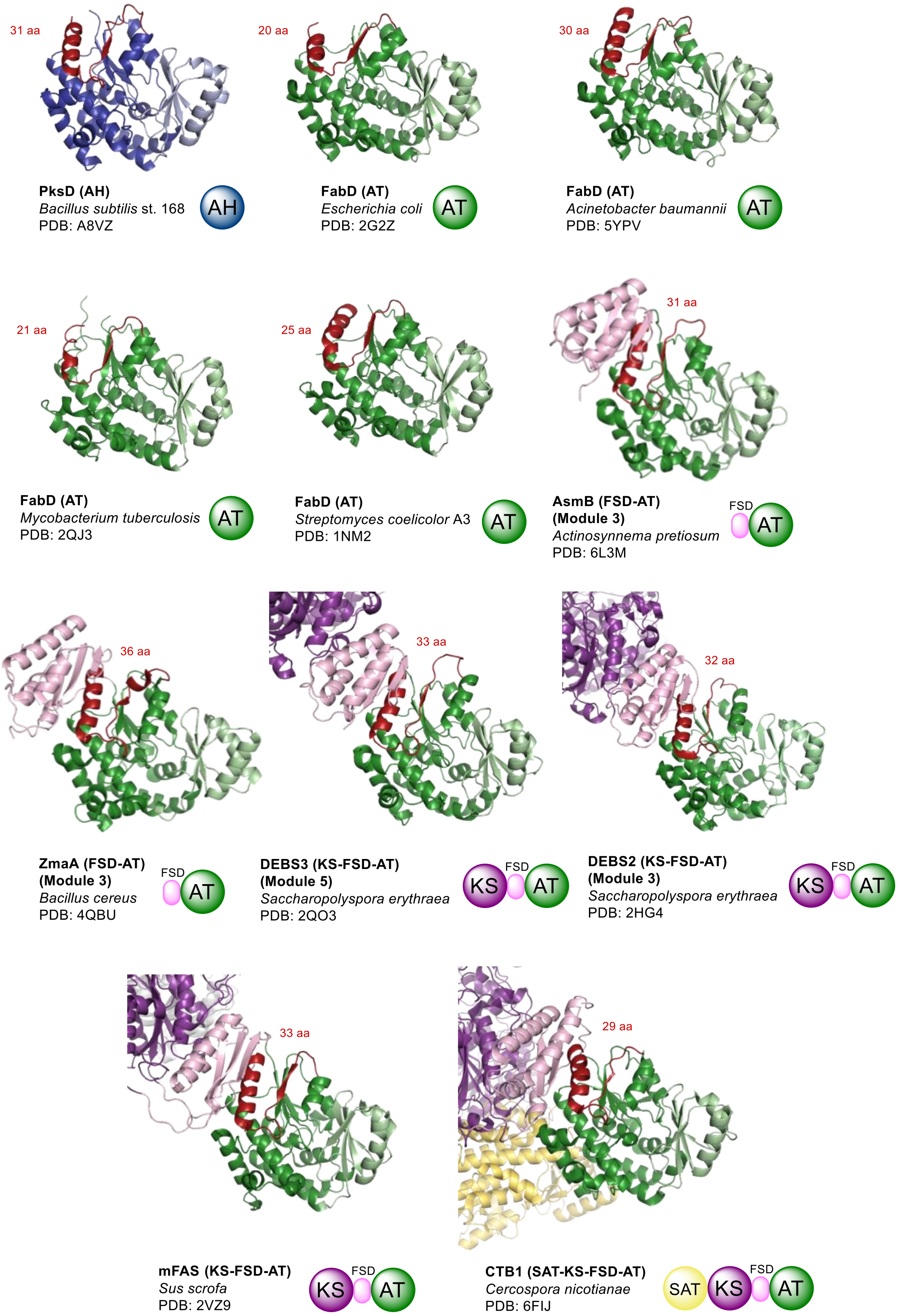

**Figure S4. Occurrence of the C-terminal latch in other structures of AT domains.** Examples of AT domain structures that possess a C-terminal latch region (highlighted in red, with number of constituent residues (aa) denoted above). The domain organisation for each structure is depicted in coloured circles, with corresponding PDB identifiers and producing organisms shown. Abbreviations: SAT (starter unit acyltransferase), FSD (flanking subdomain).

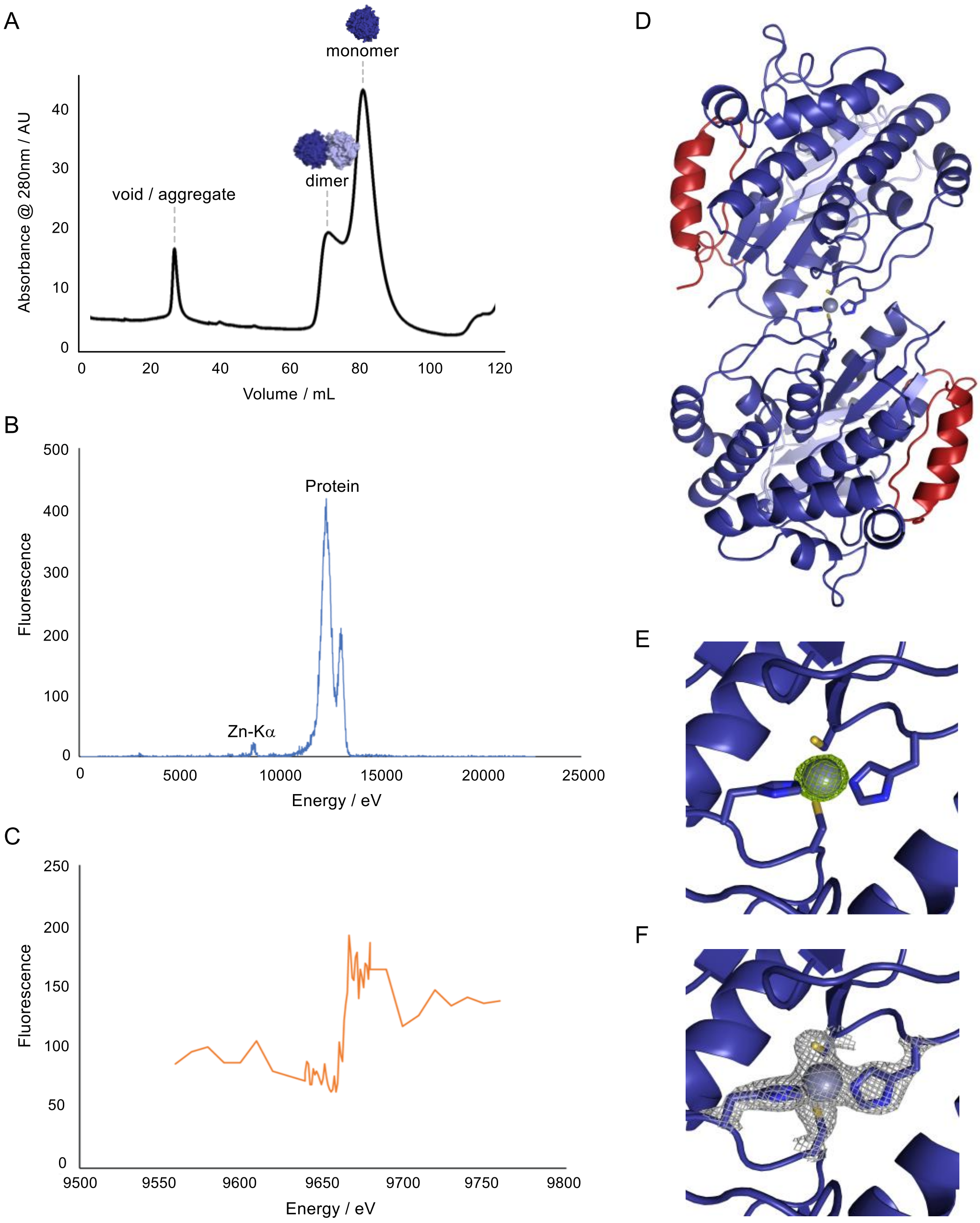

**Figure S5. Detection of zinc at the dimer interface of PksD crystals.** A). Elution profile of PksD from a HiLoad 16/600 Superdex S200 pg size-exclusion chromatography column, showing two overlapping peaks for monomeric and dimeric forms. B). X-ray fluorescence emission spectrum, revealing a small but significant peak for zinc (Zn-Kα). C). X-ray florescence zinc edge scan. Calculated peak energy = 9666.0 eV, $f''$ = 6.27 e, and $f'$ = -9.39 e. Calculated inflection energy = 9663.0 eV, $f''$ = 2.94 e, and $f'$ = -10.49 e. D). Structural representation of the PksD dimer (SeMet derivative), showing coordination of a $Zn^{2+}$ ion (purple sphere) via Cys236 and His238 residues (sticks) supplied by each of two monomers. PksD is coloured as in **Fig. 2**. E-F). Electron density maps of the $Zn^{2+}$ binding site between chains A and C of SeMet-derived PksD: E). Polder omit map (σ = 10.0) and F). $2F_o$-$F_c$ map (σ = 2.5).

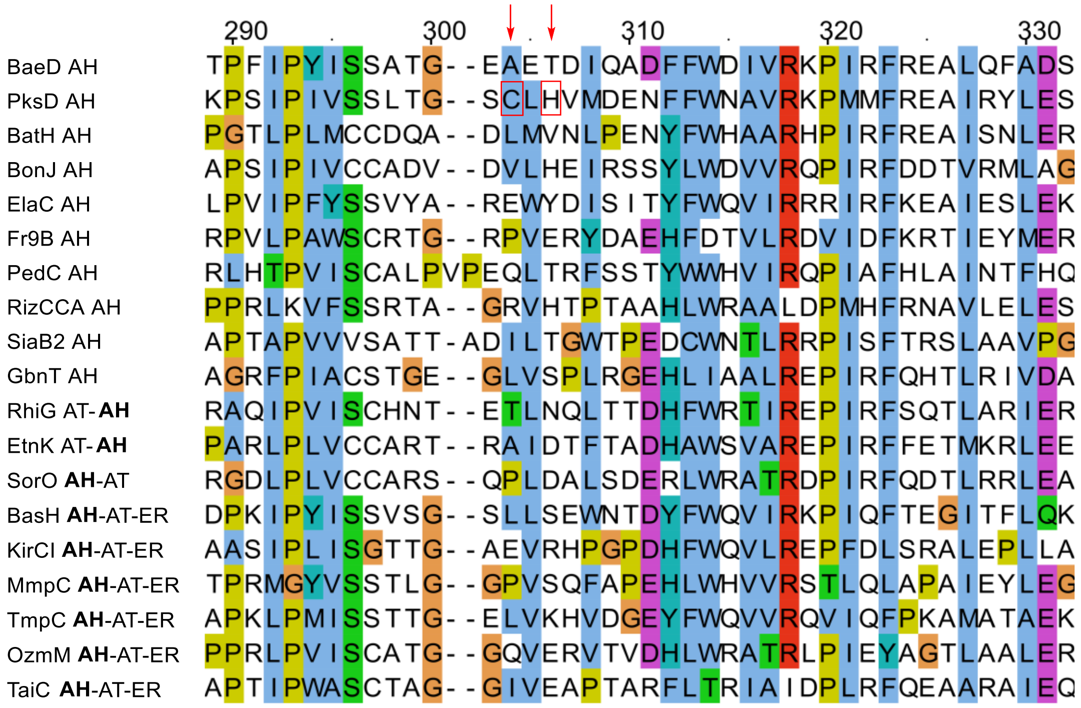

**Figure S6. Multiple sequence alignment of AH domains showing the lack of conservation of zinc-binding residues.** Positions of the $Zn^{2+}$-coordinating Cys236 and His238 of PksD are indicated with red arrows and boxes. In the case of multi-domain proteins, the relevant AH domain is bolded at left. All sequences were collected from the MIBiG database,[2] aligned using Clustal Omega[3] on the EMBL-EBI server[4] and visualized in JalView.[5]

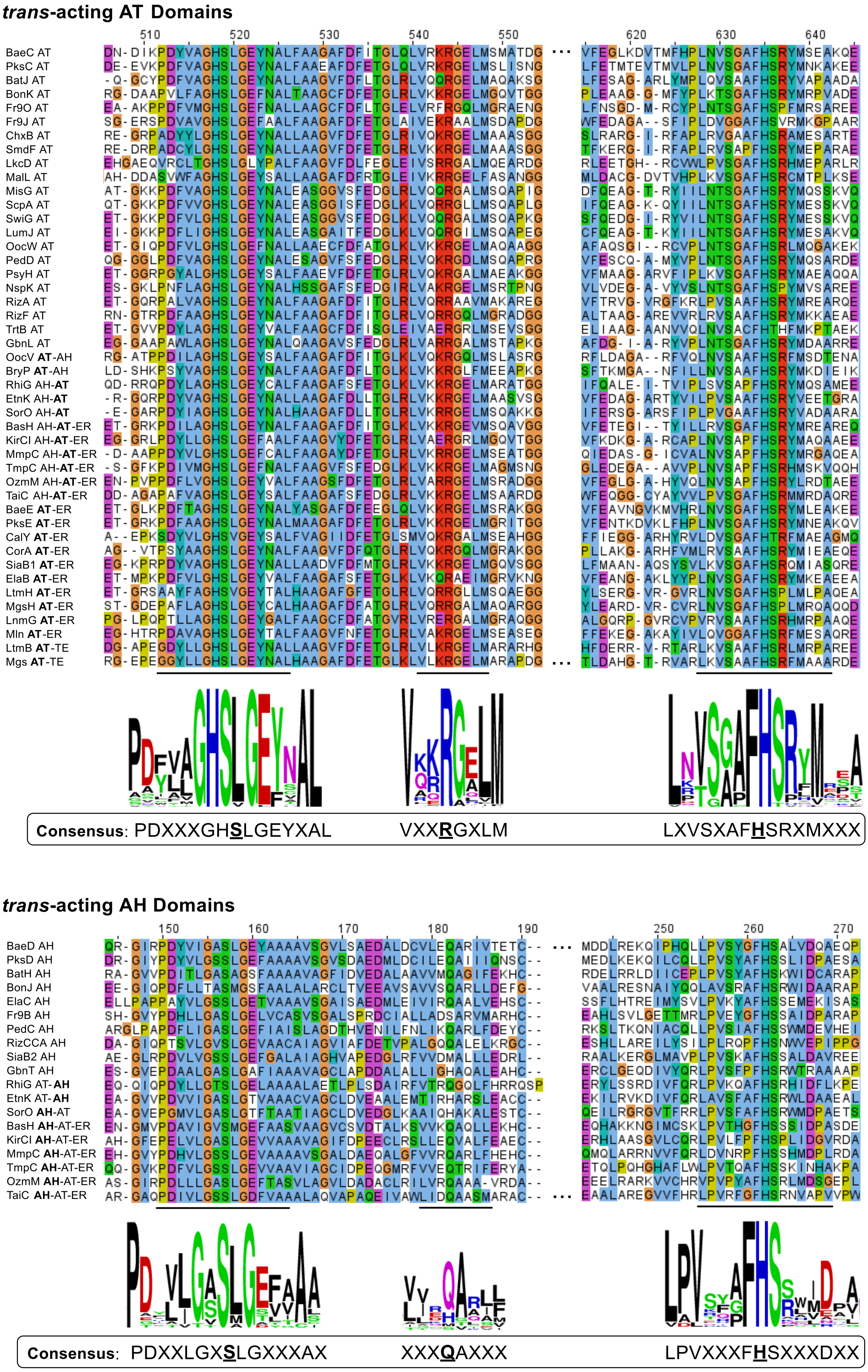

**Figure S7. Multiple sequence alignment of AT and AH domains from *trans*-AT PKSs identifies signature active site motifs.** Regions containing the active site Ser, His and Arg/Gln are shown, and consensus sequences around these residues are highlighted. In the case of multi-domain proteins, the relevant domain is bolded at left. All sequences were collected from the MIBiG database,[2] aligned using Clustal Omega[3] on the EMBL-EBI server[4] and visualized in JalView.[5] Sequence logo diagrams were constructed using the WebLogo server.[6]

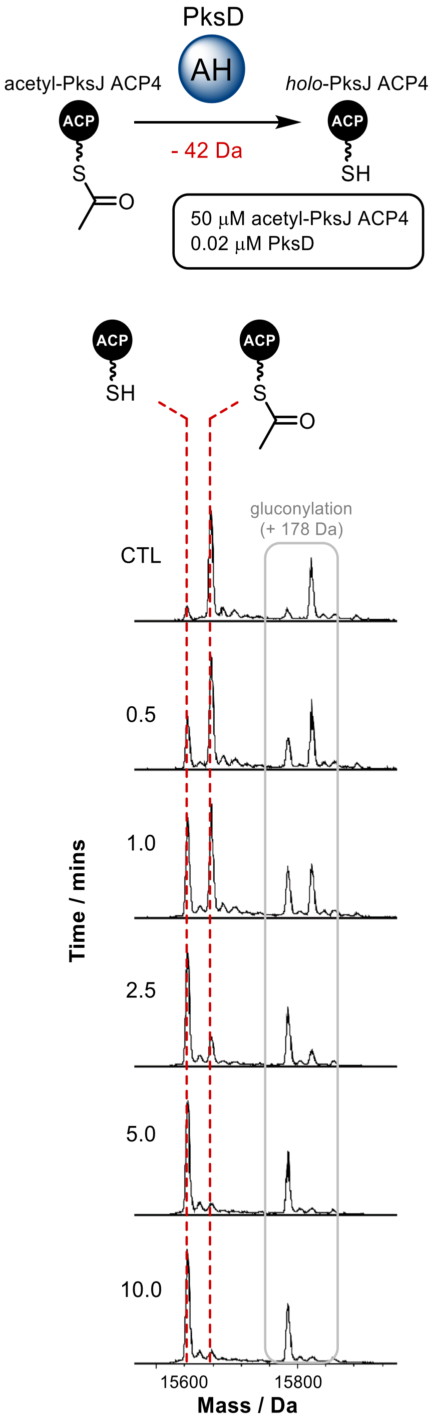

**Figure S8. Time course of PksD catalytic activity, monitored by intact protein-MS.** Top: reaction scheme and conditions for PksD-catalysed hydrolysis of acetyl-PksJ ACP4. Bottom: stacked, deconvoluted ESI-Q-TOF spectra of representative timepoints from the hydrolysis reaction. Hydrolysis results in a -42 Da mass shift. Peaks corresponding to the *holo*-ACP (product) and acetyl-ACP (substrate) species are highlighted with dotted red lines. A negative control reaction (CTL) was performed in the absence of PksD. Peaks corresponding to N-terminal gluconylation are indicated (see also **Fig. S1**).[7]

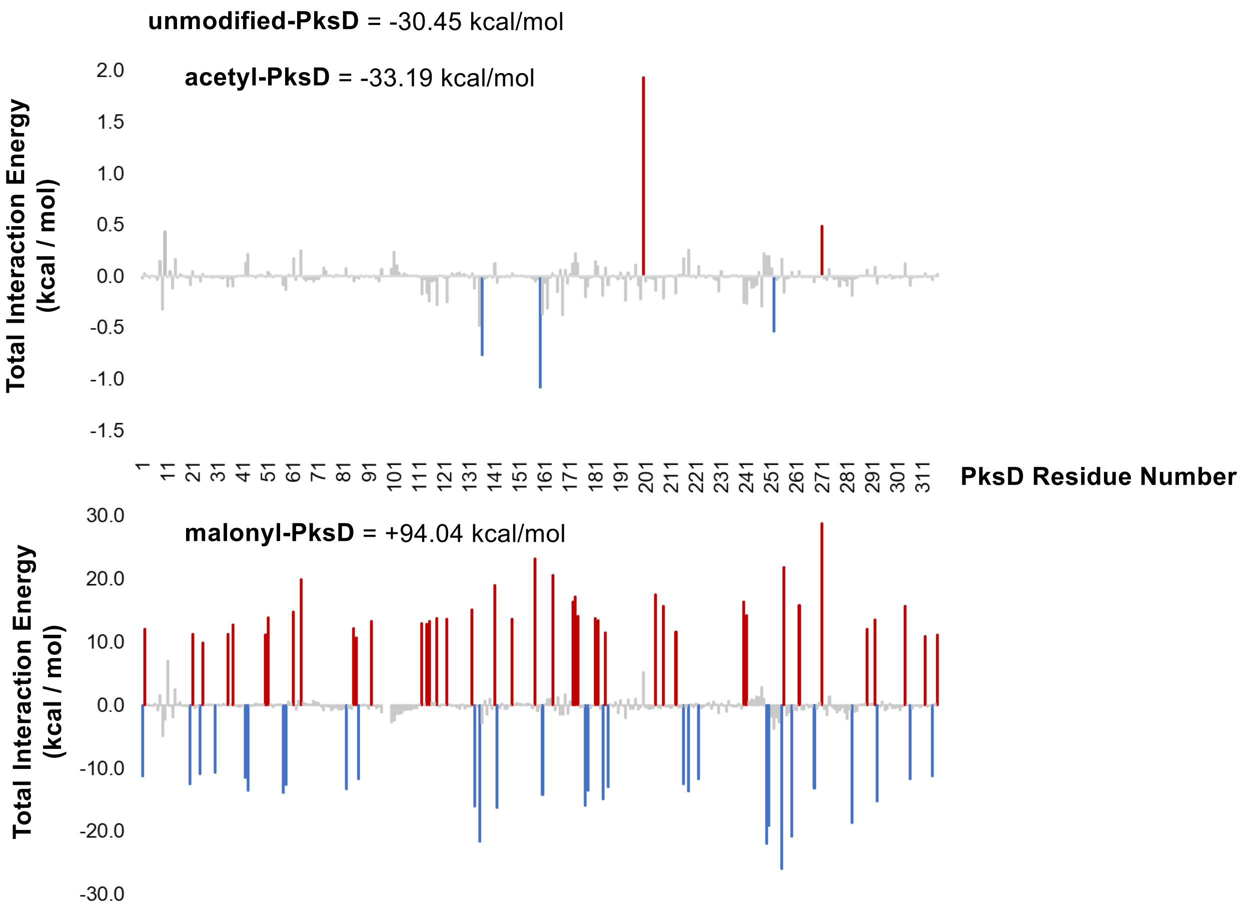

**Figure S9. Energy decomposition analysis of acetyl- and malonyl-PksD.** Total interaction energy of Ser99-modified PksD with respect to the unmodified form. Differences in energy between acetyl-bound and unmodified (*top*) vs. malonyl-bound and unmodified (*bottom*) are shown. A negative interaction energy stabilises the system (blue), whereas a positive interaction energy destabilises the system (red). Major stabilising and destabilising residues are highlighted in the plots.

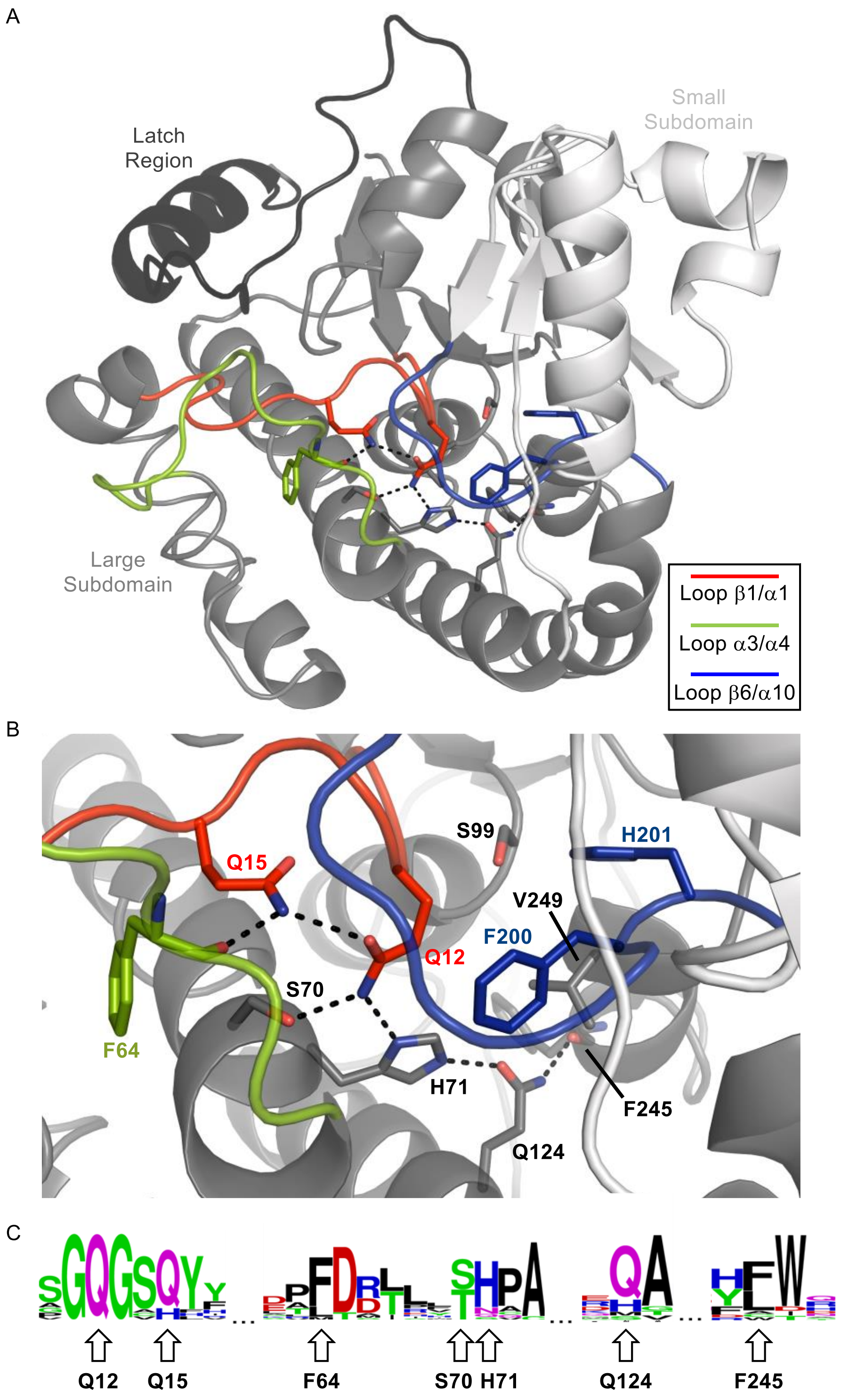

**Figure S10. A network of hydrogen bonds promotes constriction of the substrate binding pocket of PksD.** A). Overall view of PksD and B). magnified view of the pocket. Acyl-Pant substrates linked to ACP domains enter the pocket from above/between Ser99 and His201 as depicted. The catalytic dyad and select residues that shape the substrate binding pocket are shown as sticks. Loops that enclose the pocket, named according to bounding secondary structures from PksD, are highlighted as in panel A (note that helix α3 was not rendered as a helix by PyMOL). C). Consensus sequences of amino acids involved in constraining the active site, generated by the WebLogo server.[6] Residue numbering corresponds to PksD.

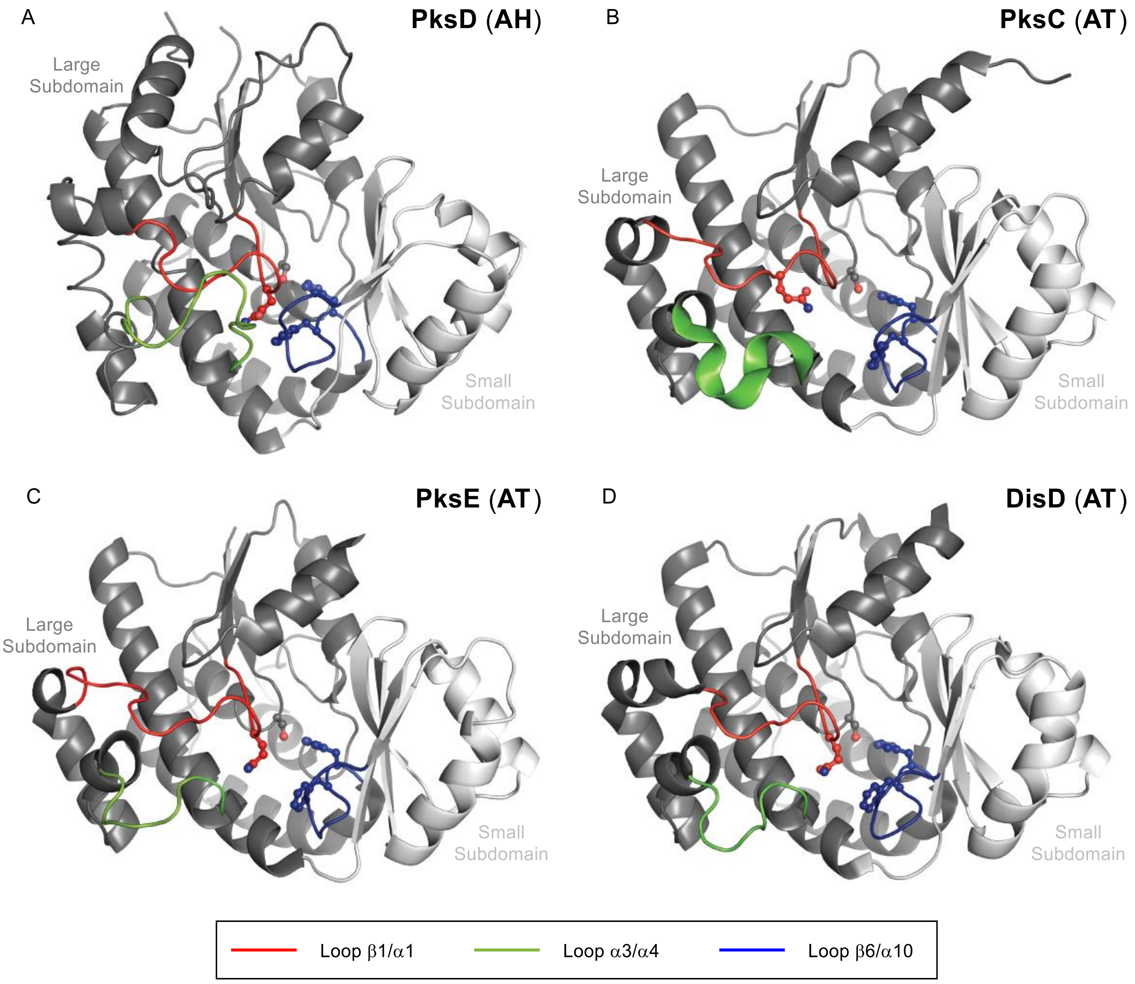

**Figure S11. Structural comparison of binding pocket loops among AH and AT domains (cartoon rendering).** Juxtaposition of A). PksD (PDB: 8AVZ) with B). PksC (PDB: 5DZ6), C). PksE (PDB: 5DZ7; ER domain not shown) and D). DisD (PDB: 3RGI) reveals a more constricted substrate binding pocket for PksD due to the tighter interactions of loops enveloping the pocket (e.g. see hydrogen bonding network in **Fig. S10**). These loops are named according to bounding secondary structures in PksD. Gln12, Ser99, Phe200 and His201 of PksD and corresponding AT domain residues are depicted as ball-and-sticks coloured by atom. See also **Fig. S12**.

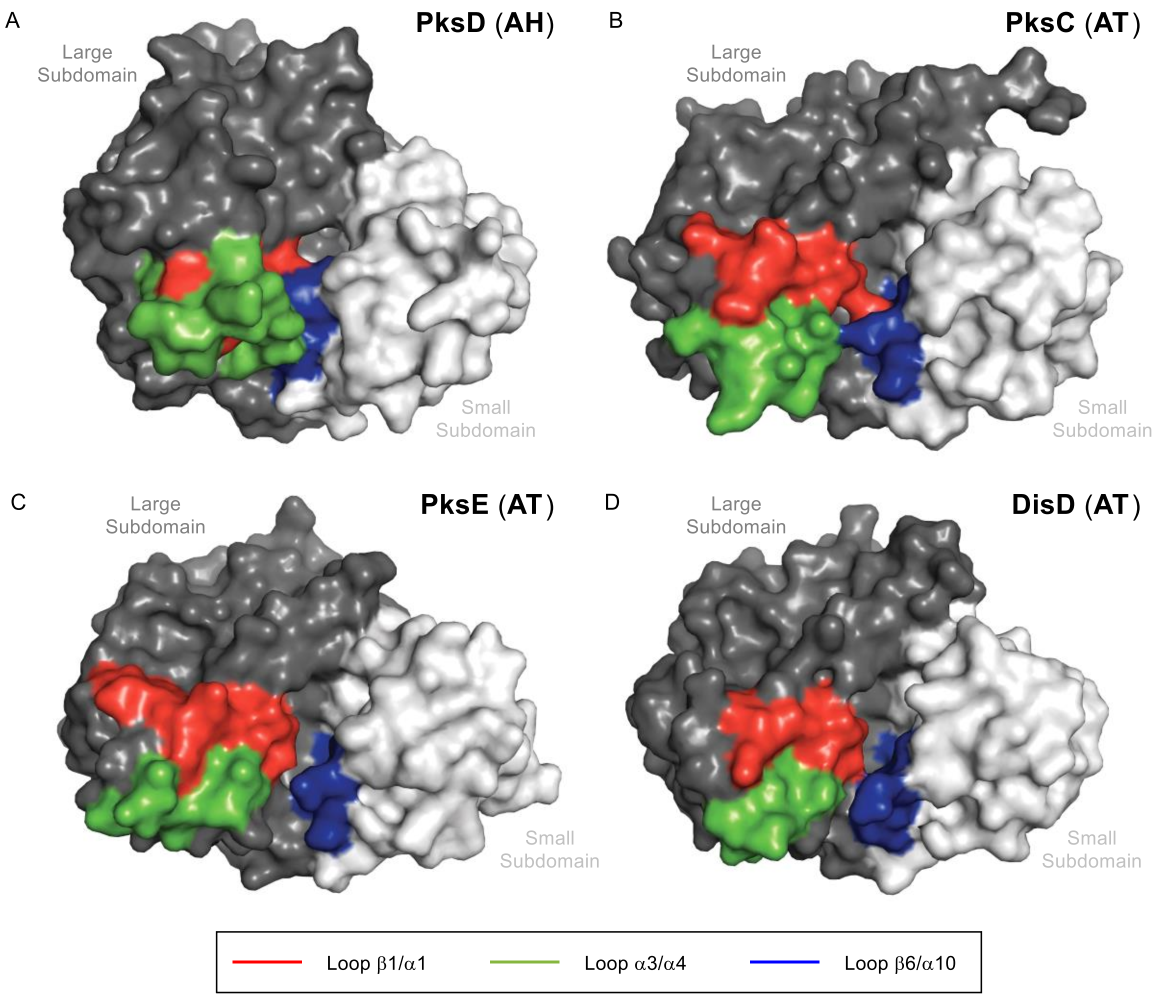

**Figure S12. Structural comparison of binding pocket loops among AH and AT domains (surface rendering).** Juxtaposition of A). PksD (PDB: 8AVZ) with B). PksC (PDB: 5DZ6), C). PksE (PDB: 5DZ7; ER domain not shown) and D). DisD (PDB: 3RGI) reveals a more constricted substrate binding pocket for PksD due to the tighter interactions of loops enveloping the pocket (e.g. see hydrogen bonding network in **Fig. S10**). These loops are named according to bounding secondary structures in PksD. See also **Fig. S11**.

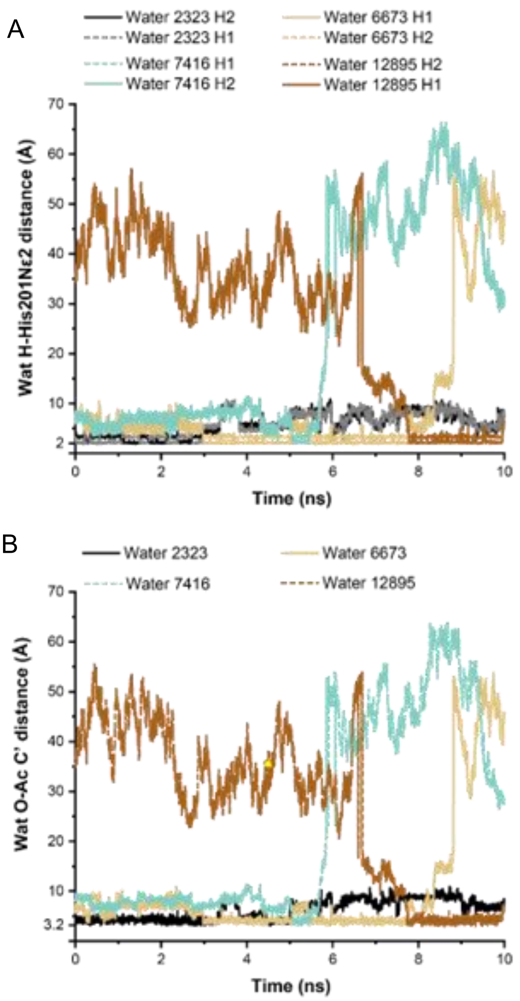

**Figure S13. Recruitment of water molecules to the His201 side chain via H-bonding.** A 10 ns cMD simulation for PksD with acetylated Ser99, restarted from a representative aMD frame (frame 1650) in which water molecules adopt a geometry conducive to nucleophilic attack on the acetyl carbonyl C1 atom. During the simulation, four different water molecules are recruited by the His201 side chain and remain there for up to ~3 ns. A). Distances between water H atoms and His201[Nε2]. B). Distances between water O atoms and the Ser99-Ac carbonyl C1 atom. Water molecules 2323, 5573 and 12895 but not 7416 (with the shortest residence time near His201) meet the criteria for nucleophilic attack in multiple frames.

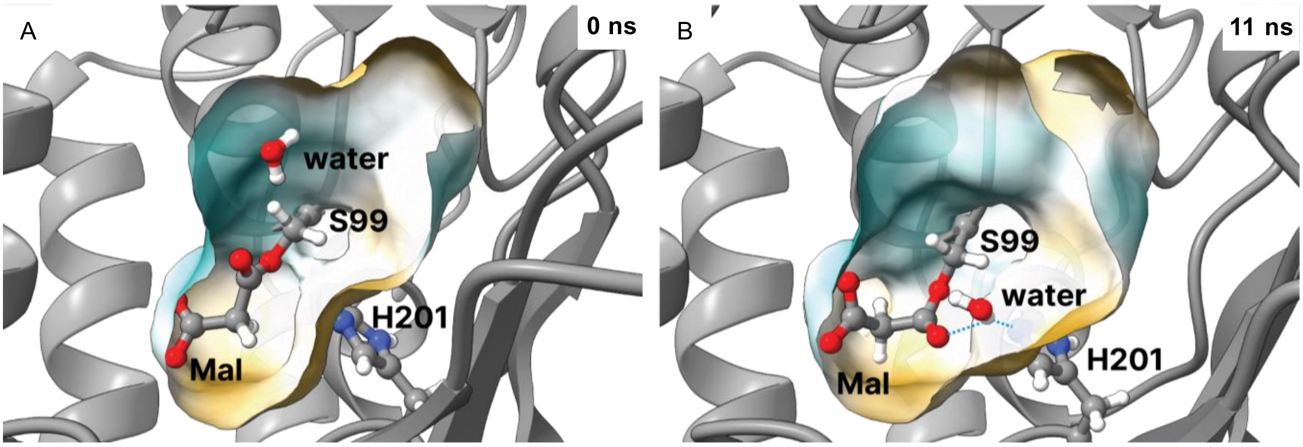

**Figure S14. Representative frames from a 215 ns cMD simulation of malonyl-PksD.** A). Frame at the beginning of the simulation. B). Frame at approximately 11 ns along the trajectory. The starting coordinates for the simulation were taken from the frame in **Fig. 5A** for acetyl-PksD AH, with the acetyl chain manually modified to a malonyl chain. After ~10 ns, the malonyl carboxylate moiety moves away from the hydrophobic part of the pocket, which leads to rotation of the carbonyl moiety with CO bond pointing towards H201. The surface of the pocket is coloured according to hydrophobicity, with yellow and cyan indicating hydrophobic and hydrophilic character, respectively. The water molecule closest to the H201 side chain is also depicted. While some frames with configurations appropriate for hydrolytic attack can be found in the first 6 ns of the trajectory, no such conditions are satisfied after the malonyl group moves away from the hydrophobic part of the pocket (water molecules H-bonded to the H201 side chain are positioned unfavourably with respect to the carbonyl group for nucleophilic attack) (**Video S3**).

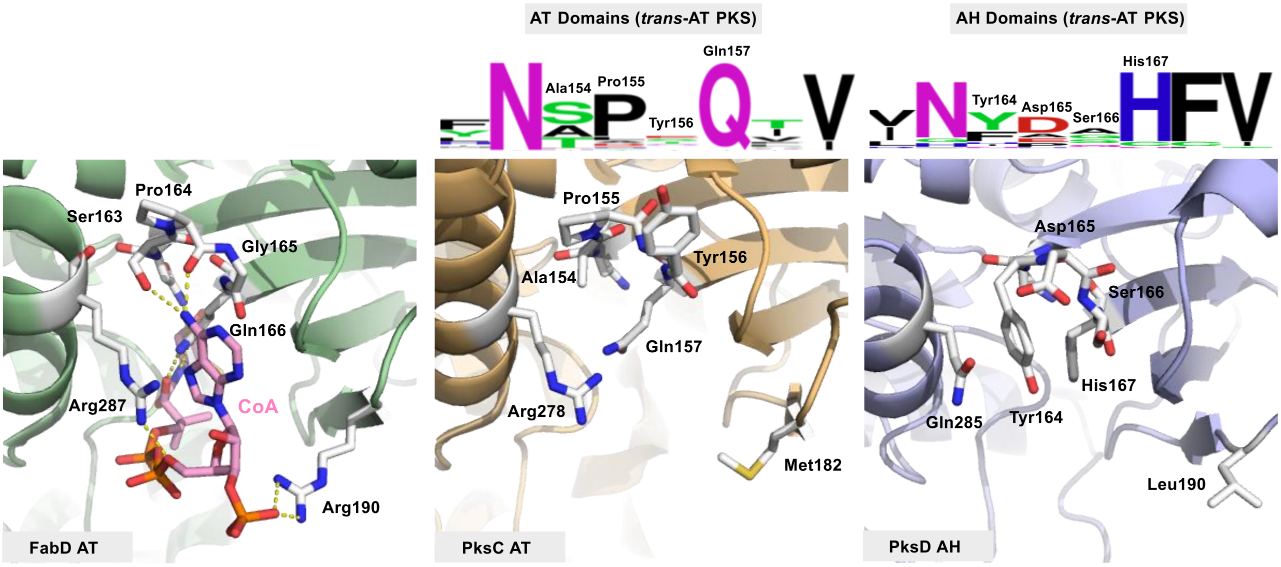

**Figure S15. Comparison of the 3′-phosphoadenosine-binding pocket in AT versus AH domains.** Renderings of the 3′-phosphoadenosine-binding region for FabD+CoA (PDB: 2G2Z), PksC (PDB: 5DZ6) and PksD (PDB: 8AVZ). Consensus sequences of amino acids involved in the binding region are shown for AT and AH domains from *trans*-AT PKS systems, generated by the WebLogo server.[6] The 3′-phosphoadenosine-binding regions for FabD and PksC are very similar, providing an open channel lined with conserved residues that make polar contacts with CoA. The equivalent region in PksD presents an occluded channel, resulting from the presence of Tyr164 (in place of an Ala in PksC). This position is often occupied by a bulky residue (typically Tyr or Phe) in AH domains.

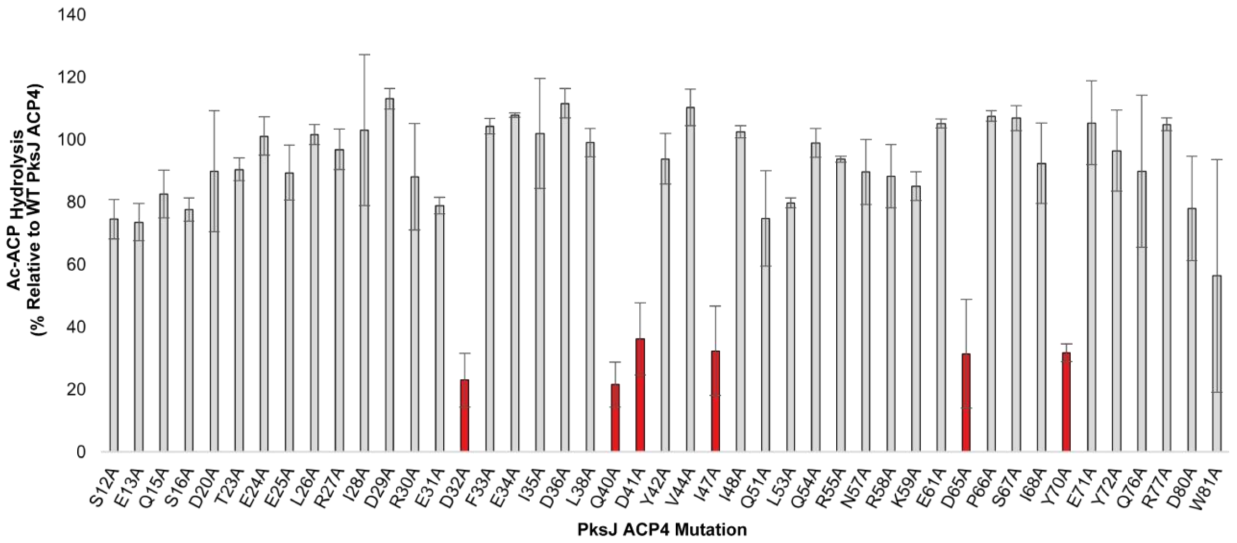

**Figure S16. Effect of alanine scanning mutants of PksJ ACP4 on acetyl hydrolysis.** Profile of PksD-catalysed acetyl group hydrolysis towards scanning alanine mutants of PksJ ACP4, expressed as a percentage of activity relative to WT PksJ ACP4. Mutations that significantly decreased the hydrolytic activity are highlighted in red. Error bars represent the standard deviation of three replicates.

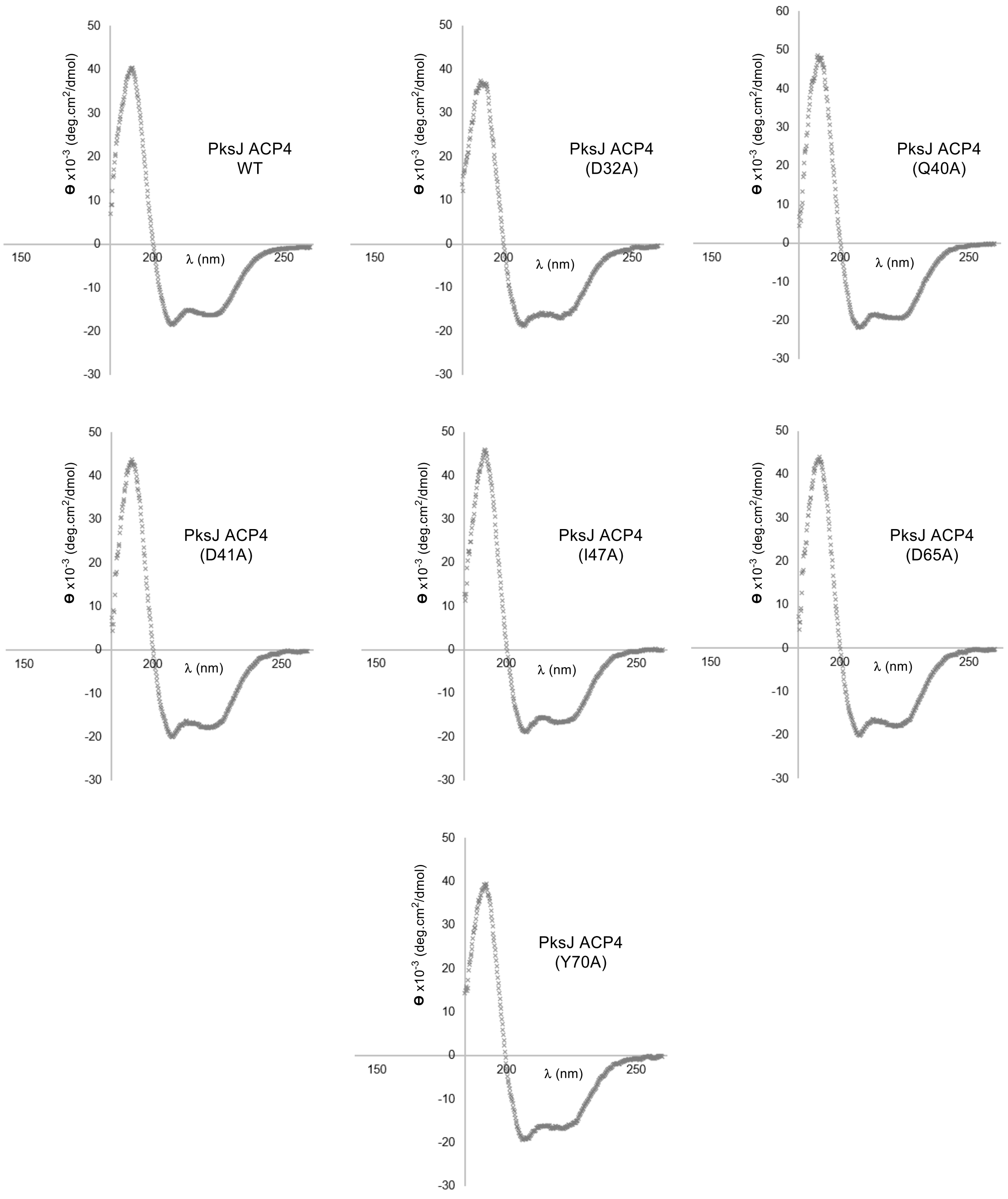

**Figure S17. Circular dichroism (CD) spectroscopic analyses of PksJ ACP4 mutants.** CD spectra for wild-type PksJ ACP4 and all alanine variants that significantly disrupted the interaction with PksD. The CD spectrum obtained for each mutant is in agreement with that of wild-type PksJ ACP4, suggesting that no major gain or loss of secondary structure has occurred upon mutation.

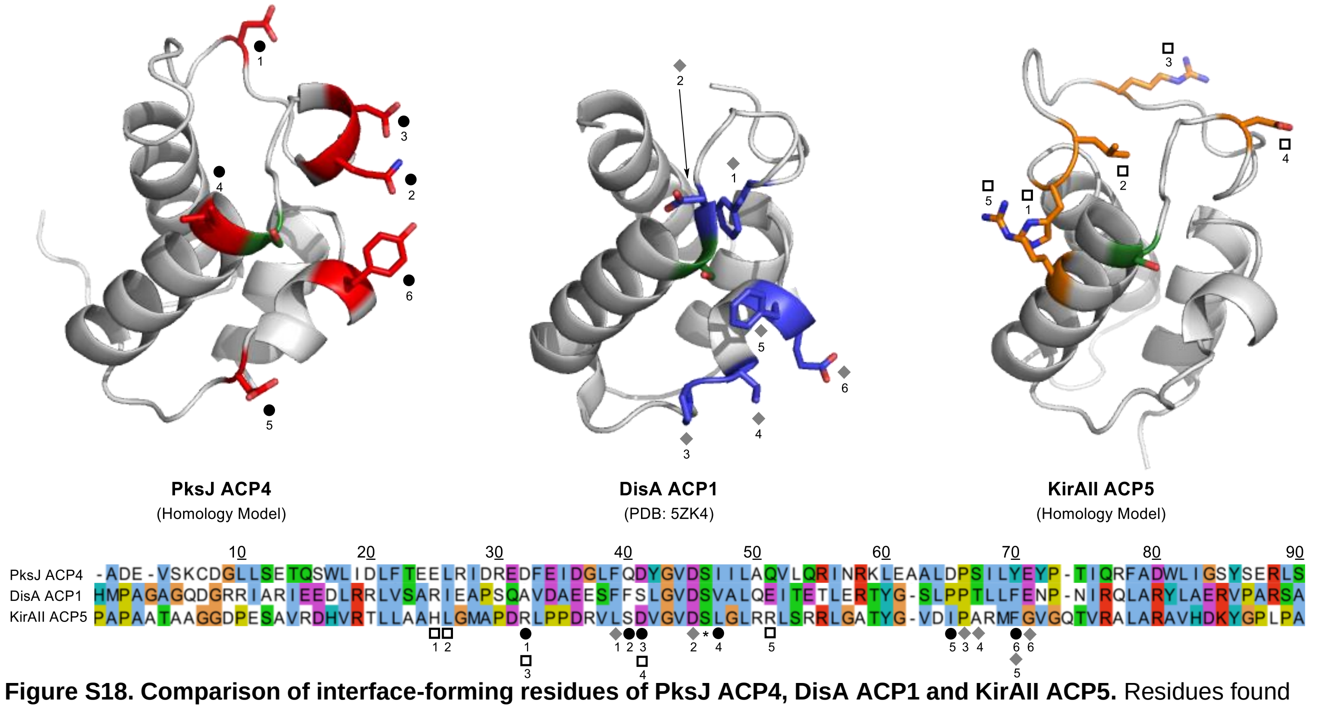

**Figure S18. Comparison of interface-forming residues of PksJ ACP4, DisA ACP1 and KirAII ACP5.** Residues found to be involved in ACP:AH or ACP:AT domain interactions are highlighted and indicated below the sequence alignment as follows: PksD:PksJ ACP4 (black circles), DisD:DisA ACP1 (grey diamonds)[8], KirCII:KirAII ACP5 (white squares)[9]. It is worth noting that KirAII ACP5 interacts exclusively with an ethylmalonyl-loading AT domain (KirCII), distinct from the rest of the ACP domains in the pathway. This may explain the different interaction motif compared to DisA ACP1. Sequences are numbered according to PksJ ACP4. Homology models were generated using the iTASSER server.[10]

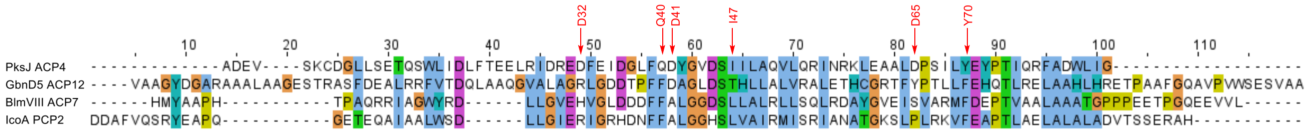

**Figure S19. Sequence alignment of PksJ ACP4 with other carrier protein domains tested experimentally with PksD.** Residues involved in the PksD:PksJ ACP4 interface are highlighted with red arrows. The tested carrier protein domains originate from the following pathways: GbnD5 ACP12 (gladiolin *trans*-AT PKS), BlmVIII ACP7 (bleomycin *cis*-AT PKS-NRPS) and IcoA PCP (icosalide A1 NRPS).

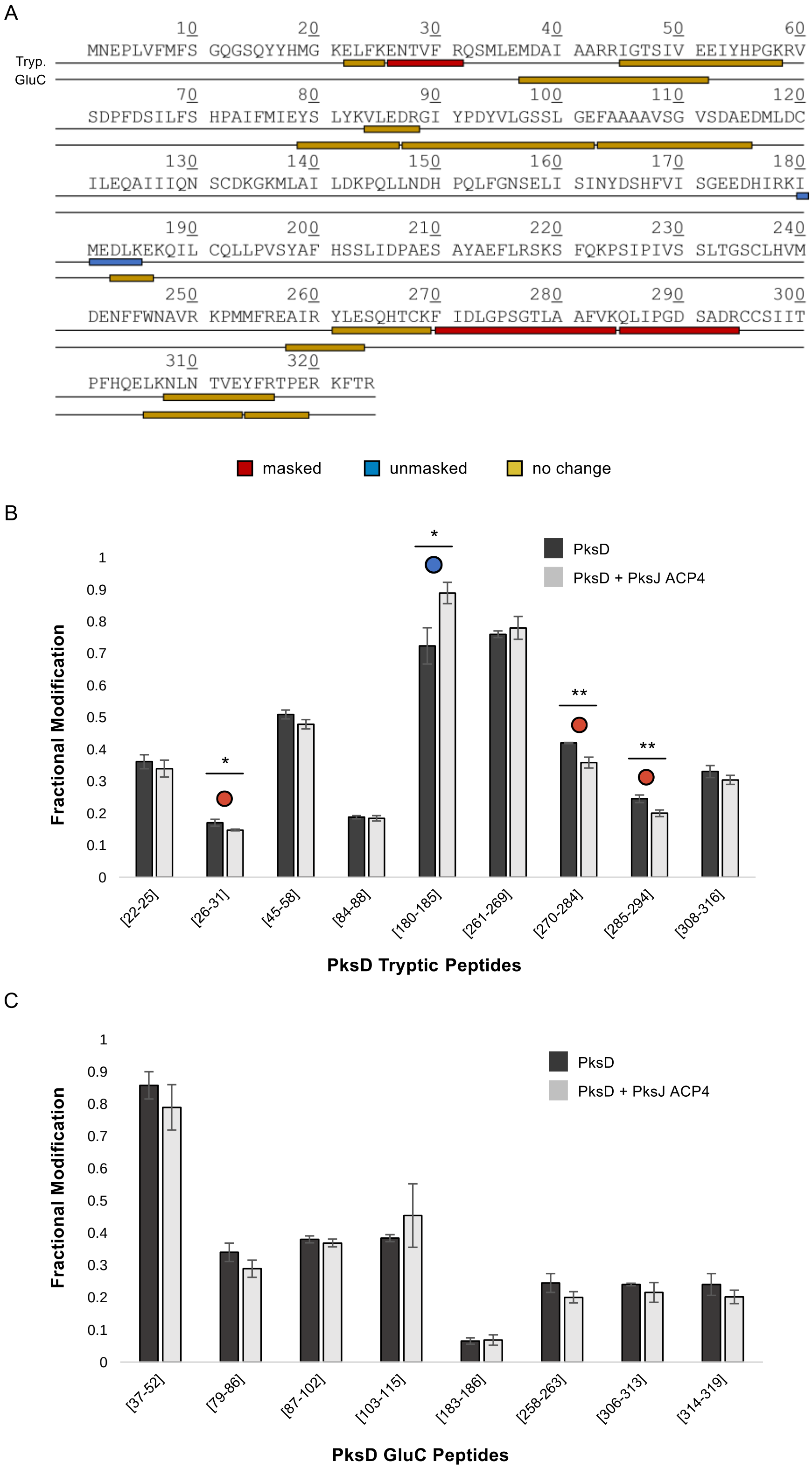

**Figure S20. Carbene footprinting of PksD and *holo*-PksJ ACP4.** A). Combined sequence coverage of PksD using trypsin (Tryp.) and GluC proteases. Masked (red), unmasked (blue) and unchanged (wheat) regions are shown for each peptide. Fractional modification of PksD B). tryptic peptides and C). GluC peptides, in the presence (light grey) and absence (dark grey) of *holo*-PksJ ACP4. Masked (red dot) and unmasked (blue dot) peptides are highlighted; error bars represent standard deviation (n = 3); and the level of significance, calculated with a Student's *t*-test, is denoted with * ($p < 0.05$) or ** ($p < 0.01$).

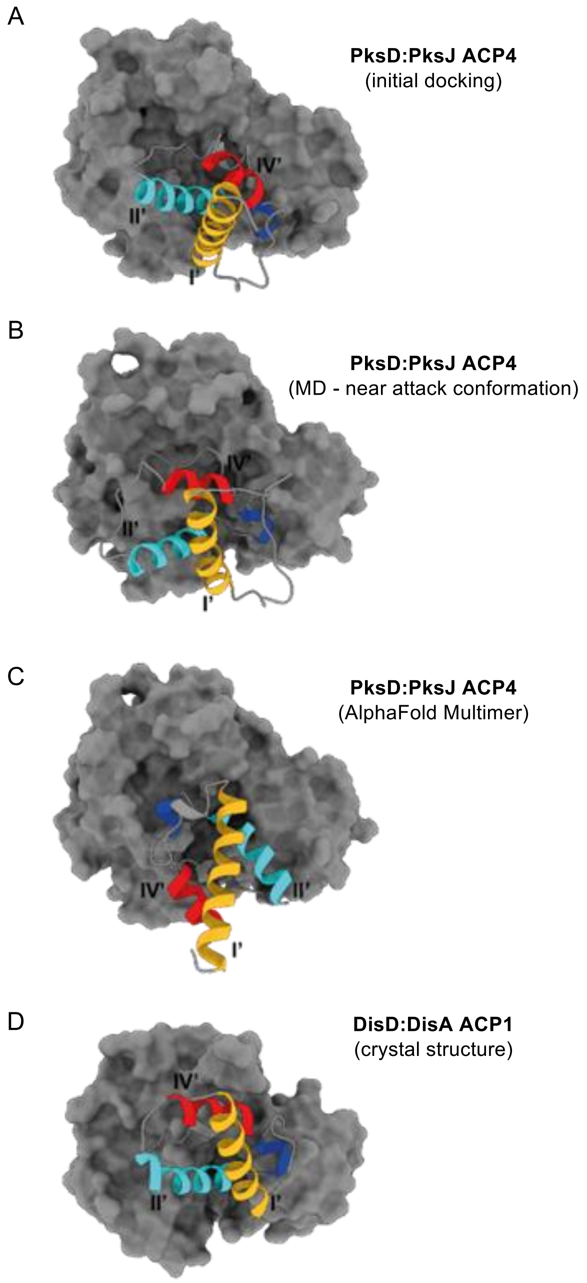

**Figure S21. Comparison of ACP domain orientations when in complex with an AH or AT domain.** A). PksD:PksJ ACP4 complex – initial docking, based on carbene footprinting and alanine scanning mutagenesis data. B). PksD:acetyl-Ppant-PksJ ACP4 complex – representative frame from aMD simulations in which conditions for a near-attack conformation are met. C). PksD:PksJ ACP4 complex – AlphaFold Multimer prediction.[11,12] D). DisD:DisA ACP1 complex – crystal structure (PDB: 5ZK4). All models were oriented through structural alignment to the AH domain. AH and AT domains are rendered as grey surfaces; ACP domain helices are depicted as coils coloured as follows: I (yellow), II (cyan), III (blue) and IV (red).

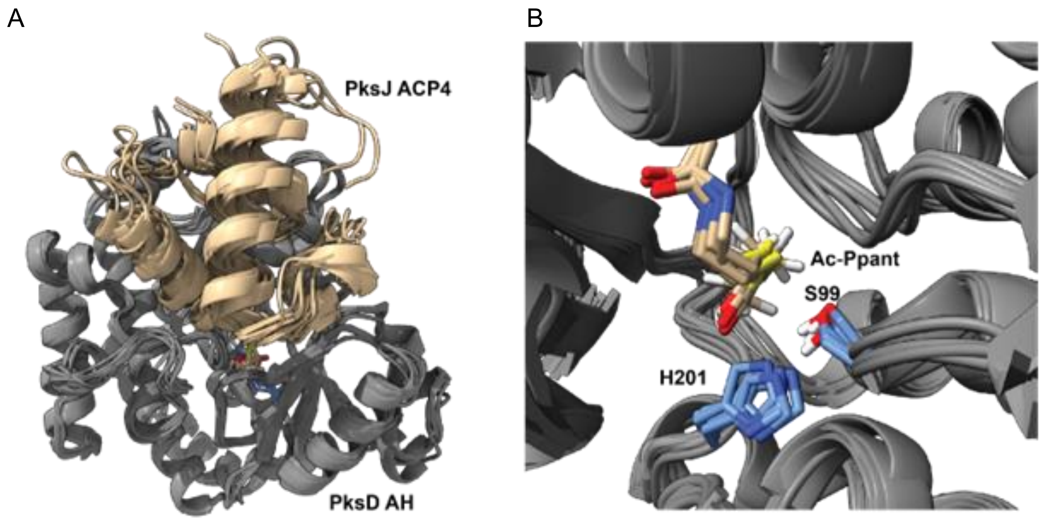

**Figure S22. Overlay of simulation frames matching the geometry for nucleophilic attack of acetyl-Ppant by Ser99.** cMD simulations initiated from aMD frames 3221 and 4230 of the PksD:acetyl-Ppant-PksJ ACP4 complex. All models were oriented through structural alignment to the AH domain. A). Overlay of the positions of the ACP domain (wheat) with respect to the AH domain (grey). B). Overlay showing the relative positions of Ser99 and His201 of PksD (light blue carbons) and the acetyl-Ppant moiety bound to PksJ ACP4 (beige carbons).

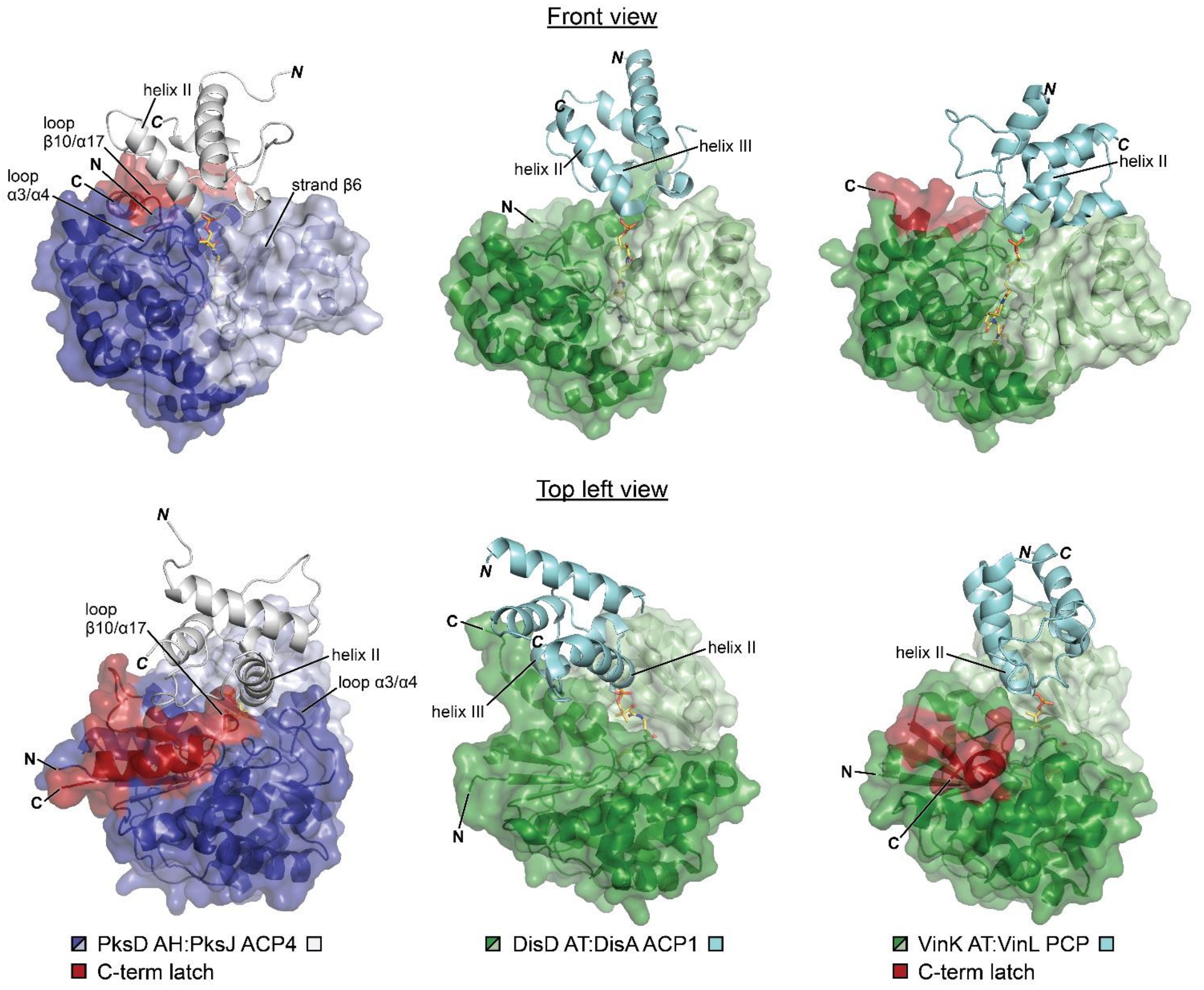

**Figure S23. Binding of carrier protein domains to AH and AT domains from *trans*-AT PKSs.** A representative frame of the PksD:PksJ ACP4 simulated complex is compared to the DisD:DisA ACP1 (PDB: 5ZK4) and VinK:VinL PCP (PDB: 5CZD) crosslinked complexes from two perspectives. The binding mode of VinL PCP to its partner catalytic domain varies considerably from that of PksJ ACP4 and DisA ACP1. AH and AT domains are rendered as cartoons and semi-transparent surfaces; carrier protein domains are rendered as cartoons; catalytic dyad residues and substrates/crosslinkers are shown as sticks coloured by atom. All models were oriented through structural alignment to the AH domain.

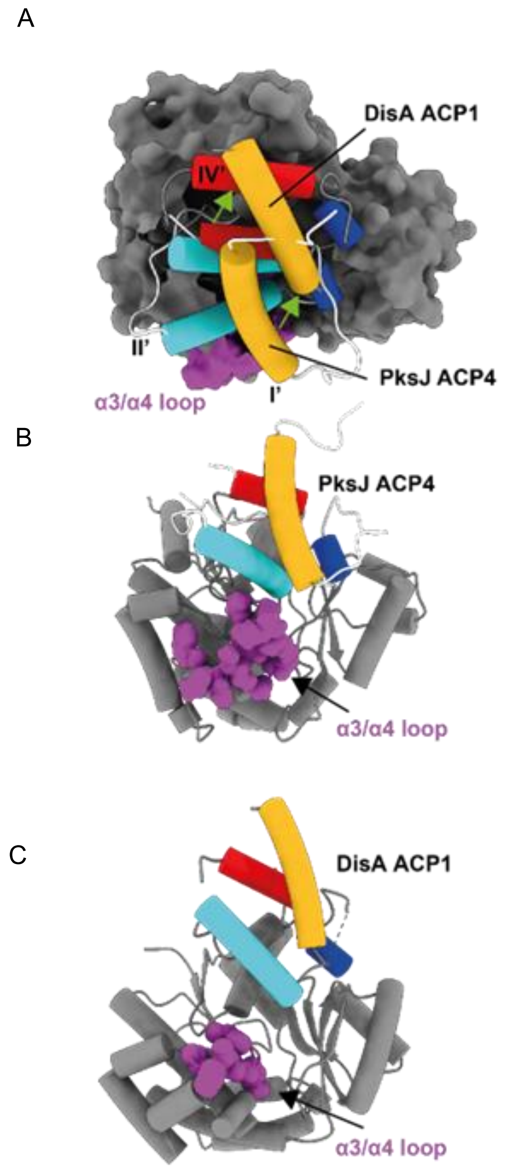

**Figure S24. Comparison of ACP domain positions when in complex with DisD and PksD.** A). Overlay of the complexes, in which models were structurally aligned between AT and AH domains. The AH:ACP domain complex is the same as that depicted in **Fig. 9** (e.g. docked and subjected to MD simulations). DisA ACP1 originates from the crystal structure of the DisD:DisA ACP1 crosslinked complex (PDB: PDB 5ZK4). AH and AT domains are rendered as grey surfaces or cylinders; ACP domain helices are depicted as cylinders coloured as follows: I (yellow), II (cyan), III (blue) and IV (red). The ACP domains assume very similar orientations with respect to their partner catalytic domains but with an overall shift indicated by green arrows. B - C). Position of the α3/α4 loop (magenta) of PksD relative to B). PksJ ACP4 and C). DisA ACP1.

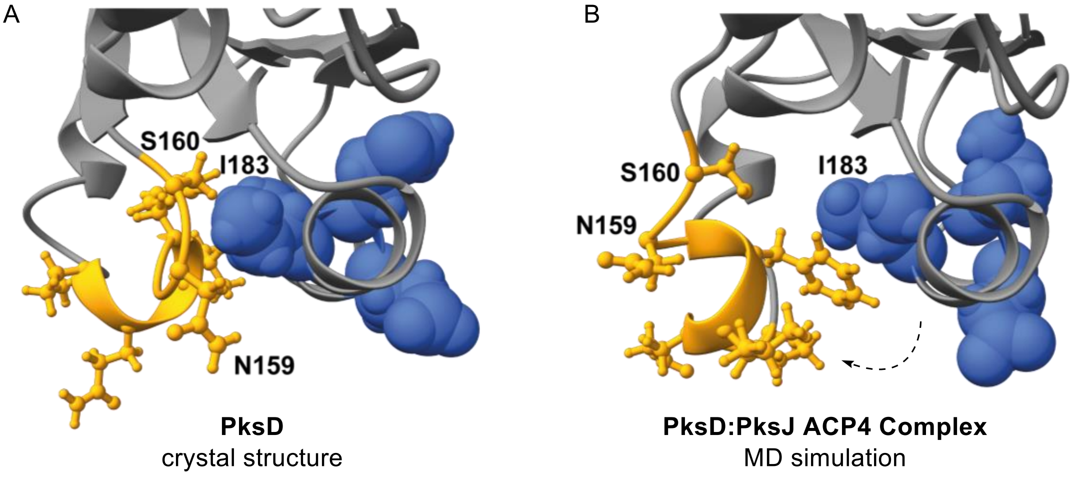

**Figure S25. Example of structural rearrangements in PksD upon ACP domain binding.** A). Crystal structure of PksD focussing on the association between helix α9 (blue space-filled residues) and adjacent loop and $3_{10}$ helix α8 (gold ball-and-stick residues). A close association between residues Ile183 and Asn159/Ser160 is observed. B). Equivalent region of PksD following docking of PksJ ACP4 and MD simulation, arriving at a complex with the required catalytic geometry for the acylation step. Here the loop and $3_{10}$ helix α8 no longer pack against helix α9, resulting in increased solvent exposure. This is reflected in our carbene footprinting data, wherein residues 180-182 become unmasked upon ACP domain binding.

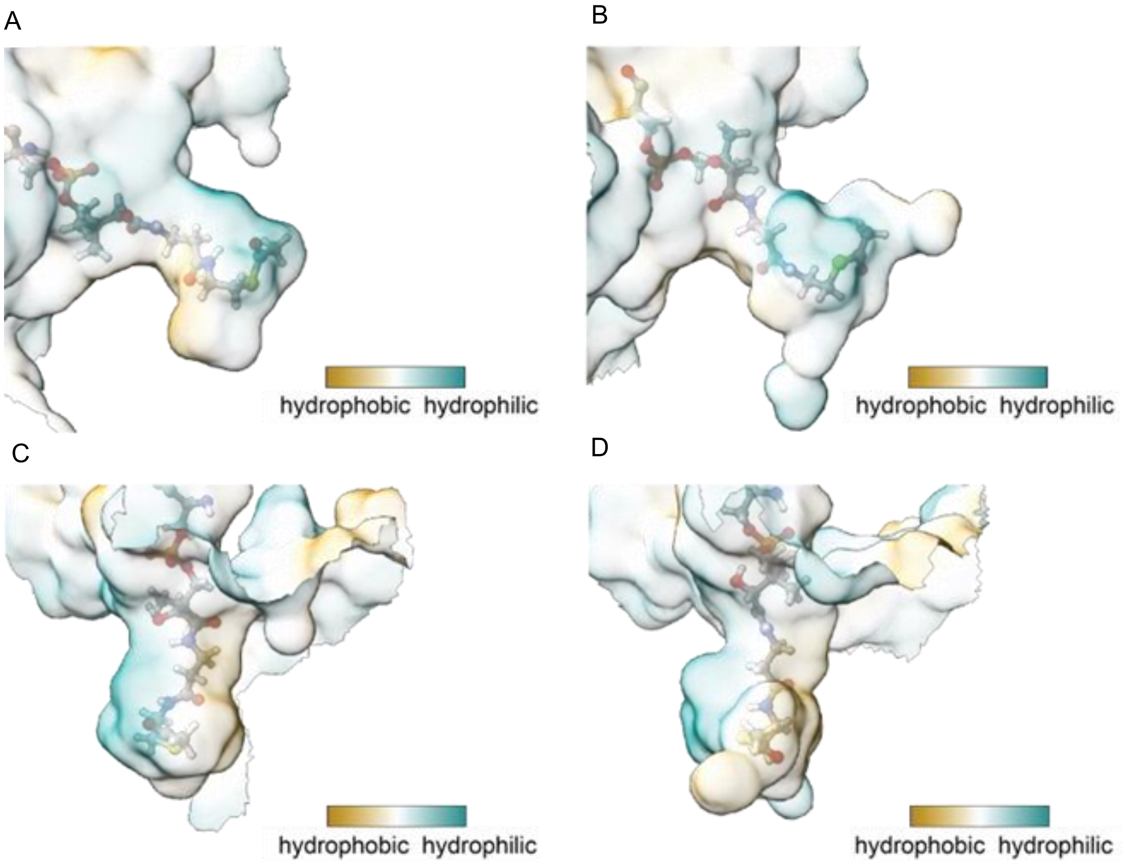

**Figure S26. Representative shapes of the substrate binding pocket of PksD.** A). and C). Different views of the pocket from the starting structure in the MD simulation, i.e. after manual placement of the substrate and energy optimisation of the crystal structure of PksD. B). and D). Different views of the pocket from the near-attack frame depicted in **Fig. 9**. The acetyl-Ppant moiety is shown as ball-and-sticks; the semi-transparent surface of the pocket is coloured according to hydrophobicity. Panels C). and D). highlight the dual hydrophobic and hydrophilic faces of the channel.

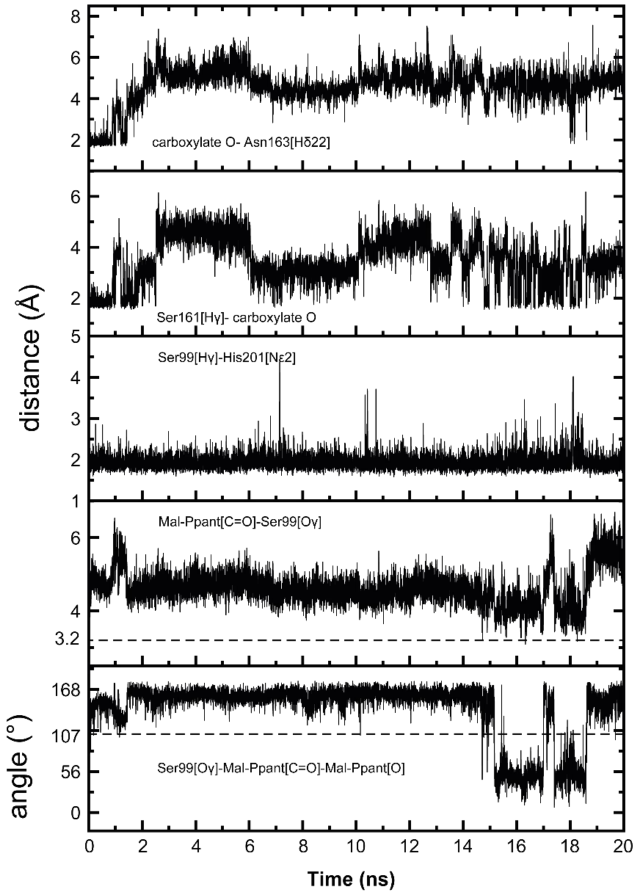

**Figure S27. A 20 ns cMD simulation initiated from an aMD frame of PksD:malonyl-Ppant-PksJ ACP4 in a near attack conformation.** Panels from top to bottom: distance between one of the terminal carboxylate oxygen atoms of malonyl-Ppant to Asn163[Hδ22] (equivalent distance to the other carboxylate oxygen not shown); distance between Ser161[Hγ] and one of the terminal carboxylate oxygen atoms of malonyl-Ppant (distance to the other carboxylate oxygen not shown); distance between Ser161[Hγ] and His201[Nε2]; distance between Ser99[Oγ] and malonyl-Ppant[C1] (dashed line indicates a distance of 3.2 Å); angle between Ser99[Oγ], malonyl-Ppant[C1] and malonyl-Ppant[O] (dashed line indicates an angle of 107°).

### 2. Biochemical, Biophysical and Computational Methods

#### 2.1 Molecular Cloning and Mutagenesis

Amplification of the *PksD* gene from *Bacillus subtilis* (strain 168) gDNA was performed using Q5 DNA polymerase (NEB) and the primers detailed in **Table S1**. PCR products were separated on a 1% agarose gel and bands were excised and purified with a GeneJet gel extraction kit (Thermo Fisher Scientific). The purified insert was digested with the respective restriction enzymes and ligated with XJ-pET28a(+) (pHis$_8$) which had been pre-digested with the complementary restriction enzymes. Ligation reactions proceeded overnight at 25 °C. In all cases, the ligation mixture was used to transform *Escherichia coli* TOP10 cells (Invitrogen), which were plated on LB agar containing kanamycin (50 µg/mL). Individual colonies were picked and grown overnight in LB medium containing (50 µg/mL). Plasmids were isolated from overnight cultures using a GeneJET Plasmid Miniprep Kit (Thermo Fisher Scientific), and inserts were sequenced (Eurofins Genomics) to verify their integrity.

The pET151-GbnD5 ACP12 expression construct was previously reported.[13] The expression construct pET151-IcoA PCP2 was obtained through amplification of relevant region of the *icoA* gene from *Burkholderia gladioli* BCC0238 gDNA using Q5 DNA polymerase (NEB) and the primers detailed in **Table S1**. PCR products were separated on a 1% agarose gel and bands were excised and purified with a GeneJet gel extraction kit (Thermo Fisher Scientific). The insert was ligated with pET151 (Invitrogen) following the manufacturer's instructions. The resulting vector was used to transform *E. coli* TOP10 cells (Invitrogen), which were plated on LB agar containing ampicillin (100 µg/mL). Colonies were picked and grown overnight in LB medium. Plasmids were isolated from the culture using a miniprep kit (Thermo), and the inserts were sequenced to verify their integrity.

In order to obtain BlmVIII ACP7, genomic DNA from the bleomycin producer *Streptomyces verticillus* was isolated using an adapted salting out protocol.[14] TSB medium was inoculated with freeze-dried *S. verticillus* ATCC15003 (DSMZ) and incubated with shaking at 30 °C. The next morning, mycelia were pelleted (1500 rcf, 10 min). Several inoculation loops worth of pelleted mycelia were transferred to 500 mL SET Buffer (75 mM NaCl, 25 mM EDTA pH 8.0, 20 mM Tris pH 7.5). This mixture was incubated with lysozyme (1 mg/mL; Sigma-Aldrich) at 37 °C for 30 min, with regular mixing by inversion. Next, proteinase K (0.5 mg/mL; NEB) and SDS (1%) were added, and the mixture was incubated at 55 °C for 1 h, with regular mixing by inversion. After adding NaCl (1.25 M), DNA was extracted by adding 500 mL chloroform and mixing by inversion. Following centrifugation (15,000 rcf, 10 min), the supernatant was transferred to a separate tube, and a second round of extraction was carried out by adding another 500 mL chloroform and mixing by inversion. After centrifugation (15,000 rcf, 10 min), the supernatant was transferred to a separate tube, and DNA was precipitated by adding isopropanol (0.6 volumes), mixing by inversion, and incubating on ice for 10 min. Following centrifugation (15,000 rcf, 10 min, 4 °C), the supernatant was discarded, and the sample was washed by adding 1 mL ethanol (70%) and mixing by inversion. Following another round of centrifugation (15,000 rcf, 4 °C, 10 min), the supernatant was discarded and the sample was air dried at room temperature overnight. Residual DNA was resuspended in 50 µL mQ$H_2O$. Gene fragment inserts and vector backbones were PCR-amplified from the purified gDNA and intact pET28a, respectively, using Phusion High-Fidelity DNA Polymerase (NEB) and primers (Sigma-Aldrich) listed in **Table S1**. Amplified DNA was purified by agarose gel electrophoresis, then gel-extracted and cleaned up with a QIAquick Gel Extraction Kit (Qiagen). Cloning was accomplished by Gibson assembly in accordance with the manufacturer's protocol (NEB). The product was transformed into TOP10 *E. coli* cells, selected for on LB agar supplemented with 50 µg/mL kanamycin and isolated from a single transformant using a QIAprep Spin Miniprep Kit (Qiagen). Sequence integrity was confirmed by DNA sequencing (Eurofins Genomics). The correctly sequenced plasmid was then transformed into *E. coli* BL21(DE3) for heterologous expression and subsequent protein purification, as performed above.

PksD(S98A), PksD(S98H), PksD(S99A), PksD(H201N), and PksD(Q124A) were produced using the Q5 site-directed mutagenesis kit (NEB), with the wild-type construct as a template and the primers detailed in **Table S1**. PCR products were processed according to the manufacturer's protocol, and resulting plasmids sequenced (Eurofins Genomics) to verify the success of mutagenesis. The library of PksJ ACP4 X→Ala mutants was reported previously.[15]

### 2.2 Protein Overproduction and Purification

#### *2.2.1 Native PksD, PksJ ACP4 and Associated Mutants*

A single colony of *E. coli* BL21(DE3) cells that had been transformed with the appropriate expression vector was picked and used to inoculate LB medium (5 or 10 mL) containing kanamycin (50 µg/mL). The resulting culture was incubated overnight at 37 °C and 180 rpm, then used to inoculate LB medium (0.5 or 1 L) containing kanamycin (50 µg/mL). The resulting culture was incubated at 37 °C and 180 rpm until the optical density of the culture at 595 nm reached 0.6, whereupon IPTG (1 mM) was added, and the temperature was reduced to 15 °C. After overnight incubation, cells were harvested by centrifugation (4000 rcf, 15 min, 4 °C) and resuspended in buffer (20 mM Tris-HCl, 100 mM NaCl, 20 mM imidazole, pH 7.4) at 10 mL/L of growth medium. Cell lysis using an MC Cell Disrupter (Constant Systems Limited) followed.

The lysate was centrifuged (37,000 rcf, 30 min, 4 °C) and the resulting supernatant was loaded onto a HiTrap FF Chelating Column (GE Healthcare Life Sciences), which had been pre-loaded with 100 mM $NiSO_4$ and equilibrated in resuspension buffer (20 mM Tris-HCl, 100 mM NaCl, 20 mM imidazole, pH 7.4). Proteins were eluted in a stepwise manner with resuspension buffer containing increasing concentrations of imidazole – 50 mM (5 mL), 100 mM (3 mL), 200 mM (3 mL) and 300 mM (3 mL). The presence of the protein of interest in elution fractions was confirmed by SDS-PAGE. Where necessary, an additional gel filtration step (HiLoad 16/600 Superdex 75 pg or 200 pg; GE Healthcare Life Sciences) was used to further purify proteins by means of an ÄKTA Pure Protein Purification System (GE Healthcare Life Sciences). Fractions containing the protein of interest were pooled and concentrated to 250 - 400 µM using a Vivaspin 20 Ultrafiltration Unit (Sartorius) at an appropriate molecular weight cut-off. Samples were snap-frozen in liquid $N_2$ and stored at -80 °C until further use.

#### *2.2.2 SeMet-Derived PksD*

For preparation of SeMet-derived protein, the plasmid encoding PksD was transformed into methionine-auxotrophic *E. coli* B834(DE3) cells and selected for on LB agar supplemented with kanamycin (50 µg/mL). From a single colony, an overnight culture was prepared in LB medium containing kanamycin with shaking at 180 rpm and 37 °C. This culture was pelleted (2000 rcf, 5 min, 4 °C) and resuspended in M9 minimal media supplemented with kanamycin, vitamins and trace elements (**Tables S2** and **S3**), nineteen of the twenty proteinogenic amino acids (40 mg/L culture; excluding methionine), and selenomethionine (50 mg/L culture). Once the culture reached an optical density at 595 nm of 0.6 - 0.8, the temperature was decreased to 15 °C and amino acids were added again as previously. After 15 minutes, protein expression was induced through addition of 1 mM IPTG.

After 16 - 20 h, cells were harvested (5000 rcf, 20 min, 4 °C) and resuspended in Ni-NTA Buffer (30 mM HEPES pH 7.5, 500 mM NaCl, 10% glycerol). Cells were incubated with DNase I (Sigma-Aldrich) and hen egg white lysozyme (Alfa Aesar) on ice for 30 min and then lysed using a Vibra-Cell Ultrasonic Liquid Processor (Sonics & Materials, Inc). The lysate was clarified by centrifugation (30,000 rcf, 1 h, 4 °C), after which the supernatant was filtered (0.45 µm cellulose acetate membrane; Sartorius Stedim Biotech) and loaded onto a 1 mL HisTrap HP column (GE Healthcare Life Sciences) equilibrated with Ni-NTA Buffer for affinity chromatography by means of an ÄKTA Pure Protein Purification System (GE Healthcare Life Sciences). Washing and elution were accomplished by flowing Ni-NTA Buffer containing 10 and 150 mM imidazole, respectively, over the column. Fractions containing pure protein, as identified by SDS-PAGE, were concentrated and exchanged into Ni-NTA Buffer in an Amicon Ultra-15 Centrifugal Filter Unit (30 kDa MWCO; EMD Millipore).

### 2.3 Crystallisation and Structure Determination of PksD

Prior to crystallisation of PksD, the N-terminal His-tag was cleaved by incubating the protein with bovine thrombin (GE Healthcare Life Sciences) overnight at 4 °C. Completion of cleavage was confirmed by an SDS-PAGE gel shift. Through means of an ÄKTA Pure Protein Purification System (GE Healthcare Life Sciences), thrombin was removed by passing the cleavage reaction over a column containing 1 mL Benzamidine Sepharose 4 Fast Flow (GE Healthcare Life Sciences) equilibrated with Ni-NTA Buffer. Elution fractions were concentrated as above and further polished by size exclusion chromatography on a HiLoad 16/600 Superdex 200 pg column (GE Healthcare Life Sciences) equilibrated with SEC Buffer (50 mM Tris, 100 mM NaCl, pH 8.0) using an ÄKTA Pure Protein Purification System. Pure fractions, as identified by SDS-PAGE, were concentrated as above to ~11 mg/mL (assuming MW = 37,095 g/mol and $\varepsilon$ = 23,380 $M^{-1}cm^{-1}$)[16] and subjected sparse-matrix crystal screening at 20 °C. Native crystals, grown by vapour diffusion, were optimised in hanging drops containing 0.5 - 1.5 µL crystallisation solution (0.1 M Tris pH 8.5, 6 - 11% (w/v) PEG 8000) and 0.5 - 1.5

μL protein solution. Just before flash-freezing in $LN_2$, crystallisation solution containing 35% glycerol was pipetted directly onto the crystal drop; crystals were then passed through a fresh drop of the same cryosolution.

SeMet-derived crystals, which failed to grow under screening conditions, were generated using Phil Jeffrey's streak seeding protocol.[17] Briefly, a drop containing native PksD crystals was transferred to an Eppendorf tube containing 100 μL of reservoir solution and vortexed for 30 s. Serial dilutions of 1:10 - 1:1000 of this seed stock were also prepared in reservoir solution. Crystal drops were set up as for native protein and allowed to equilibrate for 2 - 3 h at 20 °C. Following equilibration, hanging drop cover slips were removed, a cat whisker was dipped into one of the seed stocks and dragged through the crystal drop and the cover slip was replaced. Crystals were allowed to further mature at 20 °C.

Native and single-wavelength anomalous dispersion datasets were collected from single crystals at 100 K at Diamond Light Source Beamlines I03 and I04-1 and wavelengths of 0.97628 Å and 0.91188 Å, respectively. Reflections were processed in XDS[18,19] and scaled in Aimless[20] in the CCP4 software suite,[21] with free-R flags added to 5% of reflections. Whilst searching for twelve NCS copies of PksD, AutoSol[22] of the PHENIX software suite[23] located 73 Se atoms in a *C*2 unit cell with the following parameters: number of refined sites = 73, FOM = 0.416, overall score = 58.42 +/- 6.99, map skew = 0.25, correlation of local RMS density = 0.90. AutoBuild[24] of the PHENIX software suite further constructed 690 waters and 1253 residues in five fragments with map-model CC = 0.77, $R_{work}/R_{free}$ = 0.1948/0.2173. Several rounds of iterative refinement were carried out on four SeMet-derived copies of PksD in Coot,[25] Refmac5[26] of the CCP4 software suite, and phenix.refine[27] of the PHENIX software suite. To phase native reflections, Phaser MR[28] of the CCP4 software suite was employed to search a *C*2 unit cell for eight copies of a SeMet-derived monomer.

Both structures were subjected to further rounds of iterative refinement as above, with data initially scaled to a mean $I/\sigma(I)$ of 2.0 in the outermost shell with Aimless. Resolution was later extended by rescaling in Aimless, with limits substantiated using the paired refinement[29,30] feature of the PDB-REDO server.[31] Throughout refinement, the MolProbity server[32] was consulted for structure validation. X-ray fluorescence experiments for the purpose of identifying $Zn^{2+}$ were conducted on native crystals at 100 K at Diamond Light Source Beamline I04. All structural figures were prepared in the PyMOL Molecular Graphics System (Version 1.3, Schrödinger, LLC) or ChimeraX.[33–35] Atomic coordinates and structure factors for SeMet-derived and native PksD were deposited in the Protein Data Bank (PDB) under accession codes 8AVZ and 8AW0, respectively.

#### 2.4 UHPLC-ESI-Q-TOF-MS Analysis of Intact Proteins

All intact protein mass spectrometry analyses were conducted on a Bruker MaXis II ESI-Q-TOF-MS connected to a Dionex 3000 RS UHPLC fitted with an ACE C4-300 RP column (100 x 2.1 mm, 5 μm, 30 °C). The column was eluted with a linear gradient of 5 - 100% MeCN containing 0.1% formic acid over 30 min. The mass spectrometer was operated in positive ion mode with a scan range of 200 - 3000 *m/z*. Source conditions were: end plate offset at -500 V; capillary at -4500 V; nebulizer gas ($N_2$) at 1.8 bar; dry gas ($N_2$) at 9.0 L $min^{-1}$; dry temperature at 200 °C. Ion transfer conditions were: ion funnel RF at 400 Vpp; multiple RF at 200 Vpp; quadrupole low mass at 200 *m/z*; collision energy at 8.0 eV; collision RF at 2000 Vpp; transfer time at 110.0 μs; pre-pulse storage time at 10.0 μs.

#### 2.5 Acyl Chain Hydrolysis Assays

PksJ ACP4 or associated alanine mutants (200 μM) were converted to their acetyl-, butyryl-, hexanoyl-, octanoyl-, β-hydroxybutyryl- or malonyl-PksJ ACP4 forms by incubation in 20 mM Tris, 100 mM NaCl, 2 μM Sfp, 1 mM acyl-CoA (Sigma-Aldrich) and 10 mM $MgCl_2$ in a total volume of 50 μL for 30 min at room temperature. Acyl chain hydrolysis assays were performed in 30 μL total volume of 20 mM Tris, 100 mM NaCl. The resulting acyl-ACP species (50 μM) were incubated with PksD (0.05 μM) for 5 min at room temperature, followed by quenching via addition of formic acid to a final concentration of 1% v/v. Samples were then diluted five-fold with $mQH_2O$, then analysed by UHPLC-ESI-Q-TOF-MS.

#### 2.6 Energy Decomposition Analysis of PksD with Acetyl and Malonyl Groups

Three systems were considered: unmodified PksD (no acetyl / malonyl group bound to Ser99), acetyl-PksD and malonyl-PksD. Non-standard acetyl and malonyl groups were parameterized through a Gaussian optimisation at the HF/6-31G(d) level to obtain RESP charges. The R.E.D. server[36] was used to obtain the final parameter files for both acetyl and malonyl groups. Protonation states of all ionizable amino acids were determined using ProPKA,[37] and clashes resolved using MolProbity.[32] The LEaP module[38] of AMBER18[39] was used to neutralise the systems with potassium counterions and to solvate in a cubic box with a minimum distance between the edge of the protein and the box of 10 Å using the ff14SB[35] and TIP3P force fields. All molecular dynamics (MD) simulations were carried out with the AMBER18 pmemd.cuda program.[39] All systems were initially subjected to minimisation of the solvation box applying a restraint of 500 kcal $mol^{-1}$ $Å^2$ on all protein and substrate atoms for 100 cycles using steepest descent followed by 50 cycles with conjugate gradient. Following the minimisation, each of the systems was heated at constant volume, and the protein was restrained with a force constant of 500 kcal $mol^{-1}$ $Å^2$. The heating was done using Langevin dynamics[35] by gradually increasing the temperature by 50 K in 10,000 MD step intervals, until a total of 100,000 MD steps had been completed, resulting in a final temperature of 300 K. Subsequently, the systems were equilibrated using the NVT ensemble with a 1 fs time step reducing the restraints on protein (see **Table S6**). Production consisted of 250 ns runs using the NPT ensemble with a 2 fs time step in triplicate without restraint, which resulted a total of 750 ns for each system. The temperature and pressure were held constant at 300 K and 1.0 bar using Langevin thermostat and Berendsen barostat.[40,41] The bonds with hydrogen atoms were treated with SHAKE.[42] The smooth Particle-Mesh Ewald method was used to treat long-range electrostatic interactions and Isotropic Periodic Sum method was used to treat Van der Waals interactions using a cutoff of 10 Å.[43,44] A FORTRAN90 program was used for the energy decomposition analysis (EDA)[45,46] to calculate non-bonded interactions, namely Coulombic and Van der Waals interaction energies between Ser99 and the rest of the protein. R and Excel were used for statistical analysis. The unmodified PksD structure was considered as the reference and the total difference in interaction energy was calculated between this and the acetyl- / malonyl-modified structures.

#### 2.7 Circular Dichroism Analyses of ACP Domain Variants

Circular dichroism spectra of PksJ ACP4 mutants (0.1 mg/mL, dissolved in 0.5 mM Tris HCl, 5 mM NaCl, pH 7.4) were recorded between 190 and 260 nm at room temperature on a JASCO J-1500 CD Spectrophotometer using a 1 mm pathlength quartz cuvette. Raw spectral data were extracted and plotted in Microsoft Excel.

#### 2.8 Carbene Footprinting of PksD:PksJ ACP4 Complex

Photolysis experiments were carried out using either PksD (50 µM), *holo*-PksJ ACP4 (50 µM) or PksD:PksJ ACP4 complex (50 µM) in 20 mM Tris, 100 mM NaCl, 10 mM 4-(3-trifluoromethyl)-3H-diazirin-3-yl)benzoic acid (Sigma-Aldrich), pH 7.4 (total volume of 20 µL). The mixture was left to equilibrate for 5 min at room temperature before 6 µL aliquots were placed in crystal clear vials (Fisher Scientific UK) and snap-frozen in liquid nitrogen.

The labelling reaction was initiated by photolysis of the mixture using the third harmonic of a Nd:YLF laser (Spectra Physics, repetition frequency 1000 Hz, pulse energy 130 µJ) at a wavelength of 347 nm. The frozen samples were irradiated for 16 s. All experiments were performed in triplicate. Following irradiation, samples were thawed, reduced (10 mM DTT, in 10 mM ammonium bicarbonate), alkylated (55 mM iodoacetamide, in 10 mM ammonium bicarbonate) and incubated with trypsin or GluC at 37 °C overnight (1:20 protease/protein ratio in 10 mM ammonium bicarbonate).

The analysis of the digests was carried out on a Bruker MaXis II ESI-Q-TOF-MS connected to a Dionex 3000 RS UHPLC fitted with an ACE C18 column (100 x 2.1 mm, 5 µm, 30 °C). The column was eluted with a linear gradient of 5 - 100% MeCN containing 0.1% formic acid over 40 min. The mass spectrometer was operated in positive ion mode with a scan range of 200 - 3000 *m/z*. Source conditions were: end plate offset at -500 V; capillary at -4500 V; nebulizer gas ($N_2$) at 1.6 bar; dry gas ($N_2$) at 8 L $min^{-1}$; dry temperature at 180 °C. Ion transfer conditions were: ion funnel RF at 200 Vpp; multiple RF at 200 Vpp; quadrupole low mass at 55 *m/z*; collision energy at 5.0 eV; collision RF at 600 Vpp; ion cooler RF at 50-350 Vpp; transfer time at 121 s; pre-pulse storage time at 1 µs. A previously described method was used to quantitate the fraction of each peptide modified.[47] Briefly, the chromatograms for each singly-labelled and unlabelled peptide were extracted within a range of ± 0.05 *m/z* and the spectrum for each peak was manually inspected to ensure the sampling of the correct ion only. The peptide fractional modification was calculated using **Equation 1**.

**Equation 1:**

$$P = \frac{A_{labelled}}{A_{labelled} + A_{unlabelled}}$$

where $A_{labelled}$ and $A_{unlabelled}$ correspond, respectively, to the peak area of each labelled and unlabelled peptide. Differences in the extent of labelling between peptides were considered significant when the *p*-value obtained from a Student's *t*-test was $< 0.05$.

### 2.10 Bioinformatics Processing of AH and AT Sequences

Amino acid sequences for AH and AT domains from *trans*-AT PKS pathways were extracted from the MIBiG repository.[2] Multiple sequence alignments were conducted using Clustal Omega[3] on the EMBL-EBI server,[4] and all alignments were processed and visualised in JalView.[5] Logos for consensus sequences were constructed using the WebLogo server.[48]

### 2.11 Molecular Modelling, QM/MM and Docking Simulations

#### *2.11.1 Molecular Simulations of acetyl- / hexanoyl- / malonyl-Modified forms of PksD*

For simulations of PksD modified with Ac / Hex / Mal groups, the Ser99 was manually appended with an Ac group using virtual reality (VR) in ChimeraX[33–35] to Ser99 from chain A from the crystal structure, such that C2 of the respective acyl chains pointed into the substrate channel (note: the carbonyl group was not rotated towards the predicted oxyanion hole). The AMBER ff19SB force field[49] was used for standard residues and parametrised serine derivatives (*O*-acetyl-Ser99, *O*-malonyl-Ser99 and *O*-hexanoyl-Ser99) with ANTECHAMBER[50] and GAFF force field.[51] The charge states of the amino acids at pH 7.4 were evaluated using the H++ webserver.[52–54] The initial structures for the simulations were neutralised with $Na^+$ ions and solvated with OPCBOX water,[55] such that no protein atoms were positioned < 10 Å from any box edge, using the TLEAP program. Additional $Na^+$ and $Cl^-$ ions were added to obtain 100 - 150 mM final salt concentration. MD heating, equilibration, and production steps were performed using the GPU-accelerated[56,57] AMBER 20[58] software on a local workstation equipped with Nvidia GeForce 2080 Ti graphics cards or HPC cluster equipped with Nvidia RTX 6000 graphics cards. Simulation used the SHAKE algorithm[42] to constrain all protein bonds involving a hydrogen atom; a 1.0 - 2.0 fs time-step was used in these simulations. Long-range electrostatics were calculated using the Particle Mesh Ewald (PME) method[44,59] with a 12.0 Å cutoff. PME was used for nonbonded interactions. In all simulations, the Langevin thermostat ($\gamma$ = 2.0 $ps^{-1}$) was used to maintain temperature control.[60] The solvated protein was then equilibrated by carrying out a short minimisation, 50 ps of heating and 50 ps of density equilibration with weak restraints on the protein followed by 500 ps of constant pressure equilibration at 300 K. More specifically, after a two-step minimization process, in which solvent molecules were allowed to relax before the entire system was minimized, the system was slowly heated to 300 K over 0.1 ns in a canonical ensemble (NVT) simulation, then equilibrated for 2 ns by performing isothermal-isobaric (NPT) simulations at 300 K using a Berendsen barostat.[41]

WT PksD AH with *O*-acetyl-Ser99 was initially simulated for 50 ns of cMD followed by ~320 ns of aMD saved every 20 ps. The aMD parameters (estimated from the cMD simulation) were: a) EthreshP = -171521.1 kcal $mol^{-1}$; b) alphaP = 11630.8 kcal $mol^{-1}$; c) EthreshD = 2856.5 kcal $mol^{-1}$; d) alphaD = 224 kcal $mol^{-1}$. For all the aMD simulations, long-range electrostatics were calculated using the PME method with a 10.0 Å cutoff. 10 ns cMD simulations were restarted from frames 1650 and 2183 of the aMD simulations, reading in only the coordinates and not the velocities. Data shown in the figures are from the cMD simulation restarted from the frame 1650. To identify frames which fulfilled the Bürgi-Dunitz criteria, the trajectories were filtered using 3.2 Å and 3.0 Å cutoffs for water O atom-to-Ac carbonyl C1 atom distance and water H atom-to-H201Nε2 distance, respectively, and 100 - 110° angle between water O atom and Ac carbonyl C1 and O atoms. The initial coordinates for the simulation of the WT PksD with *O*-malonyl-Ser99 were taken from the frame depicted in **Fig. 5A**, with the Mal chain inserted manually in VR. The system was simulated for 50 ns of cMD. The initial coordinates for the simulation of the WT PksD with *O*-hexanoyl-Ser99 were taken from the frame depicted in **Fig. 5A,** with the Hex chain inserted manually in VR. The system was simulated for 105 ns of cMD. The procedure for filtering frames fulfilling near-attack conformation criteria was the same as for WT PksD with *O*-acetyl-Ser99.

#### *2.11.2 QM/MM Simulations of PksD Reaction Pathway*

QM/MM calculations were employed to investigate the hydrolysis reaction which takes place in PksD acetylated structure. The representative snapshots were selected from production trajectories based on the distances in oxyanion hole, the distance between Nε2 of His201 and the hydrogen of nucleophilic water, and the distance between oxygen of the nucleophilic water molecule and the carbonyl carbon of the acetyl group of Ser99. QM/MM optimizations were carried out for selected snapshots from the MD trajectories using TINKER7[61] and Gausian16[62] via LICHEM[63]. The QM region was described using the **ω**B97XD functional[64] and 6-31G(d)[65] basis set. All MM atoms were described with the AMBER FF14SB[6] force field. There were 77 QM atoms in total comprising residues His201, Ser98, Ser99, Leu100, Gly101, Gly11, Gln12, Gly13, and a nucleophilic water molecule. The pseudobond approach[66] was employed to treat the covalent boundary between the QM and MM subsystems which includes 5 pesudobonds for residues His201, Ser98, Gly101, Gly11 and Gly13. The optimizations were carried out where all the residues and solvent within a 15 Å sphere of the carbonyl carbon of acetyl group of Ser99 in the QM region, considering as active MM region and the remaining residues and solvent outside this region were kept frozen. The convergence criteria for QM; Root-Mean-Square (RMS) deviation of the atomic positions of QM atoms was 0.001 Å, RMS force was 0.005 Hartree/Bohr, maximum force was 0.015 Hartree/Bohr and RMS deviation of MM atoms was 0.1 Å.[67] The QM/MM long-range electrostatic correction (QM/MM-LREC) with a cutoff of 25 Å was used for the QM subsystem, coupled with sPME for the MM subsystem.[44,68] After the reactant optimisation, the product was built from the optimized reactant and QM/MM optimisation was done for the product to calculate the reaction energy. Optimization of the reaction path was performed with the Quadratic String Method (QSM).[69] Given that a minimum energy path was not obtained (see main text), constrained optimisations of selected structures from the QSM calculation along the reaction path were optimised by constraining selected atoms whilst performing unconstrained QM/MM optimisations on all other atoms. For each constrained structure, no more than three atoms were constrained at a time to obtain the (constrained) optimised structures.

#### *2.11.3 Docking and Molecular Simulation of PksJ ACP4:PksD Complex*

For the simulations of the PksD:PksJ ACP4 complex, the structure of PksJ ACP4 (predicted using Colabfold with Alphafold[11,12] weights) was docked to the crystal structure of PksD in VR whilst accounting for the results of alanine scanning mutagenesis and carbene footprinting as well as complementarity of interactions and distance to the active site. Ppant with acetyl or malonyl substrates were also added in VR by manually feeding them into the substrate channel whilst avoiding steric clashes. The obtained models provided initial coordinates for MD simulations. The AMBER ff19SB force field[49] was used for standard residues and parametrised pantetheine derivatives (Ser+Ppant+acetyl substrate, Ser+Ppant+malonyl substrate).

For the simulation of PksJ ACP4 with acetyl-Ppant in complex with PksD, 50 ns of cMD was followed by ~576 ns of aMD saved every 20 ps (the first 320 ns of the trajectory are shared online). The aMD parameters (estimated from the cMD simulation) were: a) EthreshP = -203183.6 kcal $mol^{-1}$; b) alphaP = 13935.6 kcal $mol^{-1}$; c) EthreshD = 3592.6 kcal $mol^{-1}$; d) alphaD = 278.6 kcal $mol^{-1}$.The frames from aMD simulations were filtered by distance between S99Oγ and Ac carbonyl C1 atom (< 3.5 Å), distance between S99Hγ and H201Nε2 (< 3.5 Å) and the angle between S99Oγ and Ac carbonyl C1 and O atoms (100° < angle < 110°). 20 ns cMD simulations were restarted from frames 3221 and 4230 (reading in only the coordinates and not the velocities), which were close to the geometry required for nucleophilic attack. Initial velocities were disregarded for these simulations. For the simulation of PksJ ACP4 with malonyl-Ppant in complex with PksD, we initially simulated 50 ns of cMD followed by ~288 ns of aMD. The aMD parameters (estimated from the cMD simulations) were: a) EthreshP = -208814.5 kcal $mol^{-1}$; b) alphaP = 14299 kcal $mol^{-1}$; c) EthreshD = 3575.3 kcal $mol^{-1}$; d) alphaD = 278.6 kcal $mol^{-1}$. A 50 ns cMD simulation was restarted from frame 8549 of the aMD simulation (reading in only the coordinates and not the velocities). All the analyses were performed using CPPTRAJ[70] and ChimeraX.[33–35]

The topology files, trajectories and force field parameters for the non-standard residues are shared as open access Zenodo record 7657415 (DOI: 10.5281/zenodo.7657414).

**3. Chemical Synthesis**

**3.1. Synthesis of pantetheine** (**4**).
[(*R*)-2,4-dihydroxy-*N*-(3-((2-mercaptoethyl)amino)-3-oxopropyl)-3,3-dimethylbutanamide]

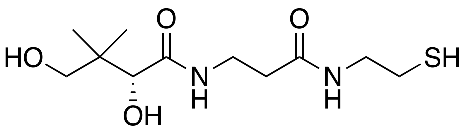

Acetonide protected pantetheine was synthesised according to literature procedures.[71]

To acetonide protected pantetheine (30 mg, 0.094 mmol) was added Dowex 50WX8 resin (30 mg) and methanol (1 mL) and the reaction was stirred overnight. Following filtration and concentration the crude oil was further purified *via* silica gel chromatography (DCM : MeOH = 9 : 1) to yield a colourless oil (9 mg, 0.032 mmol, 34 %).

$\delta_H$ (500 MHz, methanol-d$_4$) 3.89 (s, 1H, C**H**OH), 3.54 – 3.32 (m, 6H, C**H$_2$**OH, CONHC**H**$_2$CH$_2$SH, CHOHCONHC**H$_2$**), 2.59 (t, *J* = 7.0, C**H$_2$**SH), 2.44 (t, *J* = 7.0, CHOHCONHCH$_2$C**H$_2$**), 0.92 (s, 6H, C(C**H$_3$**)$_2$). $\delta_C$ (125 MHz, methanol-d$_4$) 176.1 (**C**ONH(CH$_2$)$_2$SH), 173.9 (CHOH**C**ONH), 77.3 (**C**HOH), 70.3 (**C**H$_2$OH), 43.9 (**C**H$_2$CH$_2$SH), 40.4 (**C**(CH$_3$)$_2$), 36.4 (CHOHCONHCH$_2$**C**H$_2$), 36.3 (CHOHCONH**C**H$_2$), 24.5 (**C**H$_2$SH), 21.3, 20.9 ((**C**H$_3$)$_2$).

HR-MS (*m/z*): $[C_{11}H_{22}N_2O_4SNa]^+$ calculated 301.1198, found 301.1197.

**3.2. Synthesis of acetyl-pantetheine** (**5**).
[(*R*)-*S*-(2-(3-(2,4-dihydroxy-3,3-dimethylbutanamido)propanamido)ethyl) ethanethioate]

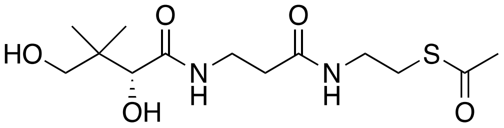

Acetyl-pantetheine (**5**) was synthesised using an adapted literature procedure.[72]

To acetonide protected pantetheine (**4**), (150 mg, 0.440 mmol, 1.0 eq.) and 4-Dimethylaminopyridine (120 mg, 0.982 mmol, 2.2 eq.) in DCM (10 mL) at 0 °C was added acetyl chloride (100 μL, 1.41 mmol, 3.2 eq.). The mixture was then stirred at room temperature overnight. The reaction was quenched *via* addition of 1M HCl. The aqueous layer was extracted twice with DCM. The organics were combined, washed with brine, dried ($MgSO_4$) and concentrated to yield a milky white oil (100 mg, 0.278 mmol, 63.1 %). To the isolated oil (50 mg, 0.157 mmol) was added Dowex 50WX8 resin (50 mg) and methanol (1 mL) and the reaction stirred overnight. Following filtration and concentration the crude oil isolated was further purified *via* silica gel chromatography (DCM : MeOH = 9 : 1) to yield a colourless oil (9 mg, 0.028 mmol, 18 %).

$\delta_H$ (500 MHz, methanol-d$_4$) 3.88 (s, 1H, C**H**OH), 3.53 – 3.32 (m, 6H, C**H$_2$**OH, CONHC**H**$_2$CH$_2$SH, CHOHCONHC**H$_2$**), 2.99 (t, *J* = 6.5, 2H, C**H$_2$**SH), 2.40 (t, *J* = 6.5, 2H, C**H**$_2$CONHCH$_2$), 2.33 (s, 3H, SCOC**H$_3$**), 0.92 (s, 6H, C(C**H$_3$**)**$_2$**). $\delta_C$ (125 MHz, methanol-d$_4$) 197.0 (S**C**O), 176.1 (NH**C**OCHOH), 173.9 CH$_2$**C**ONHCH$_2$), 77.3 (**C**HOH), 70.4 (**C**H$_2$OH), 40.4 (**C**(CH$_3$)$_2$), 40.0 (NH**C**H$_2$CH$_2$SH), 36.4 (**C**H$_2$CONHCH$_2$), 36.3 (CHOHCONH**C**H$_2$), 30.5 (CO**C**H$_3$), 29.4 (**C**H$_2$SH), 21.3, 20.9 (C(**C**H$_3$)**$_2$**).

HR-MS (*m/z*): $[C_{13}H_{24}N_2O_5SNa]^+$ calculated 343.1304, found 343.1308.

### 4. Sequences and Tables

Sequences for PksJ ACP4 and GbnD5 ACP12 have been reported previously.[13,15]

**pHis$_8$-PksD**

```
MGHHHHHHHH SSGLVPRGSH

        10         20         30         40         50         60
MNEPLVFMFS GQGSQYYHMG KELFKENTVF RQSMLEMDAI AARRIGTSIV EEIYHPGKRV

        70         80         90        100        110        120
SDPFDSILFS HPAIFMIEYS LYKVLEDRGI YPDYVLGSSL GEFAAAAVSG VSDAEDMLDC

       130        140        150        160        170        180
ILEQAIIIQN SCDKGKMLAI LDKPQLLNDH PQLFGNSELI SINYDSHFVI SGEEDHIRKI

       190        200        210        220        230        240
MEDLKEKQIL CQLLPVSYAF HSSLIDPAES AYAEFLRSKS FQKPSIPIVS SLTGSCLHVM

       250        260        270        280        290        300
DENFFWNAVR KPMMFREAIR YLESQHTCKF IDLGPSGTLA AFVKQLIPGD SADRCCSIIT

       310        320
PFHQELKNLN TVEYFRTPER KFTR-
```

**MW (+HisTag, -NMet) =** 38,945 Da

**pHis$_6$-BlmVIII ACP7**

```
MGSSHHHHHH SSGLVPRGS

        10         20         30         40         50         60
HMYAAPHTPA QRRIAGWYRD LLGVEHVGLD DDFFALGGDS LLALRLLSQL RDAYGVEISV

        70         80         90
ARMFDEPTVA ALAAATGPPP EETPGQEEVV L
```

**pHis$_6$-IcoA PCP2**

```
MHHHHHHGKP IPNPLLGLDS TENLYFQGID PFT

        10         20         30         40         50         60
DDAFVQSRYE APQGETEQAI AALWSDLLGI ERIGRHDNFF ALGGHSLVAI RMISRIANAT

        70         80         90
GKSLPLRKVF EAPTLAELAL ALADVTSSER AH
```

**Table S1.** Primers, annealing temperatures and restriction sites for cloning and mutagenesis of PksD, BlmVIII ACP7 and IcoA PCP2.

| Construct / Mutation | Forward Primer (5'-3') | Reverse Primer (5'-3') | Temp. (°C) |
|---|---|---|---|
| **pHis$_8$-PksD AH** (XJ-pET28a) | ATA**CATATG**AATGAACCGCTTGT (NdeI) | ATA**GGATCC**TCATCTTGTAAACTTCC (BamHI) | 58 |
| **pET28a-BlmVIII ACP7** (Gibson) | **Backbone:** GAATTCGAGCTCCGTCGACAAGCTTG **Insert:** GTGCCGCGCGGCAGCCATATG TACGCGGCCCCCCACACGCCCGCC | **Backbone:** CATATGGCTGCCGCGCGGCAC **Insert:** CAAGCTTGTCGACGGAGCTCGAATTC TCACAGCACCACCTCTTCCTGGCCGGGC | 50 |
| **pHis$_6$-IcoA PCP2** (pET151) | **CACC**GACGACGCTTTCGTCCA | **TCA**ATGAGCGCGTTCAGACGA | 69 |
| **S98A** | CGTATTGGGA**GCT**AGTCTGGGAG | TAATCAGGATAAATTCCCCTG | 60 |
| **S98H** | CGTATTGGGA**CAC**AGTCTGGGAGAATTC | TAATCAGGATAAATTCCCC | 56 |
| **S99A** | ATTGGGATCA**GCT**CTGGGAGAATTCGC | ACGTAATCAGGATAAATTCC | 58 |
| **Q124A** | CATACTTGAA**GCT**GCTATTATCATCCAGAATTC | CAATCCAGCATATCCTCTG | 56 |
| **F200A*** | GTCCTATGCC**GCT**CATTCTTCGCTTATTGATC | ACAGGAAGCAGCTGGCAA | 56 |
| **H221N** | CTATGCCTTT**AAT**TCTTCGCTTATTGATC | GACACAGGAAGCAGCTGG | 64 |
| **Δ290-324*** | GATTCCGGGT**TGA**TCAGCTGACCG | AGCTGTTTCACAAAAGCAG | 50 |
| **Δ302-324*** | CATAACACCA**TGA**CATCAGGAGCTG | ATTGAGCAGCAACGGTCA | 53 |

*These constructs yielded insoluble protein.

**Table S2.** Recipe for trace element stock (100×) in 1 L*.

| Component | Amount (g) |
|---|---|
| $CaCl_2 \cdot 2H_2O$ | 6.0 |
| $FeSO_4 \cdot 7H_2O$ | 6.0 |
| $MnCl_2 \cdot 4H_2O$ | 1.15 |
| $CoCl_2 \cdot 6H_2O$ | 0.80 |
| $ZnSO_4 \cdot 7H_2O$ | 0.70 |
| $CuCl_2 \cdot 2H_2O$ | 0.30 |
| $H_3BO_3$ | 0.020 |
| $(NH_4)_6Mo_7O_{24} \cdot 4H_2O$ | 0.25 |
| EDTA · 2Na+ · $2H_2O$ | 5.0 |

*The solution was sterile-filtered (0.2 µm cellulose acetate membrane; Sartorius Stedim Biotech) and stored at 4 °C (short-term) or -20 °C (long-term).

**Table S3.** Recipe for vitamin stock (500×) in 300 mL*.

| Component | Amount (g) |
|---|---|
| Choline chloride | 1.0 |
| Folic acid | 1.0 |
| Pantothenic acid | 1.0 |
| Nicotinamide | 1.0 |
| Myo-inositol | 2.0 |
| Pyridoxal hydrochloride | 1.0 |
| Thiamine | 1.0 |
| Riboflavin | 0.10 |
| Disodium adenosine 5'-triphosphate | 0.30 |
| Biotin | 0.20 |

*After placing in $mQH_2O$ with stirring, 10 M NaOH was slowly added until all components were completely dissolved (at a pH of ~12). The clear, dark orange solution was then sterile-filtered (0.2 µm cellulose acetate membrane; Sartorius Stedim Biotech) and stored at 4 °C (short-term) or -20 °C (long-term).

**Table S4.** Data collection, phasing and refinement statistics for PksD.

| | SeMet-derived PksD | Native PksD |
|---|---|---|
| **Data collection** | | |
| X-ray source | DLS, I04-1 | DLS, I03 |
| Temperature (K) | 100 | 100 |
| Wavelength (Å) | 0.91188 | 0.97628 |
| Space group | *C*2 | *C*2 |
| Cell dimensions: | | |
| *a*, *b*, *c* (Å) | 207.71, 83.56, 145.74 | 151.20, 163.33, 186.87 |
| α, β, γ (°) | 90.00, 119.71, 90.00 | 90.00, 104.50, 90.00 |
| Monomers per asym. unit | 4 | 8 |
| Resolution (Å) | 90.20-1.96 (1.98-1.96)* | 90.46-2.20 (2.24-2.20) |
| Nominal resolution (Å)** | 90.20-2.07 | 90.46-2.45 |
| $R_{meas}$ | 0.116 (2.485) | 0.069 (3.166) |
| $\langle I/\sigma(I) \rangle$ | 13.6 (1.2) | 15.2 (0.5) |
| $CC_{1/2}$ | 0.999 (0.675) | 0.999 (0.223) |
| Total no. reflections | 2104939 (98375) | 1524232 (47635) |
| No. unique reflections | 155431 (7639) | 220807 (10069) |
| Completeness (%) | 100.0 (100.0) | 99.5 (91.6) |
| Redundancy | 13.5 (12.9) | 6.9 (4.7) |
| **Refinement** | | |
| Resolution (Å) | 59.79-1.96 (1.98-1.96) | 51.08-2.20 (2.23-2.20) |
| No. unique reflections | 155172 (5110) | 220207 (6326) |
| $R_{work}$ / $R_{free}$ | 0.179/0.203 (0.398/0.442) | 0.198/0.227 (0.469/0.476) |
| No. atoms | 10968 | 20885 |
| Protein | 10295 | 20324 |
| Ligand | 2 | 4 |
| Solvent | 671 | 557 |
| Average *B*-factor (Å$^2$) | 47.9 | 75.4 |
| Protein | 47.4 | 75.4 |
| Ligand | 45.0 | 75.6 |
| Solvent | 54.4 | 77.4 |
| R.m.s. deviations | | |
| Bond lengths (Å) | 0.012 | 0.005 |
| Bond angles (°) | 1.206 | 0.790 |
| Ramachandran | | |
| Favoured (%) | 98.0 | 97.3 |
| Allowed (%) | 2.0 | 2.7 |
| MolProbity | | |
| Clashscore | 4.06 (99$^{th}$ percentile) | 4.81 (98$^{th}$ percentile) |
| MolProbity score | 1.19 (100$^{th}$ percentile) | 1.38 (99$^{th}$ percentile) |
| PDB code | 8AVZ | 8AW0 |

*Values in parentheses refer to the highest-resolution shell. Each dataset was acquired from a single crystal.

**Resolution range for which the mean $I/\sigma(I)$ = 2.0 in the highest-resolution shell. Data were processed to higher resolution limits (see 'Resolution (Å)' row) using paired refinement (see **Section 2.3**).

**Table S5**. Measured masses of PksJ ACP4 mutants in *apo*-, *holo*- and acetylated forms.

| Mutation | *apo*- Calculated Mass (-NMet) / Da | Observed Mass / Da | *holo*- Calculated Mass (-NMet) / Da | Observed Mass / Da | Acetyl Calculated Mass (-NMet) / Da | Observed Mass / Da |
|---|---|---|---|---|---|---|
| **WT** | 15216.92 | 15217.18 | 15557.26 | 15556.37 | 15641.26 | 15640.37 |
| **S12A** | 15200.92 | 15199.28 | 15541.26 | 15540.41 | 15625.26 | 15626.44 |
| **E13A** | 15158.88 | 15158.21 | 15499.22 | 15499.51 | 15583.22 | 15584.87 |
| **Q15A** | 15159.87 | 15158.29 | 15500.21 | 15499.37 | 15584.21 | 15585.41 |
| **S16A** | 15200.92 | 15199.48 | 15541.26 | 15541.18 | 15625.26 | 15625.18 |
| **D20A** | 15172.91 | 15172.34 | 15513.25 | 15513.39 | 15597.25 | 15598.42 |
| **T23A** | 15186.89 | 15186.32 | 15527.23 | 15525.29 | 15611.23 | 15612.32 |
| **E24A** | 15158.88 | 15158.31 | 15499.22 | 15498.29 | 15583.22 | 15584.32 |
| **E25A** | 15158.88 | 15158.34 | 15499.22 | 15498.27 | 15583.22 | 15583.28 |
| **L26A** | 15174.84 | 15173.26 | 15515.18 | 15514.24 | 15599.18 | 15600.27 |
| **R27A** | 15131.81 | 15131.28 | 15472.15 | 15471.22 | 15556.15 | 15557.24 |
| **I28A** | 15174.84 | 15174.29 | 15515.18 | 15514.22 | 15599.18 | 15600.26 |
| **D29A** | 15172.91 | 15172.32 | 15513.25 | 15512.29 | 15597.25 | 15598.32 |
| **R30A** | 15131.81 | 15131.27 | 15472.15 | 15471.19 | 15556.15 | 15557.22 |
| **E31A** | 15158.88 | 15158.31 | 15499.22 | 15497.25 | 15583.22 | 15582.29 |
| **D32A** | 15172.91 | 15172.34 | 15513.25 | 15512.26 | 15597.25 | 15598.30 |
| **F33A** | 15140.82 | 15140.29 | 15481.16 | 15480.22 | 15565.16 | 15566.25 |
| **E34A** | 15158.88 | 15158.29 | 15499.22 | 15497.24 | 15583.22 | 15584.28 |
| **I35A** | 15174.84 | 15174.26 | 15515.18 | 15515.22 | 15599.18 | 15600.24 |
| **D36A** | 15172.91 | 15172.33 | 15513.25 | 15512.32 | 15597.25 | 15598.36 |
| **L38A** | 15174.84 | 15174.28 | 15515.18 | 15514.27 | 15599.18 | 15600.30 |
| **Q40A** | 15159.87 | 15159.30 | 15500.21 | 15499.29 | 15584.21 | 15585.32 |
| **D41A** | 15172.91 | 15172.33 | 15513.25 | 15512.32 | 15597.25 | 15598.36 |
| **Y42A** | 15124.82 | 15124.29 | 15465.16 | 15464.29 | 15549.16 | 15550.32 |
| **V44A** | 15188.86 | 15188.30 | 15529.20 | 15528.28 | 15613.20 | 15613.31 |
| **I47A** | 15174.84 | 15173.27 | 15515.18 | 15514.28 | 15599.18 | 15598.29 |
| **I48A** | 15174.84 | 15174.26 | 15515.18 | 15514.28 | 15599.18 | 15599.30 |
| **Q51A** | 15159.87 | 15159.29 | 15500.21 | 15498.29 | 15584.21 | 15583.31 |
| **L53A** | 15174.84 | 15174.25 | 15515.18 | 15516.27 | 15599.18 | 15600.30 |
| **Q54A** | 15159.87 | 15159.30 | 15500.21 | 15499.26 | 15584.21 | 15585.29 |
| **R55A** | 15131.81 | 15131.23 | 15472.15 | 15470.24 | 15556.15 | 15556.27 |
| **N57A** | 15173.89 | 15173.26 | 15514.23 | 15513.30 | 15598.23 | 15599.34 |
| **R58A** | 15131.81 | 15131.23 | 15472.15 | 15471.24 | 15556.15 | 15556.15 |
| **K59A** | 15159.82 | 15159.25 | 15500.16 | 15499.25 | 15584.16 | 15585.28 |
| **E61A** | 15158.88 | 15158.30 | 15499.22 | 15498.29 | 15583.22 | 15583.32 |
| **D65A** | 15172.91 | 15172.32 | 15513.25 | 15512.32 | 15597.25 | 15598.35 |
| **P66A** | 15190.88 | 15190.30 | 15531.22 | 15530.30 | 15615.22 | 15616.33 |
| **S67A** | 15200.92 | 15200.22 | 15541.26 | 15541.38 | 15625.26 | 15626.98 |
| **I68A** | 15174.84 | 15174.27 | 15515.18 | 15514.26 | 15599.18 | 15599.29 |
| **Y70A** | 15124.82 | 15124.29 | 15465.16 | 15464.36 | 15549.16 | 15550.39 |
| **E71A** | 15158.88 | 15158.22 | 15499.22 | 15498.75 | 15583.22 | 15583.80 |
| **Y72A** | 15124.82 | 15123.29 | 15465.16 | 15464.31 | 15549.16 | 15550.35 |
| **Q76A** | 15159.87 | 15159.28 | 15500.21 | 15499.34 | 15584.21 | 15584.37 |
| **R77A** | 15131.81 | 15130.23 | 15472.15 | 15471.30 | 15556.15 | 15557.34 |
| **D80A** | 15172.91 | 15171.33 | 15513.25 | 15512.40 | 15597.25 | 15597.42 |
| **W81A** | 15101.78 | 15101.20 | 15442.12 | 15441.72 | 15526.12 | 15526.57 |

**Table S6**. Reduction of restraints throughout equilibration using the NVT ensemble with a 1 fs time step.

| Restraints (kcal mol$^{-1}$ Å$^2$) | Time (ns) |
|---|---|
| 100 | 0.02 |
| 50 | 0.02 |
| 10 | 0.02 |
| 1 | 0.02 |
| 0 | 0.1 |

**Table S7**. Statistical analysis of PksD tryptic peptides from carbene footprinting experiments.

| | PksD | | PksD + PksJ ACP4 | | |
|---|---|---|---|---|---|
| **PksD Tryptic Peptides** | **Frac. Mod.** | **Std. Dev.** | **Frac. Mod.** | **Std. Dev.** | ***p* value** |
| [22-25] | 0.3619655 | 0.02162113 | 0.3403422 | 0.02638486 | 0.3339 |
| [26-31] | 0.1709334 | 0.01067518 | 0.1480967 | 0.00285247 | 0.0232 |
| [45-58] | 0.5102484 | 0.0137947 | 0.4789754 | 0.0146829 | 0.0547 |
| [84-88] | 0.1883377 | 0.00505558 | 0.184956 | 0.00850978 | 0.5858 |
| [180-185] | 0.724245 | 0.0570438 | 0.8898502 | 0.03346225 | 0.0123 |
| [261-269] | 0.760896 | 0.01070731 | 0.7808187 | 0.03550909 | 0.4048 |
| [270-284] | 0.4205259 | 0.00174796 | 0.3594469 | 0.0170572 | 0.0035 |
| [285-294] | 0.2458937 | 0.01173018 | 0.2007147 | 0.0105231 | 0.0077 |
| [308-316] | 0.3313128 | 0.01863848 | 0.3047545 | 0.01422335 | 0.1213 |

**Table S8**. Statistical analysis of PksD GluC peptides from carbene footprinting experiments.

| | PksD | | PksD + PksJ ACP4 | | |
|---|---|---|---|---|---|
| **PksD GluC Peptides** | **Frac. Mod.** | **Std. Dev.** | **Frac. Mod.** | **Std. Dev.** | ***p* value** |
| [37-52] | 0.8584059 | 0.04232745 | 0.789951 | 0.07033083 | 0.2221 |
| [79-86] | 0.34071 | 0.02870983 | 0.2897179 | 0.02662175 | 0.0871 |
| [87-102] | 0.3805682 | 0.01070016 | 0.3694584 | 0.01194798 | 0.2964 |
| [103-115] | 0.3846386 | 0.01053699 | 0.4547373 | 0.09863152 | 0.2881 |
| [183-186] | 0.0653324 | 0.00997191 | 0.0688625 | 0.01613445 | 0.7633 |
| [258-263] | 0.245309 | 0.02925405 | 0.2010908 | 0.01739893 | 0.0876 |
| [306-313] | 0.2409599 | 0.00362323 | 0.2165576 | 0.03081721 | 0.2448 |
| [314-319] | 0.2406711 | 0.03366666 | 0.2026361 | 0.02086001 | 0.1716 |
